# Learning the Shape of Evolutionary Landscapes: Geometric Deep Learning Reveals Hidden Structure in Phenotype-to-Fitness Maps

**DOI:** 10.1101/2025.05.07.652616

**Authors:** Manuel Razo-Mejia, Madhav Mani, Dmitri A. Petrov

## Abstract

Elucidating the complex relationships between genotypes, phenotypes, and fitness remains one of the fundamental challenges in evolutionary biology. Part of the difficulty arises from the enormous number of possible genotypes and the lack of understanding of the underlying phenotypic differences driving adaptation. Here, we present a computational method that takes advantage of modern high-throughput fitness measurements to learn a map from high-dimensional fitness profiles to a low-dimensional latent space in a geometry-informed manner. We demonstrate that our approach using a Riemannian Hamiltonian Variational Autoencoder (RHVAE) outperforms traditional linear dimensionality reduction techniques by capturing the nonlinear structure of the phenotype-fitness map. When applied to simulated adaptive dynamics, we show that the learned latent space retains information about the underlying adaptive phenotypic space and accurately reconstructs complex fitness landscapes. We then apply this method to a dataset of high-throughput fitness measurements of *E. coli* under different antibiotic pressures and demonstrate superior predictive power for out-of-sample data compared to linear approaches. Our work provides a data-driven implementation of Fisher’s geometric model of adaptation, transforming it from a theoretical framework into an empirically grounded approach for understanding evolutionary dynamics using modern deep learning methods.

## 1 Introduction

Understanding how genotypes encode phenotypes that allow organisms to survive and reproduce in different environments is one of the central challenges for modern evolutionary biology [1, 2]. Although the field has seen a surge in predictive modeling being deployed in real-world applications ranging from viral dynamics [3] to cancer immunotherapy [4], it is still unclear to what extent evolutionary trajectories under different environmental pressures can be predicted. Multiple challenges make this question difficult to answer. Among these, the vast number of genetic changes available to a population makes it impossible to exhaustively sample and measure the phenotypic consequences of each mutation [5]. Additionally, for most but a few simple model systems, it is unknown what the phenotypic changes driving adaptation are [6].

Nevertheless, advances in modern high-throughput technologies have made it possible to measure the fitness of multiple genotypes, opening up the possibility of extracting valuable information about the genotype-phenotype-fitness map from these datasets [7, 8, 9, 10]. Moreover, an extensive body of empirical studies and theoretical work has suggested that the dynamics of adaptation are highly constrained, making them potentially predictable [11, 12, 5, 13].

Inspired by this body of work and the impressive results that the deep-learning revolution are starting to yield in the life sciences [14], we set out to develop a data-driven framework to uncover the underlying structure of the phenotype-fitness map from high-throughput fitness data. We consider three main ingredients for a computational method to tackle this challenge:

1. The method should be able to take advantage of the data-rich landscape of modern biology, where high-throughput fitness measurements are increasingly common [7, 8, 9, 10].
2. It should be able to uncover the potentially hidden simplicity in the structure of the high-dimensional datasets generated by these experiments by projecting them into a lower-dimensional space in a principled way visualizable.
3. The generated low-dimensional map should be accompanied by the corresponding geometric information on how the high-dimensional space is folded into the map, making the distances on the map meaningful in the high-dimensional space.

Here, we propose the use of a geometry-informed variational autoencoder—a neural network architecture that embeds high-dimensional data into a low-dimensional space while simultaneously learning the geometric transformations that map the learned low-dimensional space back into the high-dimensional space via a metric tensor [15]. To validate the applicability of this method to high-dimensional fitness data, we develop a simple model of adaptive dynamics that captures the main qualitative features of the data generated by high-throughput fitness assays. In this model, multiple lineages perform a biased random walk on a low-dimensional phenotypic space towards regions of higher fitness, defined by the environment-specific topography of the fitness landscape. Of course, the true phenotypic state of a cell is controlled by many interacting variables—membrane permeability, metabolic state, stress responses, and more. Our assumption is not that these axes disappear, but that adaptation proceeds along effective, emergent low-dimensional manifolds that summarize these complex processes [16, 13]. Changing the environment, therefore, amounts to changing the landscape topography while keeping the phenotypic coordinates of the lineages constant. We show that the geometry-informed variational autoencoder (hereafter *RHVAE* for Riemannian Hamiltonian Variational Autoencoder [15]) is able to learn a low-dimensional embedding that retains information about the relative positions of the simulated lineages in the original phenotypic space.

We then apply this method to a dataset of high-throughput fitness measurements of *E. coli* under different antibiotic pressures [9]. Using *IC*_50_ values across eight antibiotics for lineages evolved under tetracycline, kanamycin, and norfloxacin, we demonstrate that the RHVAE learns a low-dimensional embedding that effectively separates different evolutionary trajectories. Our analysis reveals that a two-dimensional RHVAE achieves reconstruction accuracy comparable to a five-dimensional PCA model, while providing superior predictive power for out-of-sample antibiotic responses. Through cross-validation, we show that our nonlinear model more accurately predict resistance profiles than any number of linear components. Taken together, our results suggest that high-dimensional fitness data, such as the one studied here in *E. coli*, can reveal hidden simplicity in evolutionary dynamics when analyzed with the right computational framework, offering new insights into the constraints and predictability of antibiotic resistance evolution.

## 2 Results

### 2.1 Simple Phenotype-Fitness Model

With the advent of high-throughput fitness assays, determining the fitness of multiple genotypes in a plethora of different environments has become a routine procedure in biology [7, 8, 9, 10]. Nevertheless, we argue that we still lack a simple conceptual framework to guide our interpretation of the patterns observed in these datasets. To that end, we propose a simple model of phenotype-fitness dynamics to serve as a guidepost for our interpretation of the data.

Figure 1(A) shows a schematic of the genotype-phenotype-fitness map. A genotype *g* produces a set of measurable traits or phenotypes *p*—depicted as two potential molecular phenotypes, such as an efflux pump rate and the sensitivity of a membrane receptor. Following the tradition in evolutionary biology, we take the most general definition of phenotype, which includes time-varying organism characteristics as a single phenotype. In other words, any genetically-determined organism characteristic—including time-varying features such as developmental time—is considered a single phenotype. The fitness of each genotype is determined by the value of these phenotypes and the environment *E*—depicted as the doubling time of a bacterial population. To formalize this mathematically, we represent genotypes as points in a multi-dimensional phenotypic space, where each dimension is a different trait (*p* ∈ ℝ^*P*^, where *P* is the number of traits). Thus, any genotype *g* has a unique mapping to a phenotypic coordinate. We emphasize again that even dynamically changing organism characteristics are still defined by a single fixed coordinate in phenotype space [17]. However, this mapping is not necessarily bijective, i.e., multiple genotypes may map to the same phenotype. To represent the likelihood of sampling some genotype *g* that maps to some coordinate *p* in phenotype space, we introduce a genotype-to-phenotype density map *GP* that takes a given phenotypic coordinate *p* and returns a number proportional to the probability of sampling a genotype *g* that maps to *p*. This function captures the empirical observation that the genotype-to-phenotype map is highly redundant, with many genotypes mapping to equivalent phenotypes and some phenotypes being extremely rare [5]. Figure 1(B, left) shows a schematic of one possible *GP* map overlaid on a 2D phenotype space. For illustration purposes, we set the *GP* map to contain four negative Gaussian peaks, representing four regions of the phenotype space that are unlikely to be sampled (see Supplementary Material [18] section B for a more detailed discussion). The location and size of these peaks are assumed to be fixed by the genotype-to-phenotype map, irrespective of the environment. Finally, we introduce a fitness function *F* that maps a phenotypic coordinate *p* to a fitness value in a particular environment. We assume the topography of the fitness landscape (the number, size, and position of fitness peaks) is determined by the environment *E*. Figure 1(B, right) shows a schematic of a simple fitness landscape with a single symmetric Gaussian peak.

**Figure 1.**
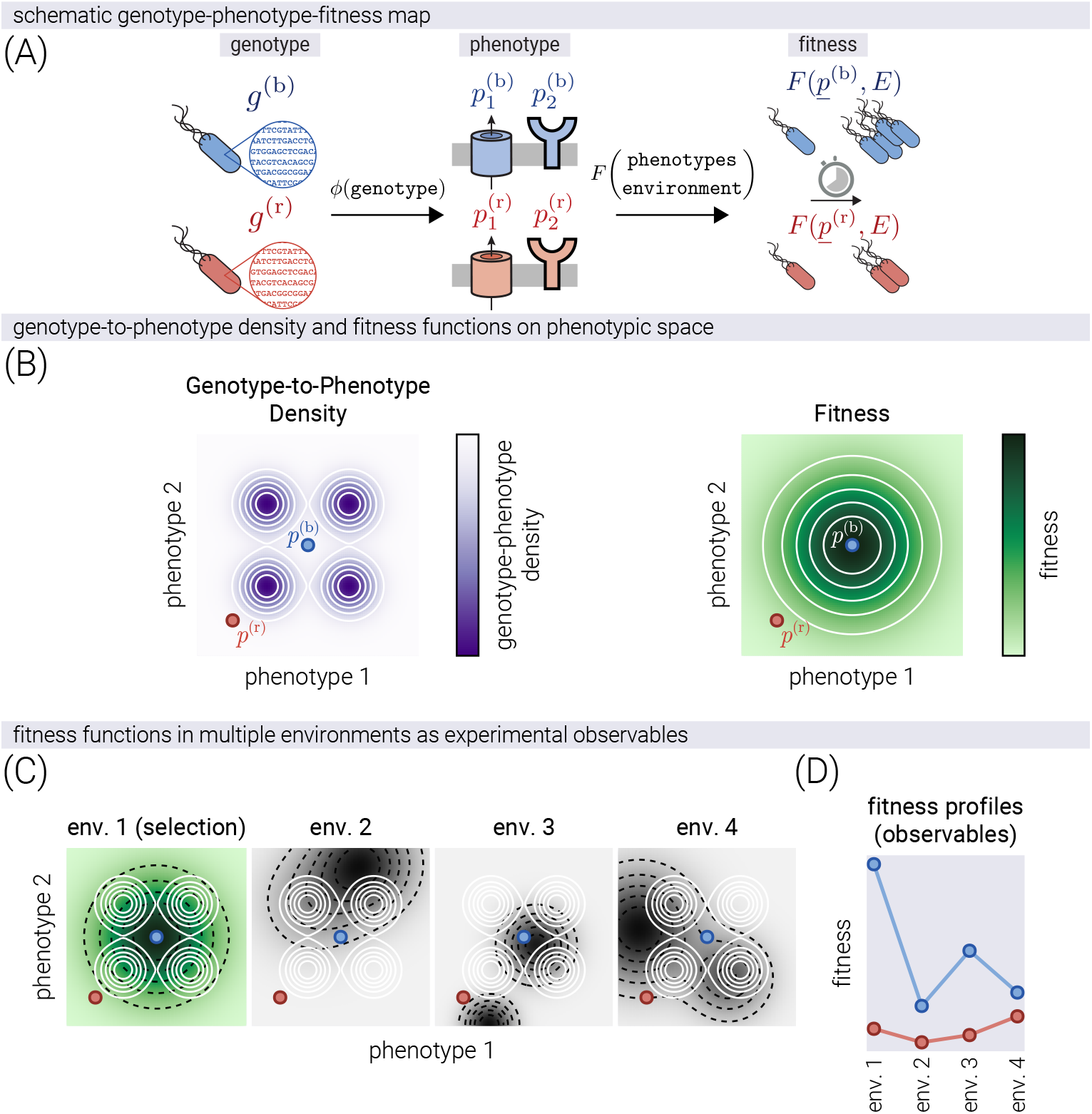
Model of phenotype-fitness dynamics. (A) Schematic of the genotype-phenotype-fitness map. A pair of genotypes *g*^(b)^ and *g*^(r)^ are shown. These genotypes map to two relevant phenotypes *p*_1_ and *p*_2_ via a function *ϕ*. The fitness of each genotype is determined by the value of these phenotypes and their relevance in the environment via a function *F*. (B) Phenotype space functions. Left: The genotype-to-phenotype density map *GP* where the probability of observing a phenotype *p* is shown as a function of the phenotype values *p*_1_ and *p*_2_. White lines show iso-probability contours. Right: The fitness landscape *F* as a function of the phenotype values *p*_1_ and *p*_2_. White lines show iso-fitness contours. The points show the coordinates of the two genotypes in phenotype space. (C) Examples of the effect of changing the environment on the phenotype-fitness landscape. The phenotypic coordinates of any genotype and the genotype-to-phenotype density map are not affected by changes in the environment. The fitness landscape, however, does change as the environment changes. Green landscape shows the evolution condition where selection happens. Gray landscapes show other environments with different topographies (position, number, and height of fitness peaks). Black dashed lines indicate fitness landscape contours, while white solid lines show genotype-phenotype density contours. (D) Fitness profiles. The fitness values of any genotype measured across multiple environments represent an experimental observable.

In recent experimental evolution studies, genotypes are evolved under a fixed environment, but the fitness is measured across multiple environments [7, 9]. To reflect this, Figure 1(C) shows the effect of changing the environment on the phenotype-fitness map. Even though the mapping from genotype to phenotype (coordinates of the example points) and the genotype-to-phenotype density map (white contours) are invariant to the environment, the fitness landscape (black dashed contours) changes as the environment varies. We model this by changing the topography of the fitness map in terms of the number, position, height, and width of the Gaussian peaks. Therefore, we conceptualize the measured fitness profiles on multiple environments as measuring the heights of a fixed phenotypic coordinate on multiple fitness landscapes. Figure 1(D) shows the equivalent experimental observable—the fitness values of any genotype *g* on multiple environments, hereafter referred to as a fitness profile. We ask whether extracting information about the latent phenotypic coordinates is possible based only on such high-dimensional fitness profiles. Conceptually, this is equivalent to asking whether it is possible to reverse the mapping from phenotypes to fitness values and predict the phenotypic coordinates of a genotype based only on its fitness values across multiple environments.

We stress that the latent space inferred by our framework should not be interpreted as a one-to-one reconstruction of the full phenotypic space. Rather, it captures effective structural features of the phenotype–fitness relationship given the data available. The fidelity of this representation will depend on experimental design, data quantity, and dimensionality, all of which we only begin to explore here.

### 2.2 Evolutionary Dynamics on Phenotype Space

Having defined a simple phenotype-fitness model, we now instantiate a set of rules for evolutionary dynamics on this phenotypic space. We implement a simple model that captures two key features of evolutionary processes: the tendency of populations to move towards higher fitness and the constraints imposed by the accessibility of different phenotypes. We frame evolution as a biased random walk where each step is influenced by both the fitness landscape *F* (*p*) and the genotype-phenotype density *GP* (*p*).

At each time step, the population can transition from its current phenotype *p* to a new phenotype *p*′, where the proposed phenotype *p*′ is drawn from a proposal distribution *q*(*p*′ ∣ *p*). Whether this transition occurs—equivalent to the mutation being fixed in the population— depends on the relative fitness of the new phenotype and how easily it can be reached through mutations. Specifically, as derived in the Supplementary Materials [18] section B, the probability of accepting a transition is given by

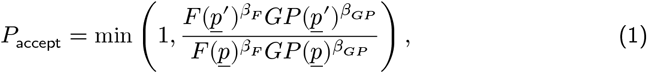

where *β*_*F*_ and *β*_*GP*_ are parameters that separately control the balance between selection and drift. These parameters can be interpreted as the strength of selection and the inaccessibility of certain phenotypes, respectively—higher values of *β*_*F*_ lead to more deterministic trajectories that closely follow fitness gradients, while lower values allow for more stochastic exploration of the phenotype space. In a similar fashion, higher values of *β*_*GP*_ lead to trajectories that avoid low genotype-to-phenotype density regions with higher probability. Detailed mathematical derivations and analysis of this algorithm can be found in the Supplementary Materials [18] section B. For simplicity, in this work we make the simplifying assumption that *β*_*F*_ = *β*_*GP*_ = *β*, as this was sufficient to explore the effects of varying evolutionary stochasticity on the overall landscape.

This formulation represents a simplified model of evolutionary dynamics with specific assumptions: First, transitions to phenotypes with higher fitness are more likely to be fixed, reflecting selection’s influence in our model, though in natural populations even beneficial mutations may be lost due to drift and other factors. Second, transitions to lower fitness phenotypes can occur with some probability, incorporating stochastic elements of evolution. Finally, the *GP* (*p*) term accounts for the mutational accessibility of phenotypes—phenotypes requiring rare genetic variants are less likely to be reached even if highly fit. This approach draws on elements from adaptive dynamics and Metropolis-Hastings algorithms applied to evolutionary processes. Figure 2(A, top) shows example trajectories in phenotypic space of multiple lineages evolving on a simple single-peaked fitness landscape. Figure 2(A, bottom) shows the fitness trajectory of these evolving lineages over time, i.e., the equivalent of the experimental observable we can access in a laboratory setting. When we examine these same phenotypic trajectories under different environmental conditions (Figure 2(B)), we observe that, as expected, while the path through phenotype space remains fixed, the fitness values we measure vary because each environment imposes its unique fitness landscape. The fitness values in all environments for a given lineage at a given time point define one fitness profile, as the ones depicted in Figure 1(D). As shown in the Supplementary Materials [18] section K, our main results using a more detailed population genetics framework yielded consistent conclusions.

**Figure 2.**
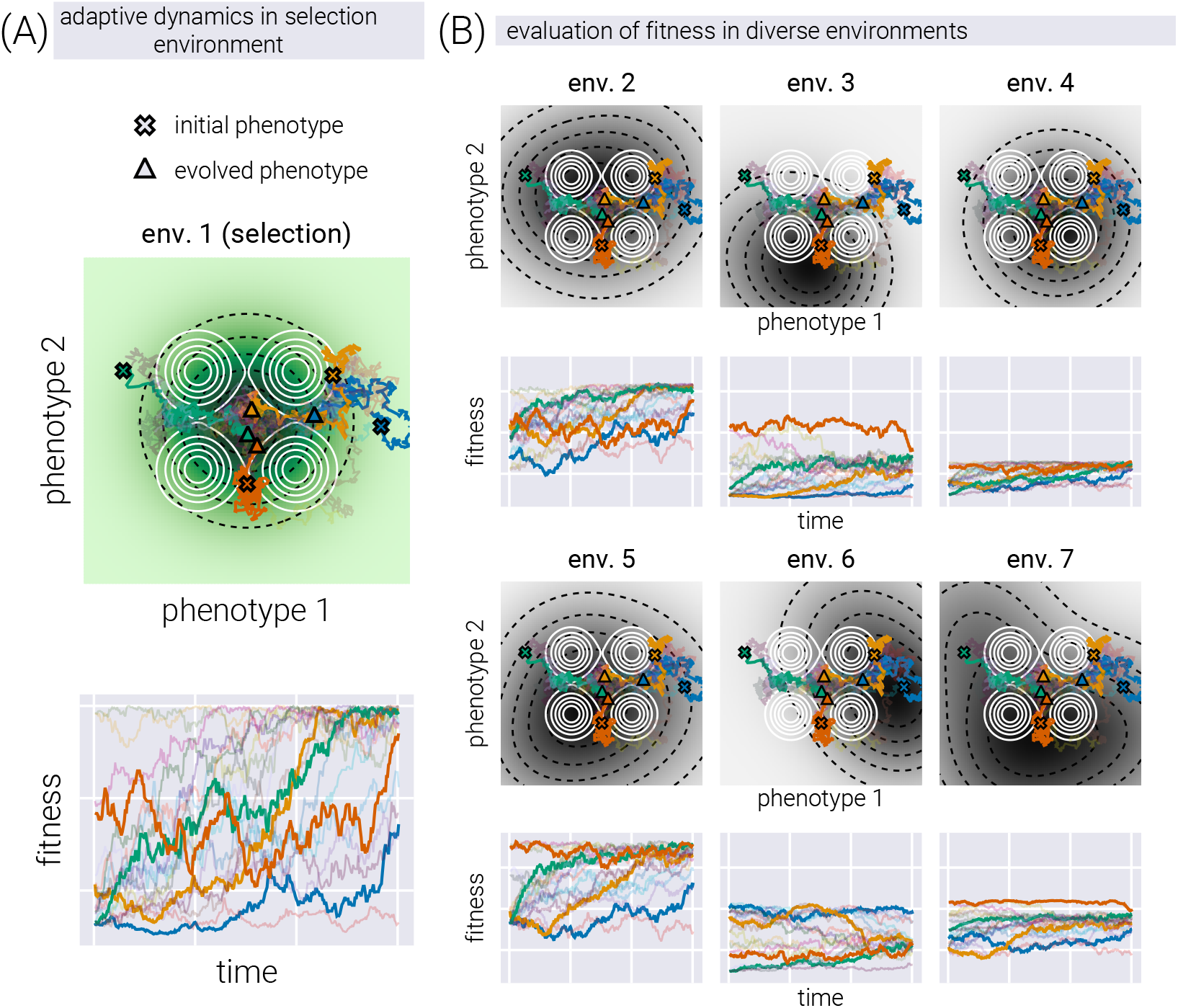
Evolutionary dynamics on phenotype space. (A) Top: Metropolis-like evolutionary dynamics on phenotype space. Each line represents the trajectory of a lineage as it evolves over time, with crosses and triangles denoting the initial phenotypic coordinate of a few selected lineages. Note that trajectories tend to move towards higher fitness values, avoiding low genotype-to-phenotype density regions. Bottom: Fitness over time of the same trajectories shown above. (B) The same trajectories shown in (A) overlaid on different fitness maps determined by different environments. Although the phenotypic coordinates of the genotypes remain the same (top panels), the resulting fitness readouts change as the topography of the environment-dependent fitness landscape changes (bottom panels).

In what follows, we use synthetic data generated from this model, where ten lineages (initial phenotype coordinates) in two replicates (different instantiations of the random dynamics) evolve for 300 time steps on 50 different fitness landscapes (different topographies, meaning number, position, and relative height of fitness peaks). For each of these conditions, the fitness in all other environments is also determined, giving a high-dimensional dataset analogous to the ones in recent experimental studies [7, 9, 10, 19]. We use this dataset to test whether it is possible to reverse the mapping from phenotypes to fitness values and reconstruct the phenotypic coordinates of a genotype based only on fitness profiles.

### 2.3 Data-Driven Dimensionality Reduction via Geometric Variational Autoencoders

Using our simulated fitness profiles that mimic data from modern experimental evolution studies [7, 8, 9, 10, 19], we now develop a data-driven approach to infer phenotypic co-ordinates from fitness measurements across different environments. Figure 3(A) shows a schematic depiction of the idea behind dimensionality reduction applied to fitness profiles. With fitness profiles as vectors on a high-dimensional data space, dimensionality reduction techniques implicitly assume that the positions of points in the data space are highly constrained to occupy only a subset of the available volume. More formally, dimensionality reduction techniques assume that the data points lie on a low-dimensional manifold embedded in the high-dimensional data space. Here, low-dimensionality refers to having the effective number of degrees of freedom required to describe the data being lower than the dimension of the data space. The goal of dimensionality reduction is to find such an embedded manifold that captures the underlying structure of the data. When possible, plotting this embedded manifold as a flat space (either two or three dimensions) allows for the visualization of the underlying data structure. Given the conceptual picture we have developed so far, the hope is that the low-dimensional latent space captures features of the structure of the underlying phenotypic space.

**Figure 3.**
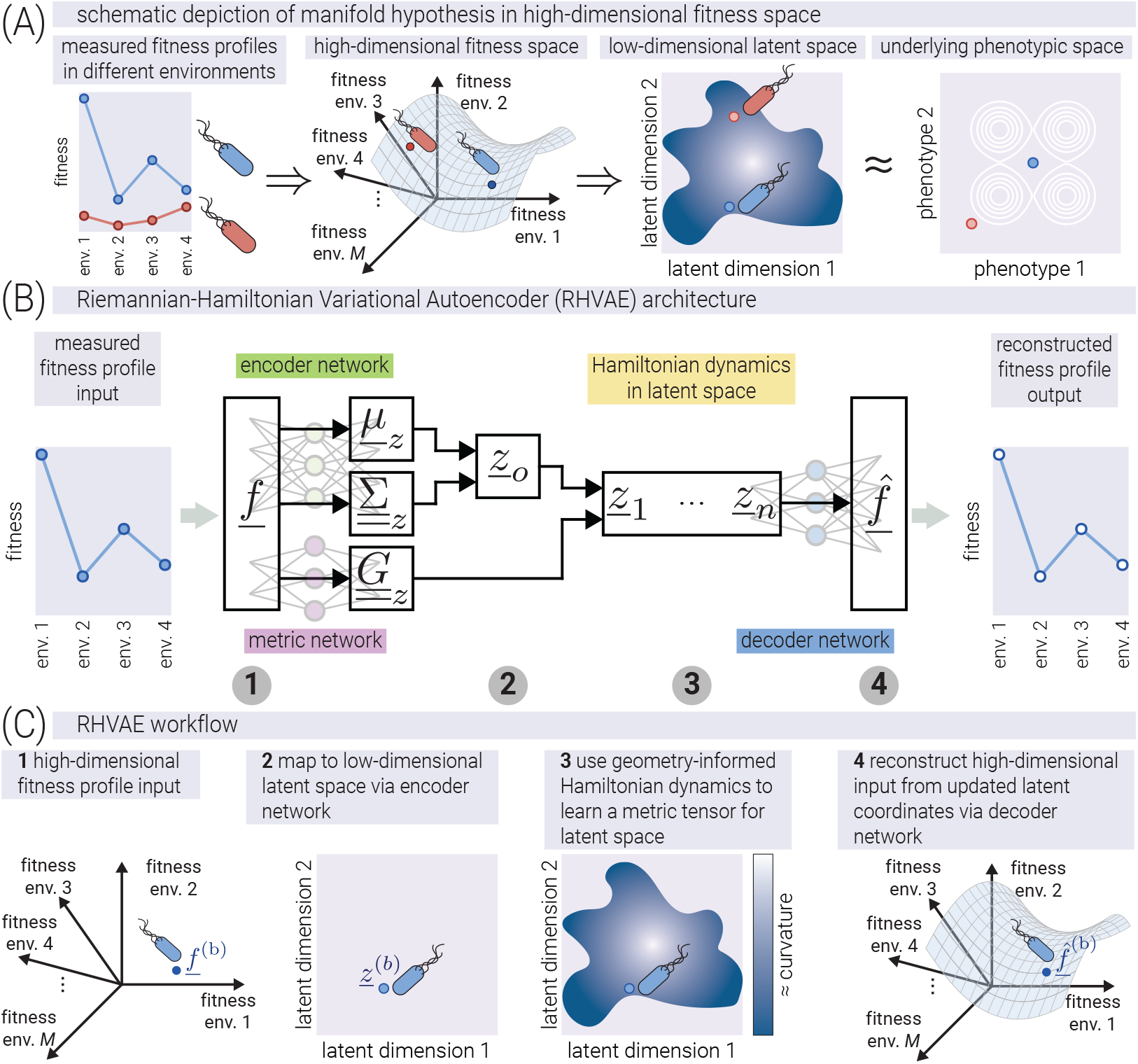
Dimensionality reduction via geometric variational autoencoder. (A) Schematic of dimensionality reduction applied to fitness profiles. High-dimensional fitness vectors are embedded onto a lower-dimensional manifold that captures the underlying structure of the data. (B) Architecture of the Riemannian Hamiltonian VAE (RHVAE). The network consists of three components: an encoder that maps high-dimensional fitness profiles to latent coordinates, a decoder that reconstructs the original fitness profiles, and a metric tensor network that learns the geometric structure of the latent space. (C) RHVAE workflow. The high-dimensional input is mapped into a low-dimensional latent space coordinate. From there, the network learns the metric tensor of the latent space via a specialized network. The latent coordinate is then decoded into an accurate reconstruction of the original fitness profile input.

Recently, the field has focused on linear dimensionality reduction techniques such as singular value decomposition [7] or related methods such as sparse structure discovery [20]. Although computationally efficient, these methods are limited by the rigid assumption that fitness is a linear function of some latent phenotypes. To step away from such strong assumptions, we can take advantage of the advances in deep-learning-based dimensionality reduction techniques, such as the variational autoencoder (VAE) [21]. However, the original VAE framework comes with the limitation that the learned latent representation—built from nonlinear transformations of the input data—does not contain any geometric information about how such nonlinear transformations affect the geometry of the learned space [22]. This is analogous to having a 2D map of the surface of the earth (the latent space), where, to know how distances between two points on the map correspond to distances on the surface of the planet (the data space), we need a position-dependent “scale bar”—more formally, a metric tensor—that allows us to convert distances between representations. This metric tensor is not encoded in the original VAE framework, rendering distances between points in the latent space meaningless as they are not directly related to the distances in the data space.

To address both limitations on the linear and non-geometric nature of popular methods, we propose the use of a recent extension of the VAE framework—the Riemannian Hamiltonian VAE (RHVAE) [15]—which models the latent space as a Riemannian manifold whose metric tensor is also learned from the data, allowing for the computation of meaningful distances between points. Figure 3(B) shows a schematic depiction of the RHVAE architecture. The network consists of three main components: an encoder network, a decoder network, and a specialized network that learns the metric tensor of the latent space. The numbers below the architecture point to the conceptual steps of the RHVAE workflow shown in Figure 3(C). The RHVAE is conceptually similar to the traditional VAE, taking a high-dimensional fitness input *f* as input into an encoder network. This encoder network outputs a sampled latent coordinate *z* [21]. However, before passing this sampled coordinate through a decoder network to reconstruct the fitness profile, the RHVAE uses concepts from Riemannian geometry and Hamiltonian mechanics to jointly train a separate network that learns the metric tensor of the latent space. This metric tensor network, thus, contains all the information needed to map distances in the low-dimensional map to distances in the original data space. We consider this feature of the RHVAE to be of utmost importance if the learned latent space is to be taken seriously as a representation of the underlying phenotypic space. Just as a map without a scale bar is not a useful guide to navigate the terrain, a latent space without a metric tensor is an incomplete representation of the underlying phenotypic space. We direct the reader to the Supplementary Material [18] section C for a mathematically rigorous description of the model.

Next, we turn our attention to applying the RHVAE to the simulated fitness profiles. It is important to highlight that at no point does the RHVAE obtain any knowledge about the identity of the input data. In other words, the RHVAE has no information about the lineage or time point of the fitness profile that it is processing. The network only uses the unidentified fitness profiles to learn the geometric structure of the latent space.

### 2.4 Geometry-Informed Latent Space Captures the Underlying Phenotypic Space

As described in the Supplementary Material [18] section H, we trained three models on the simulated profiles depicted in Figure 2 for comparison: A linear model (via PCA), a vanilla VAE, and an RHVAE. Each model allows us to project the high-dimensional fitness profiles into a low-dimensional latent space. Nevertheless, none of the resulting latent spaces need to be aligned with the ground truth phenotype, as they were constructed with no information about the relative coordinates in phenotype space. To visually compare whether the resulting latent coordinates for the simulated data preserve the general structure of the phenotype space, we aligned each latent space to the ground truth phenotype space using Procrustes analysis (see Supplementary Material [18] section J). Figure 4(A) shows the resulting latent space coordinates colored by the lineage of the simulated profiles. Qualitatively, we observe that, although all models partially capture the relative positions of each lineage with respect to the others, the RHVAE-derived latent space appears to qualitatively preserve the structure of the phenotype space better than the other two models; for example, the four regions of low genotype-to-phenotype density are clearly visible in the RHVAE-derived latent space, while they are not as pronounced in the other two latent spaces. Moreover, when looking at the commonly used PCA-derived latent space, we might be misled into believing that the two highlighted points (diamond markers) are genotypes with very similar phenotypic effects, when, in fact, they are extremely different, as captured by the nonlinear latent spaces. This point is further highlighted in Figure 4(B), where we show the comparison of all pairwise distances between genotypes in the ground truth phenotypic space vs. the different latent spaces. Overall, the RHVAE-derived latent space preserves the pairwise distances between the genotypes better than the other two models.

**Figure 4.**
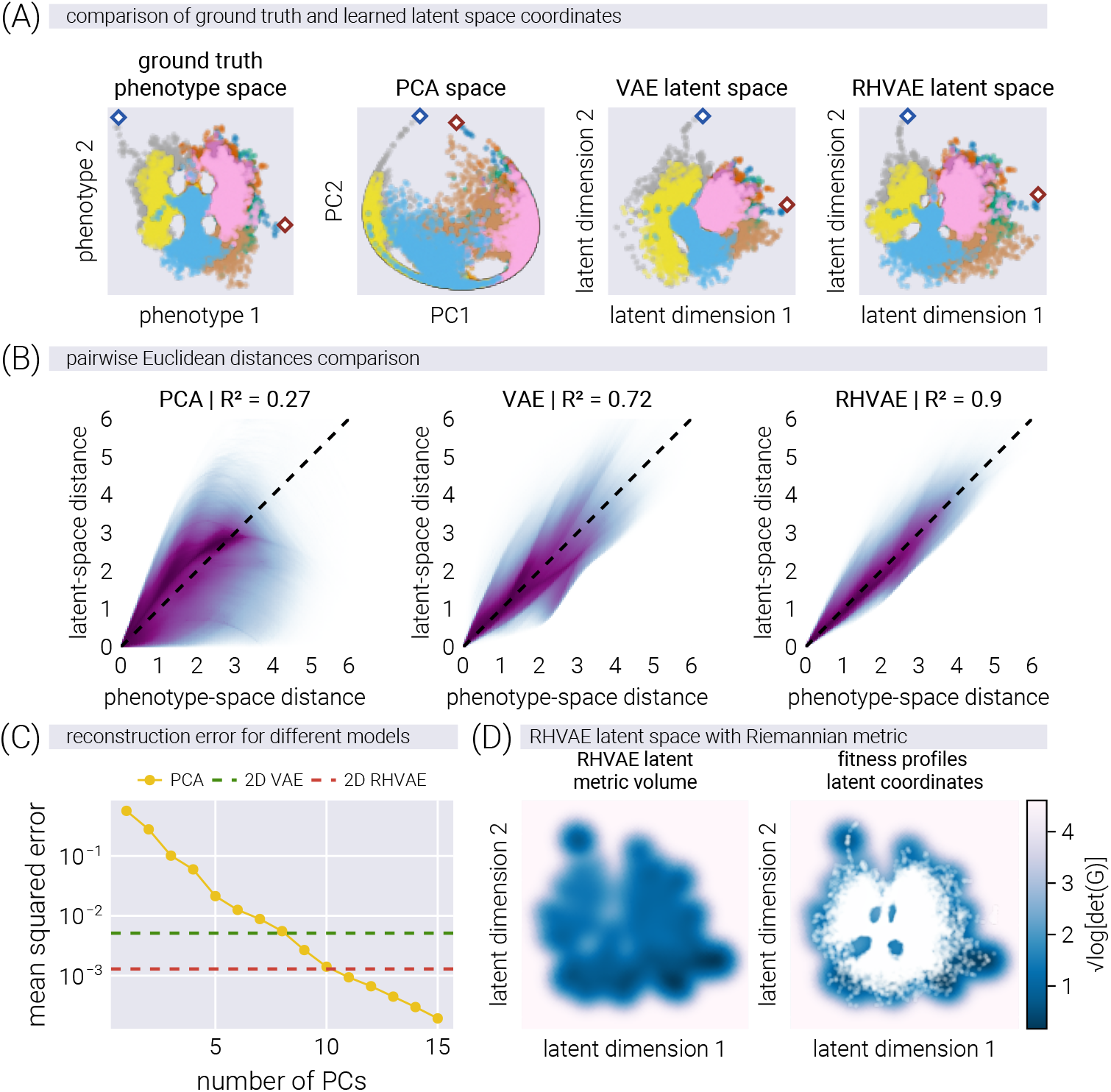
Geometry-informed latent space captures phenotypic structure and improves reconstruction accuracy. (A) Latent space coordinates of simulated fitness profiles colored by lineage, after alignment with the ground truth phenotype space using Procrustes analysis. The RHVAE better preserves phenotypic relationships than PCA or vanilla VAE, as evidenced by the highlighted points (diamond markers). (B) Comparison of all pairwise Euclidean distances between genotypes in the ground truth phenotypic space versus the different latent spaces, showing RHVAE better preserves the original distances. Given the large number of data points and the even larger number of pairwise comparisons, the data are shown as a smear rather than single points. (C) Reconstruction error (MSE) comparison showing 2D-RHVAE achieves accuracy comparable to a 10-dimensional PCA model. (D) Metric volume visualization of the RHVAE latent space (left) and with projected data points (right), revealing regions of high curvature that correspond to areas of low genotype-phenotype density in the simulation.

Having visually confirmed that the RHVAE-derived latent space preserves the structure of the phenotype space better than the other two models, we now investigate whether these latent coordinates also contain enough information to reconstruct the fitness profiles. To this end, we compare the reconstruction accuracy of the RHVAE to that of the PCA and VAE models. Figure 4(C) shows the reconstruction error, defined as

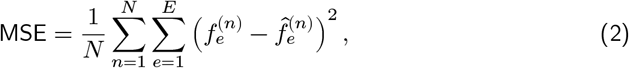

where 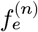and 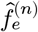are the ground truth and reconstructed fitness values, respectively, for the *n*-th genotype in the *e*-th environment. We observe that the vanilla 2D-VAE model achieves a reconstruction error equivalent to that of an eight-dimensional PCA model. In contrast, the 2D-RHVAE consistently outperforms the other two models, achieving a reconstruction error on par with a ten-dimensional PCA model. This further highlights the ability of the RHVAE to capture the underlying nonlinear structure of the phenotype space while maintaining high reconstruction accuracy.

One of the main advantages of the RHVAE is its ability to learn a metric tensor that captures the local geometry of the latent space. A way to visualize this geometric information is to plot the so-called metric volume, defined as the determinant of the metric tensor at each point in the latent space. Intuitively, the determinant of the metric tensor at a given point can be interpreted as the level of deformation of the latent space at that point. Small values of the metric volume indicate that the latent space is locally flat, while large values suggest significant local curvature. Figure 4(D, left) shows the metric volume of the latent space derived from the RHVAE model. The lighter the color, the larger the metric volume, and thus the larger the local deformation. When overlaying the points over this metric volume (Figure 4(D, right)), we observe that everything outside the data cloud is locally curved. Knowing that neural networks are not able to generalize well outside the data cloud, having this feature is particularly nice because the curvature in the latent space defines the regions where the trained model is able to make accurate predictions. Moreover, we note that regions of high curvature within the data cloud exactly correspond to the regions of low genotype-phenotype density we imposed in the simulation. This suggests that, for a properly sampled phenotypic space, the geometric information learned by the RHVAE is able to capture the non-uniformity of the phenotypic density as variations in the curvature of the latent space, offering a data-driven tool to identify accessible evolutionary paths in phenotype space.

### 2.5 Geometry-Informed Latent Space Reconstruction of Fitness Landscapes

With the ability to capture the underlying structure of the phenotypic space as shown in Figure 4, we can investigate whether the RHVAE can be used to reconstruct the underlying fitness landscapes. For this, we take advantage of the fact that the RHVAE decoder represents a continuous function from latent coordinates to fitness values in different environments. Scanning a grid of latent coordinates allows us to evaluate the fitness landscape at any point in the latent space for all three models shown in Figure 4. Figure 5 shows examples of ground truth fitness landscapes, used to generate the data (first row), and the resulting inferred fitness landscapes for the RHVAE, VAE, and PCA models (second, third, and fourth rows, respectively). Qualitatively, we see that the RHVAE is able to capture the underlying topography of the fitness landscapes, where the number of peaks and their relative positions are similar to the ground truth landscapes. Although the VAE model can also capture the general shape of the fitness landscapes, the lack of a metric in this space— limiting the predictions to the local neighborhood of the training data—makes it difficult to define where to trust the predictions of the model. The PCA model, on the other hand, is not able to capture the underlying structure of the fitness landscapes as there is a nonlinear structure that cannot be captured by a linear transformation.

**Figure 5.**
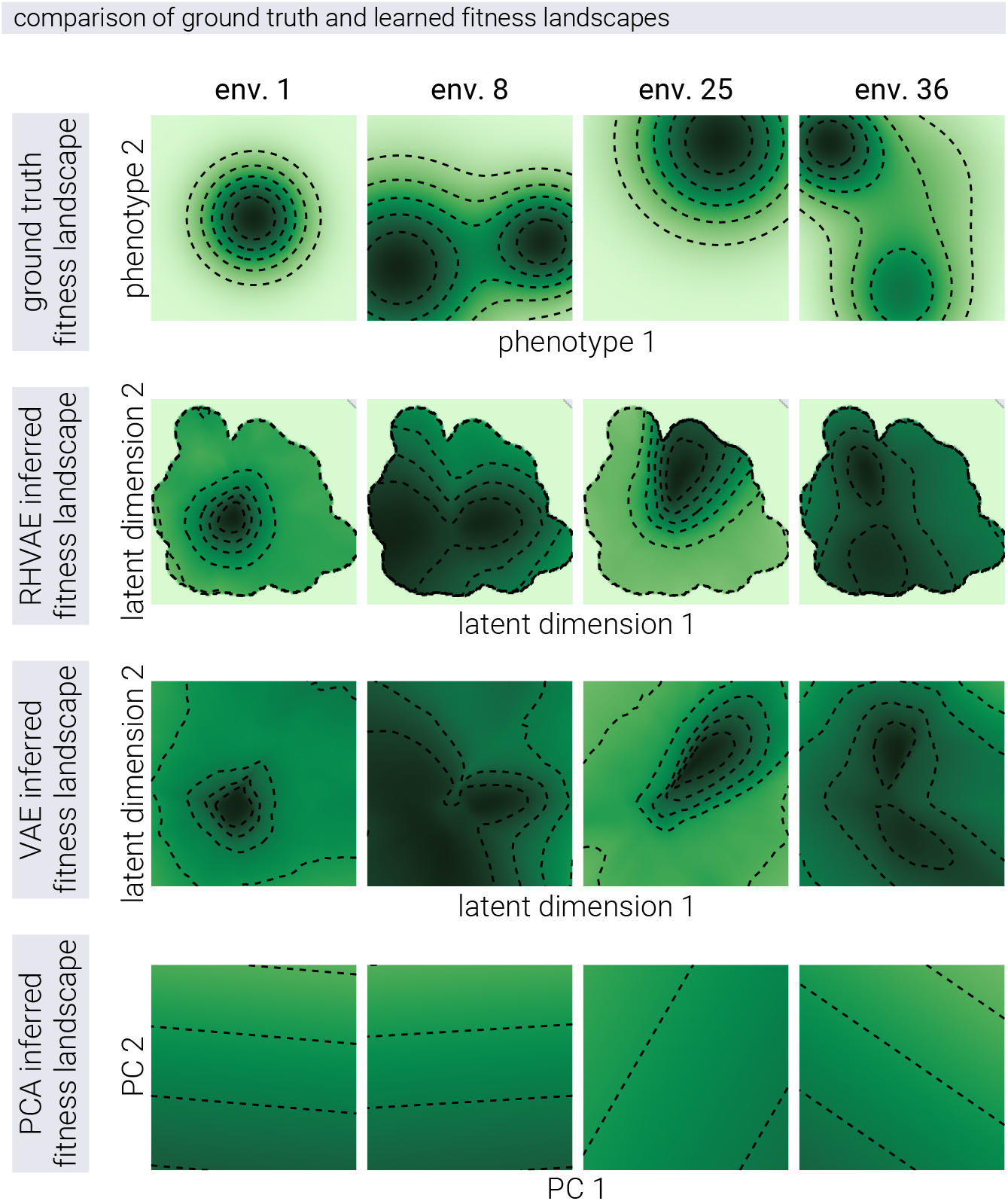
Geometry-informed latent space enables accurate reconstruction of complex fitness landscapes. Comparison of ground truth and reconstructed fitness landscapes across multiple environments. The first row shows examples of the original simulated fitness landscapes used to generate the data. Subsequent rows show reconstructions using the RHVAE (second row), VAE (third row), and PCA (fourth row) models. The RHVAE accurately captures the underlying topography, including the number and relative positions of fitness peaks. While the VAE captures general landscape shapes, it lacks a geometric metric to define prediction reliability regions. PCA fails to represent the nonlinear structure of the landscapes, demonstrating the limitations of linear dimensionality reduction for complex fitness landscape reconstruction.

### 2.6 Evolutionary Dynamics of Antibiotic Resistance in Latent Space

Having shown how the RHVAE can capture the underlying structure of the phenotypic space, we now turn to the application of the model to the analysis of antibiotic adaptation dynamics.

We use the data from Iwasawa et al. [9] where the authors evolved different *E. coli* strains on three different antibiotics: kanamycin (KM), norfloxacin (NFLX), and tetracycline (TET). For this, they used a multi-well plate format where the rows contained a gradient of different antibiotic concentrations, as depicted in Figure 6(A). Every day, a fresh plate with the same arrangement of antibiotics was inoculated with cells from the well in the evolving condition with the highest concentration of antibiotic that still allowed growth. This process was repeated every day for 27 days. Figure 6(B) shows the raw data output from one of the experiments. The optical density is plotted as a function of the antibiotic concentration, obtaining the expected sigmoidal curve. Since this particular example shows data from evolution in the antibiotic being used as the *x*-axis, we can see how, as expected, the sigmoidal curve shifts to the right as the population evolves increasing resistance to the antibiotic over time. From these titration curves, we can use Bayesian inference to extract the value of the antibiotic concentration at which the optical density is 50% of the maximum value. This value—known as the *antibiotic IC*_50_—is a measure of the antibiotic resistance of the evolved strains (see Supplementary Material [18] section A for more details).

**Figure 6.**
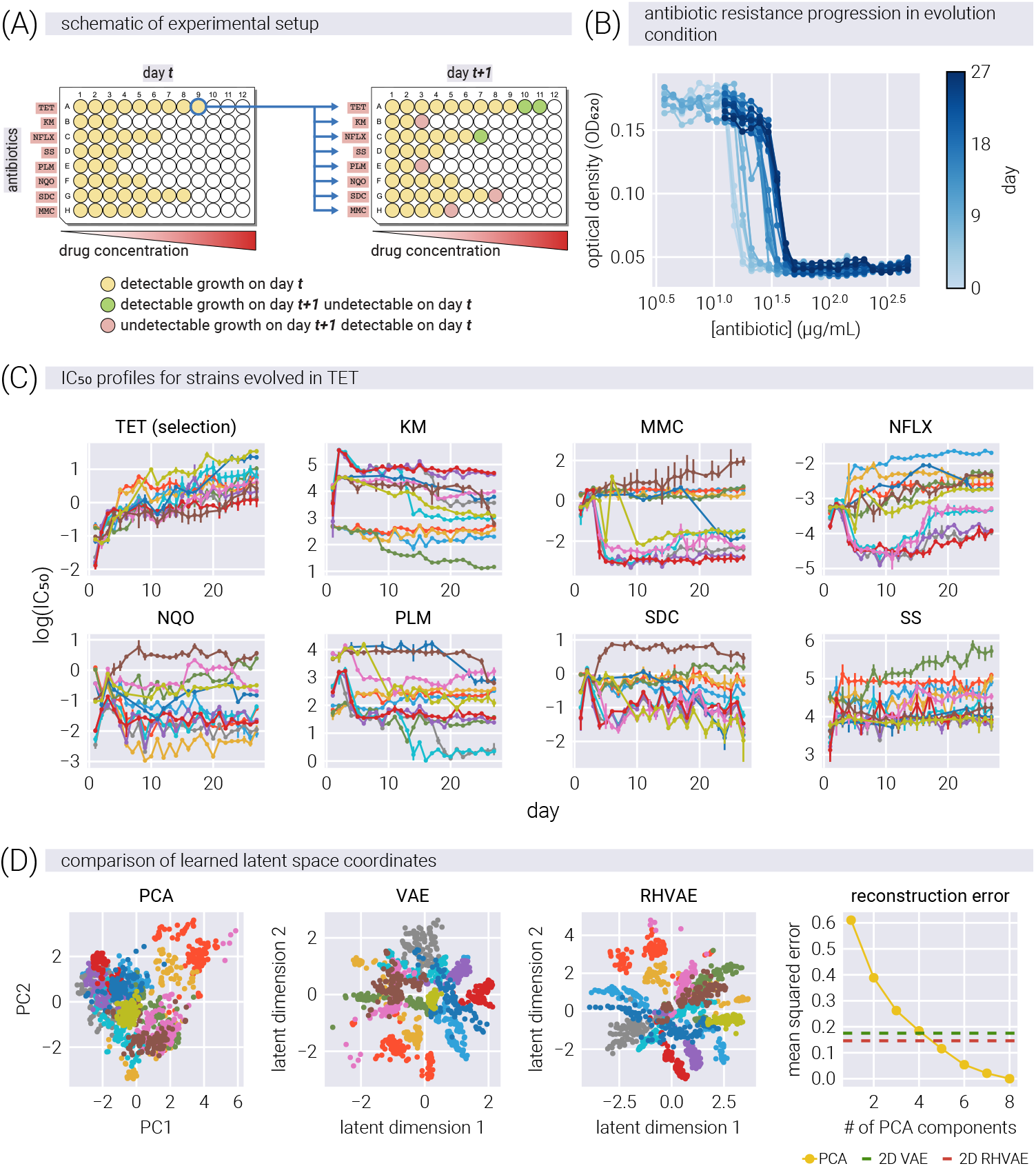
Evolutionary dynamics of antibiotic resistance captured in latent space reveals complex adaptation patterns. (A) Experimental setup using multi-well plates with antibiotic concentration gradients for evolution experiments. (B) Optical density measurements as a function of antibiotic concentration, showing rightward shifts of sigmoidal curves as resistance evolves over time. (C) Time trajectories of *IC*_50_ values across all eight antibiotics (tetracycline (TET), kanamycin (KM), mitomycin C (MMC), norfloxacin (NFLX), 4-nitroquinoline 1-oxide (NQO), phleomycin (PLM), sodium dichromate dihydrate (SDC), and sodium salicylate (SS)) for lineages evolved in tetracycline, demonstrating direct resistance increase in the selection antibiotic and complex collateral effects in other antibiotics. Error bars represent the 95% credible region as obtained via Bayesian inference (see Supplementary Material [18] section A) (D) Latent space projections using PCA, VAE, and RHVAE (first three panels) trained on *IC*_50_ values, with colors representing different evolution conditions, showing improved disentanglement of lineages in nonlinear latent spaces and superior reconstruction accuracy of RHVAE compared to other dimensionality reduction methods. Different colors represent different lineages evolved in different antibiotics. The rightmost panel shows the reconstruction error of the three methods as a function of the number of PCs used for dimensionality reduction.

Figure 6(C) shows the time trajectories of the *IC*_50_ in all eight antibiotics for all cultures evolved in TET. Over time, all lineages increase their *IC*_50_ in TET; however, the trend for all other antibiotics is more complex. Following the same approach as in Figure 4, Figure 6(D) shows the resulting latent spaces for PCA, VAE, and RHVAE trained on the *IC*_50_ values for all eight antibiotics. Each color represents a lineage evolved on a different antibiotic. These projections show that the latent coordinates for different lineages are more disentangled in the nonlinear latent spaces than in PCA. Moreover, the rightmost panel shows that in terms of reconstruction error, RHVAE shows the lowest error—comparable to that of a 5-dimensional PCA.

### 2.7 Non-Linear Latent Space Coordinates are more Predictive of Out of Sample Data

In our simulations, we designed abstract phenotypic coordinates that determined fitness across environments. Applying our proposed method demonstrated that the nonlinear latent space can recover the relative positions in this abstract phenotypic space using fitness data only. While these latent coordinates don’t necessarily represent interpretable biological traits, they effectively capture the underlying structure of the space that determines the genotype fitness in different environments. This success in recovering structural relationships from simulated data encouraged us to test whether the equivalent latent coordinates recovered from experimental data can be used to make more accurate predictions of out-of-sample data.

To test this hypothesis, we adapted the cross-validation approach developed by Kinsler, Geiler-Samerotte, and Petrov [7]. Their original method assesses how linear components predict out-of-sample data by dividing the data matrix into quadrants based on genotypes and antibiotics. We modified this approach for our nonlinear models as shown in Figure 7(A). First, we train our models (VAE and RHVAE) on fitness profiles from all but one antibiotic to generate latent space coordinates. Then, we freeze these coordinates and train only the decoder part to predict responses to the missing antibiotic. This approach allows us to directly compare how well different dimensionality reduction methods capture information needed for prediction. In essence, we use seven antibiotics to place each genotype in latent space, then test how accurately these coordinates predict resistance to the eighth antibiotic. We emphasize that this prediction is not parameter-free: the model must first be calibrated on the available environments. What is novel is that the latent structure inferred without access to the missing antibiotic nevertheless generalizes well, allowing us to reconstruct its response profile post hoc. We direct interested readers to the Supplementary Material [18] section E for a detailed description of the of both approaches.

**Figure 7.**
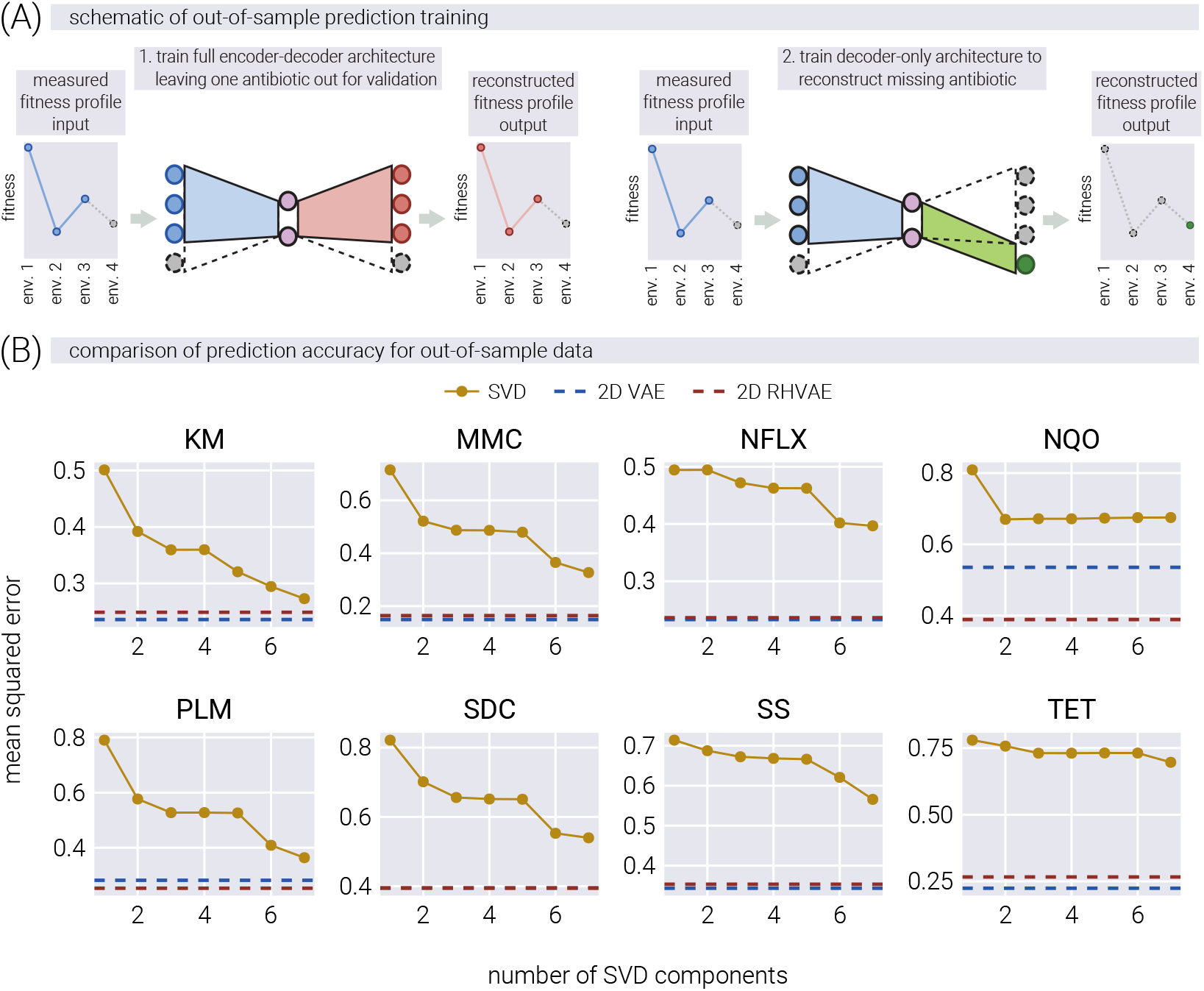
Nonlinear latent space models demonstrate superior predictive power for out-of-sample antibiotic data. (A) Schematic representation of the cross-validation approach used to test predictive power: training a full model on all but one antibiotic to generate latent space coordinates (left), then freezing the encoder parameters and training a decoder-only model on the missing antibiotic data (right). (B) Comparison of predictive accuracy across models, showing reconstruction error for the missing antibiotic as a function of linear dimensions used in SVD bi-cross validation. Horizontal dashed lines represent the accuracy of 2D-VAE (dark blue) and 2D-RHVAE (dark red) models. For all antibiotics, the 2D nonlinear latent space coordinates provide more accurate predictions than any number of linear dimensions, demonstrating that nonlinear latent space representations capture more meaningful biological information about the underlying phenotypic state.

Figure 7(B) compares the predictive power of the three models across different numbers of linear latent space dimensions in terms of reconstruction error. For each panel, the accuracy of the predicted *IC*_50_ value on the missing antibiotic over the validation set is plotted as a function of the number of linear dimensions used in the SVD bi-cross validation approach. The horizontal dashed lines represent the accuracy of the predicted *IC*_50_ value for the 2D-VAE (dark blue) and 2D-RHVAE (dark red) models. For all antibiotics, the 2D nonlinear latent space coordinates are more predictive of the missing antibiotic *IC*_50_ values than any number of possible linear dimensions. This result is consistent with the hypothesis that the nonlinear latent space coordinates capture more information about the phenotypic state of the system under study, allowing for more accurate predictions of out-of-sample data. Furthermore, increasing the dimensionality of the nonlinear latent space from two to three dimensions only marginally improves the predictive power of the model, suggesting that the 2D nonlinear latent space captures most of the information about the phenotypic state of the system (see Supplementary Material [18] section E for a detailed discussion).

## 3 Discussion

Our ability to experimentally track evolving microbial populations presents an unprecedented opportunity to better understand the complexity of the space in which evolutionary dynamics take place. The recent wave of work following the strategy of determining the fitness of multiple genotypes, not in one, but multiple environments [7, 9, 10, 19] suggests that this is a powerful approach to try to uncover the underlying structure of the genotype-phenotype-fitness map. Combined with the body of theoretical work suggesting that the genotype-phenotype map is low-dimensional—meaning that a large number of genotypes result in a small number of distinct phenotypes [16, 5, 13, 23]—there is hope that we can leverage these large-scale datasets to uncover features of the phenotypic space underlying adaptation in a data-driven way. These studies support a view where, although cellular phenotype is intrinsically high-dimensional, adaptation is constrained by a few effective degrees of freedom. This distinction between mechanistic high dimensionality and effective low dimensionality helps reconcile our assumption with the underlying biological complexity.

The main approach taken so far towards this goal has been to use linear decompositions of the fitness data to identify the principal components of phenotypic variation [7, 9, 20, 19]. The simplicity of these linear models comes at the cost of ignoring potentially important nonlinearities in the genotype-phenotype-fitness map. Applying these linear models to data generated from highly nonlinear processes can lead to misleading interpretations of the underlying dimensionality and structure of the phenotypic space, as revealed by embedding theorems from differential geometry (see Supplementary Material [18] section D for a detailed discussion). In this work, we take a step towards removing this limitation by proposing a non-linear dimensionality reduction method that can capture the underlying nonlinear structure of the phenotype-fitness map. We take advantage of recent advances in geometry-informed variational autoencoders [22, 24, 15] to learn a low-dimensional latent space built from high-dimensional fitness measurements. We demonstrate the ability of our method to recover the underlying nonlinear structure of the phenotype-fitness map on synthetic data generated from evolutionary dynamics on a simple phenotypic space with nonlinear mapping to fitness. The geometry-informed latent space not only recovers the relative position of genotypes in the phenotypic space, but also the learned function mapping from latent coordinates to fitness is able to reconstruct the topography of the underlying fitness landscape.

We then apply our method to a dataset of *E. coli* strains evolving in different antibiotics, where a set of cross-resistance values to seven other antibiotics was determined for each lineage at each time point of the adaptation process [9]. When comparing our method to a linear model based on principal components, we find that the different evolved lineages separate much more clearly in the geometry-informed latent space. Furthermore, in terms of reconstruction accuracy, our two-dimensional nonlinear latent space is equivalent to a five-dimensional linear latent space based on principal components. Finally, to show that the information captured by the learned nonlinear latent space is more predictive of the cross-resistance values, we leave one of the eight antibiotics out during the construction of the latent space, and then use these latent coordinates to predict the cross-resistance values of the left-out antibiotic. Our analysis demonstrates that these nonlinear latent coordinates provide superior predictive power compared to linear approaches. Moreover, increasing the dimensionality of the latent space from two to three dimensions only marginally improves the predictive power of the model, suggesting that only two degrees of freedom are needed to capture the changes in the phenotypic state of the evolving lineages (see Supplementary Material [18] section E for a detailed discussion). The results show that the geometry-informed latent space outperforms the linear model based on principal components by a significant margin.

Our approach does not yet allow prediction of responses to completely unseen antibiotics without calibration data. Rather, its contribution lies in demonstrating that nonlinear latent variable models can capture the effective structure of high-dimensional fitness profiles and generalize across held-out environments. This shows that the low-dimensional manifold underlying adaptation is robust enough to be recovered even when parts of the data are missing, a point that was not guaranteed from prior linear approaches.

Our interpretation of latent space as a proxy for phenotypic space comes with limitations. The degree to which this mapping is faithful depends on factors such as data richness, experimental design, and the intrinsic dimensionality of the underlying traits. A full sensitivity analysis will require larger and more varied datasets, which is beyond the scope of the present work. Thus, our results should be viewed as proof of principle rather than a definitive characterization.

Our work can be viewed as a modern, data-driven implementation of Fisher’s geometric model of adaptation [25]. While Fisher conceptualized adaptation as movement in a multidi-mensional phenotypic space, this framework has largely remained theoretical due to the challenges of empirically reconstructing such spaces. By leveraging geometry-informed variational autoencoders, we demonstrate that it is possible to reconstruct the underlying phenotypic manifold directly from fitness measurements across multiple environments. This approach suggests it is possible to transform Fisher’s geometric intuition from a conceptual tool into an empirically grounded framework, where the dimensionality, density, and metric properties of the phenotypic space can be inferred from experimental data rather than assumed a priori. The geometry-informed latent space we recover provides a picture of the adaptive landscape as originally envisioned by Fisher but with the mathematical sophistication afforded by modern deep learning methods.

One unique affordance of latent spaces endowed with a Riemannian metric is that distances in latent space can be related to the resulting distances in fitness space. Furthermore, as suggested by our results on the synthetic data, the curvature of the resulting latent space can be related to the underlying density of genotypes mapping to regions of phenotype space, revealing the accessibility of different regions. Moreover, the metric tensor allows for the identification of the shortest path between two points in latent space. This property begs the question of whether evolutionary trajectories in latent space follow the shortest path between pre- and post-evolved genotypes. Although not fully conclusive, when comparing the shortest path in latent space between the initial and final latent coordinates, some *E. coli* lineages seem to follow the predicted shortest path (See Supplementary Material [18] section F for a detailed discussion). It is tempting to speculate about the biological significance of this observation and whether it is a more general property of the evolutionary process, as it suggests a potential analogous least-action principle of evolution in phenotypic space. Nevertheless, more work is needed to explore this hypothesis further with a larger number of genotypes and environments. A crucial next step will be to integrate genomic data, which would enable a direct comparison between the geodesic paths in our inferred phenotypic space and the actual genetic changes driving adaptation. Such an analysis could reveal whether the geometrically shortest paths correspond to more frequent or probable evolutionary trajectories at the genetic level, providing a powerful, data-driven link between the geometry of adaptation and its underlying molecular mechanisms.

Moreover, it remains an open question as to what is the biological meaning of the learned latent coordinates. Complementing these fitness measurements with high-dimensional phenotypic characterizations—such as RNA-sequencing data—could provide additional insights into the biological nature of the latent dimensions. For example, following a similar approach to the cross-validation analysis presented in the main text, where the latent space was learned using only fitness information, one could then fix the encoder (effectively fixing the latent coordinates based only on fitness profiles) and then train a decoder that maps the latent coordinates to the high-dimensional phenotypic data. This approach could provide information to constrain the biological functions enriched on the latent dimensions, effectively determining the name of the axes in the latent space. We see this as an exciting direction for future work, where the combination of high-dimensional fitness and phenotypic data, guided by theoretical understanding of evolutionary dynamics, could shed light on the nature of biological adaptation.

In our study, we employed two variants of variational autoencoders: one incorporating a Riemannian metric and one without this geometric constraint. While the geometry-informed model demonstrated marginally enhanced performance in capturing the underlying structure of the fitness data, it incurs a significant computational cost during training. Nevertheless, we consider the geometric information about the latent space essential for meaningful interpretation of the distances between evolved lineages. Just as a geographical map without a scale bar provides incomplete spatial information, a latent representation without metric properties offers limited insight into the relationships between data points. If these learned latent spaces are to be considered faithful representations of biological reality rather than mere computational abstractions, the geometric structure becomes critically important. We acknowledge recent proposals by Chadebec and Allassonnière [26] that aim to approximate the metric tensor of standard variational autoencoders post-training. Investigating the efficacy of these more computationally efficient alternatives compared to explicitly geometry-informed models represents an important direction for future research.

### Scope and Hierarchy of Claims

To provide clarity on the nature of our findings and address the limitations inherent in this framework, we categorize our claims into three levels of certainty:

- **Key Results (Well-Supported by Evidence):** These include the demonstrated improvements in reconstruction accuracy and out-of-sample prediction using RHVAE, the clear separation of evolved lineages in latent space compared to linear methods, and the successful recovery of phenotypic structure in synthetic validation.
- **Plausible Interpretations (Supported by Theory):** We interpret the learned latent space as a proxy for an effective phenotypic space. This is consistent with a large body of theoretical work suggesting low-dimensional genotype-phenotype maps, though a definitive proof for any specific biological system requires further data.
- **Speculative Directions (Future Investigation):** The potential connection between geodesic paths in latent space and actual evolutionary trajectories, while intriguing, remains speculative. Similarly, the integration with genomic data and scaling to much larger environmental datasets represent future directions rather than established conclusions of this work.

## Data and code availability

All data and custom scripts were collected and stored using Git version control. Code for raw data processing, theoretical analysis, and figure generation is available on the GitHub repository [27].

## Acknowledgements

We would like to thank David Larios, Enrique Amaya, and especially Griffin Chure for their helpful advice and discussion. We are also grateful to Jose Aguilar-Rodriguez, Stefan Bassler, Benjamin Good, Shreyas Gopalakrishnan, Olivia Ghosh, Shaili Mathur, and Anastasia Lyulina for their helpful feedback on the manuscript. We are extremely grateful to Junichiro Iwasawa for sharing the raw data from his study; his openness and willingness to share data and the immediate response are examples of the best practices in open science.

This work was supported by several grants: the NIH/NIGMS, Genomics of rapid adaptation in the lab and in the wild R35GM11816506 (MIRA grant) to DAP; grants from the NSF (DMS-2235451) and Simons Foundation (MPTMPS-00005320) to the NSF-Simons National Institute for Theory and Mathematics in Biology (NITMB); the Chan Zuckerberg Initiative DAF (DAF2023-329587), an advised fund of the Silicon Valley Community Foundation; and grants from the NSF (PHY-1748958) and the Gordon and Betty Moore Foundation (2919.02) to the Kavli Institute for Theoretical Physics (M.M.). MRM was partially supported by the Schmidt Science Fellowship. M.M. was supported by the NSF-Simons Center for Quantitative Biology at Northwestern University (DMS-1764421) and the Simons Foundation (597491-RWC). M.M. is a Simons Investigator. DP is a CZ Biohub investigator.

