## Supplementary Material for "Learning the Shape of Evolutionary Landscapes: Geometric Deep Learning Reveals Hidden Structure in Phenotype-to-Fitness Maps"

### Supplementary Materials

#### A. Bayesian inference of $IC_{50}$ resistance values

The experimental data used throughout this work was kindly provided by the first author of the excellent paper [1]. In there, the authors evolved *E. coli* strains on different antibiotics via an automated liquid handling platform. Here, we will detail our analysis of the raw data to infer the $IC_{50}$ values of the strains on a panel of antibiotics.

##### A.1. Experimental design

To justify the analysis, we first need to understand the experimental design. To determine the antibiotic resistance on multiple antibiotics, the authors adopted a 96-well plate design depicted in Figure A. On a 96-well plate, each row contained a titration of one of eight antibiotics. This format allowed measuring the optical density of the bacterial culture in each well as a function of each of the antibiotic concentrations. To evolve the strains on a particular antibiotic (e.g., tetracycline as in Figure A), the authors propagated the strains every day by inoculating a fresh plate with the same panel of antibiotic titrations solely from the well with the highest tetracycline concentration in which there was still measurable growth the day before. In this way, the authors selected for the strains with the highest antibiotic resistance while still measuring the effects of this adaptation on the other antibiotics.

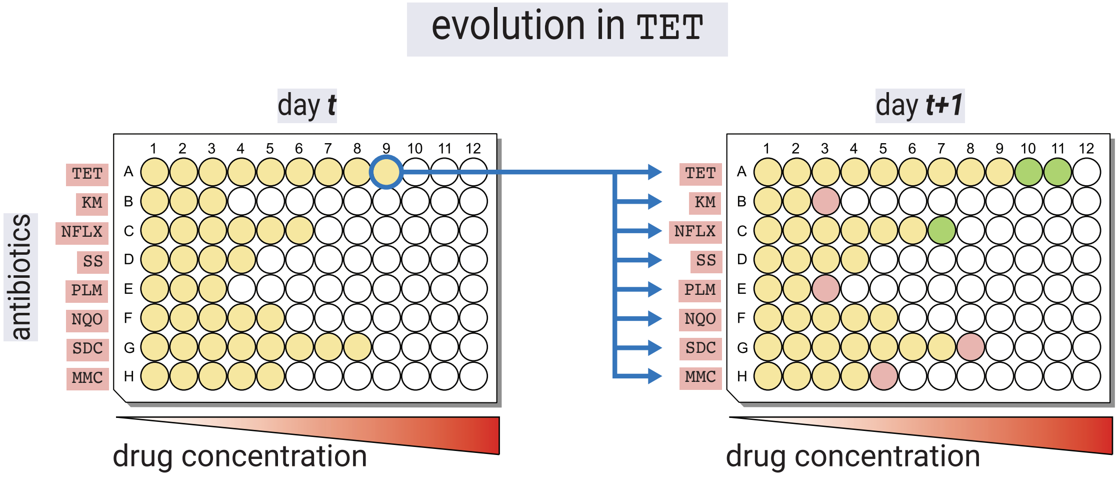

**Figure A. Iwasawa et al. experimental design.** Schematic of the experimental design used by Iwasawa et al. [1]. On a 96-well plate, each of the eight rows contained a titration of one of eight antibiotics. For strains evolved in tetracycline (TET), the plate for each day $t + 1$ was inoculated by propagating from the well with the highest tetracycline concentration in which there was still measurable growth on day $t$ to all 96 wells.

Figure B shows a typical progression of the optical density curves over the course of the experimental evolution. When adapted to a particular antibiotic, the optical density curves with their sigmoidal shape shifted to the right, indicating that the strains were able to grow at higher concentrations of the antibiotic.

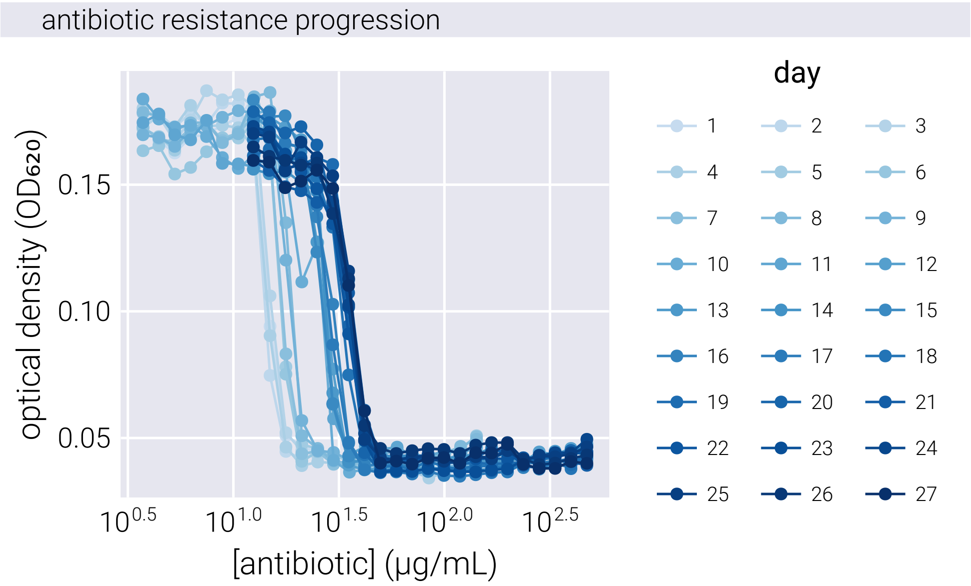

**Figure B. Iwasawa et al. typical experimental data.** The optical density curves of the bacterial culture in the evolution antibiotic over the course of the experiment. The curves are shifted to the right, indicating that the strains were able to grow at higher concentrations of the antibiotic.

#### A.2. Bayesian inference of $IC_{50}$ values

Given the natural sigmoidal shape of the optical density curves, we can model the optical density $OD(x)$ of the bacterial culture as a function of the antibiotic concentration $x$. Following the approach of [1], we model the optical density as a function of the antibiotic concentration $x$ as

$$f(x) = \frac{a}{1 + \exp\left[b\left(\log_2 x - \log_2 \mathrm{IC}_{50}\right)\right]} + c \tag{S1}$$

where $a$, $b$, and $c$ are nuisance parameters of the model, $\mathrm{IC}_{50}$ is the parameter of interest, and $x$ is the antibiotic concentration. We can define a function to compute this model.

Given the model presented in Equation S1, and the data, our objective is to infer the value of all parameters (including the nuisance parameters). By Bayes theorem, we write

$$\pi(\mathrm{IC}_{50}, a, b, c \mid \text{data}) = \frac{\pi(\text{data} \mid \mathrm{IC}_{50}, a, b, c)\pi(\mathrm{IC}_{50}, a, b, c)}{\pi(\text{data})}, \tag{S2}$$

where data consists of the pairs of antibiotic concentration and optical density. Let's begin by defining the likelihood function. For simplicity, we assume each datum is independent and identically distributed (i.i.d.) and write

$$\pi(\text{data} \mid \mathrm{IC}_{50}, a, b, c) = \prod_{i=1}^{n} \pi(d_i \mid \mathrm{IC}_{50}, a, b, c), \tag{S3}$$

where $d_i = (x_i, y_i)$ is the $i$-th pair of antibiotic concentration and optical density, respectively, and $n$ is the total number of data points. We assume that our experimental measurements can be expressed as

$$y_i = f(x_i, \mathrm{IC}_{50}, a, b, c) + \epsilon_i, \tag{S4}$$

where $\epsilon_i$ is the experimental error. Furthermore, we assume that the experimental error is normally distributed, i.e.,

$$\epsilon_i \sim \mathcal{N}(0, \sigma^2), \tag{S5}$$

where $\sigma^2$ is an unknown variance parameter that must be included in our inference. Notice that we assume the same variance parameter for all data points since $\sigma^2$ is not indexed by $i$.

Given this likelihood function, we must update our inference on the parameters as

$$\pi(\mathrm{IC}_{50}, a, b, c, \sigma^2 \mid \text{data}) = \frac{\pi(\text{data} \mid \mathrm{IC}_{50}, a, b, c, \sigma^2)\pi(\mathrm{IC}_{50}, a, b, c, \sigma^2)}{\pi(\text{data})}, \tag{S6}$$

to include the new parameter $\sigma^2$. Our likelihood function is then of the form

$$y_i \mid \mathrm{IC}_{50}, a, b, c, \sigma^2 \sim \mathcal{N}(f(x_i, \mathrm{IC}_{50}, a, b, c), \sigma^2). \tag{S7}$$

For the prior, we assume that all parameters are independent and write

$$\pi(\mathrm{IC}_{50}, a, b, c, \sigma^2) = \pi(\mathrm{IC}_{50})\pi(a)\pi(b)\pi(c)\pi(\sigma^2). \tag{S8}$$

Let's detail each prior.

1. $\mathrm{IC}_{50}$: The $IC_{50}$ is a strictly positive parameter. However, we will fit for $\log_2(\mathrm{IC}_{50})$. Thus, we will use a normal prior for $\log_2(\mathrm{IC}_{50})$. This means we have

$$\log_2(\mathrm{IC}_{50}) \sim \mathcal{N}(\mu_{\log_2(\mathrm{IC}_{50})}, \sigma^2_{\log_2(\mathrm{IC}_{50})}). \tag{S9}$$

2. $a$: This nuisance parameter scales the logistic function. Again, the natural scale for this parameter is a strictly positive real number. Thus, we will use a lognormal prior for $a$. This means we have

$$a \sim \text{LogNormal}(\mu_a, \sigma_a^2). \tag{S10}$$

3. $b$: This parameter controls the steepness of the logistic function. Again, the natural scale for this parameter is a strictly positive real number. Thus, we will use a lognormal prior for $b$. This means we have

$$b \sim \text{LogNormal}(\mu_b, \sigma_b^2). \tag{S11}$$

4. $c$: This parameter controls the minimum value of the logistic function. Since this is a strictly positive real number that does not necessarily scale with the data, we will use a half-normal prior for $c$. This means we have

$$c \sim \text{Half-}\mathcal{N}(0, \sigma_c^2). \tag{S12}$$

5. $\sigma^2$: This parameter controls the variance of the experimental error. Since this is a strictly positive real number that does not necessarily scale with the data, we will use a half-normal prior for $\sigma^2$. This means we have

$$\sigma^2 \sim \text{Half-}\mathcal{N}(0, \sigma_{\sigma^2}^2). \tag{S13}$$

##### A.2.1. Residual-based outlier detection

To identify and remove outliers from our optical density measurements, we implemented a residual-based outlier detection method. First, we fit the logistic model (Equation S1) to the data using a deterministic least-squares approach. We then computed the residuals between the fitted model and the experimental measurements. Points with residuals exceeding a threshold of two standard deviations from the mean residual were classified as outliers and removed from subsequent analysis. This approach helped eliminate experimental artifacts while preserving the underlying sigmoidal relationship between antibiotic concentration and optical density.

#### A.3. MCMC sampling of the posterior

With this setup, we can sample the posterior distribution using Markov Chain Monte Carlo (MCMC) methods. In particular, we used the `Turing.jl` package [2] implementation of the No-U-Turn Sampler (NUTS). All code is available in the GitHub repository for this work. Figure C shows the posterior predictive checks of the posterior distribution for the $IC_{50}$ parameter. Briefly, the posterior predictive checks are a visual method to assess the fit of the model to the data. The idea is to generate samples from the posterior distribution and compare them to the data. If the model is a good fit, the samples should be able to reproduce the data.

The effectiveness of the residual-based outlier detection method is demonstrated in Figure C, where the posterior predictive checks show significantly tighter credible intervals after outlier removal, particularly in regions of high antibiotic concentration where experimental noise tends to be more pronounced.

#### B. Metropolis-Hastings Evolutionary Dynamics

In the main text, we describe a simple algorithm for evolving a population as a biased random walk on a phenotypic space controlled by both a fitness function and a genotype-phenotype density function. Here, we provide a more detailed description of the algorithm and its implementation.

##### B.1. Landscape Description

We assume that phenotypes driving adaptation can be described by real-valued numbers. Let $\underline{x}(t) = (x_1(t), x_2(t), \cdots, x_N(t))$ be an $N$ dimensional vector describing the phenotype of a population at time $t$. The fitness for a given environment $E$ is a scalar function $F_E(\underline{x})$ such that

$$F_E : \mathbb{R}^N \rightarrow \mathbb{R}, \tag{S14}$$

i.e, the coordinates in phenotype space map to a fitness value. Throughout this work, we consider a series of Gaussian fitness peaks, i.e.,

$$F_E(\underline{x}) = \sum_{p=1}^{P} A_p \exp\left(-\frac{1}{2}(\underline{x} - \underline{\hat{x}}^{(p)})^T \Sigma_p^{-1} (\underline{x} - \underline{\hat{x}}^{(p)})\right), \tag{S15}$$

where $P$ is the number of peaks, $A_p$ is the amplitude of the $p$-th peak, $\underline{\hat{x}}^{(p)}$ is the $N$-dimensional vector of optimal phenotypes for the $p$-th peak, and $\Sigma_p$ is the $N \times N$ covariance matrix for the $p$-th peak. This formulation allows for multiple peaks in the fitness landscape, each potentially having different heights, widths, and covariance structures between dimensions. The covariance matrix $\Sigma_p$ captures potential interactions between different phenotypic dimensions, allowing for more complex and realistic fitness landscapes.

In the same vein, we define a genotype-phenotype density function $GP(\underline{x})$ as a scalar function that maps a phenotype to a mutational effect, i.e.,

$$GP : \mathbb{R}^N \rightarrow \mathbb{R}, \tag{S16}$$

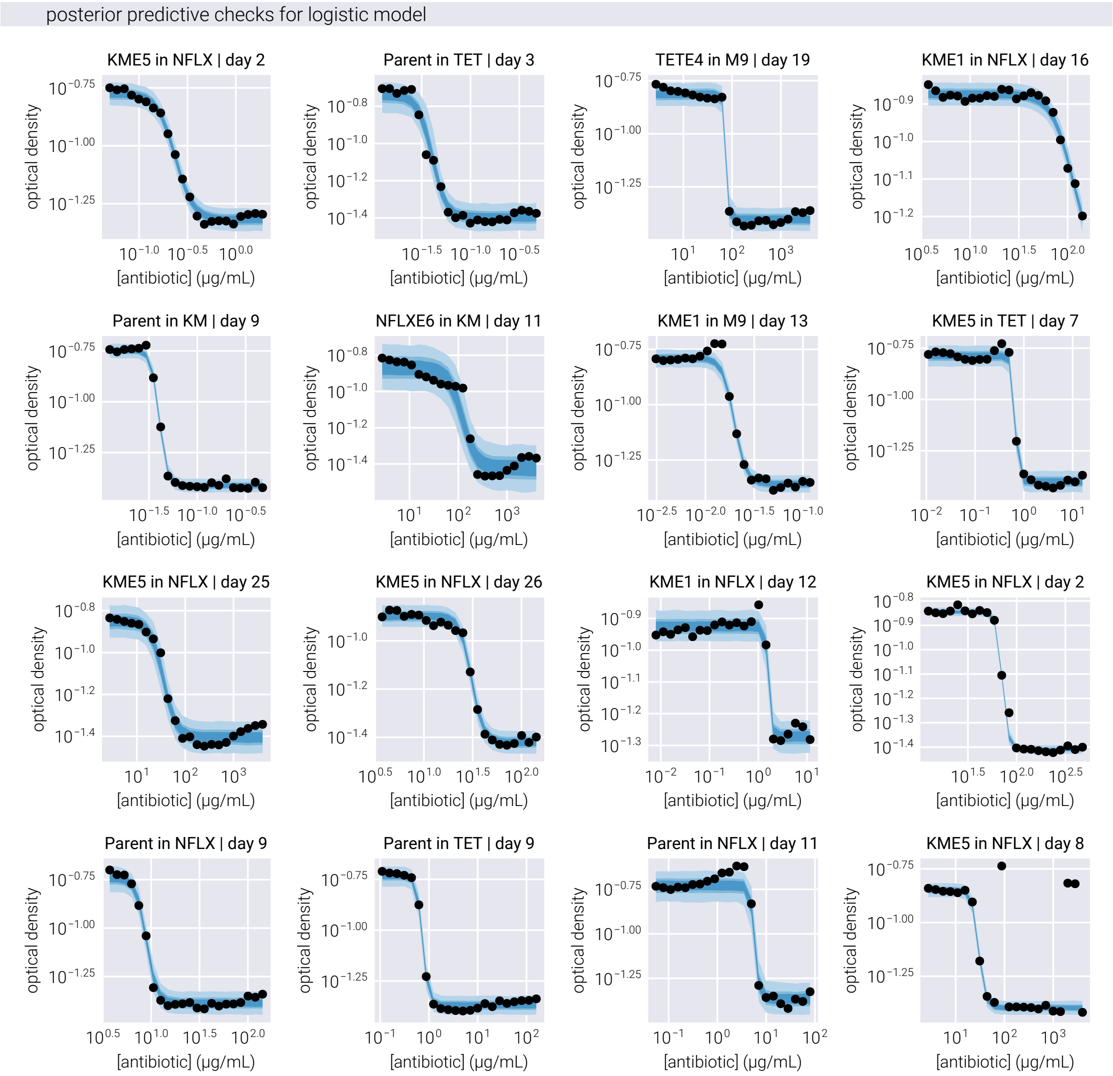

**Figure C. Posterior predictive check.** The posterior predictive check of the posterior distribution for the $IC_{50}$ parameter inference. The blue line is the mean of the posterior distribution, the different shades of blue represent the 95%, 68%, and 50% credible regions. The black dots are the raw optical density measurements as provided by Iwasawa et al. [1].

where $GP(\underline{x})$ is related to the probability of a genotype mapping to a phenotype $\underline{x}$. As a particular density function we will consider a set of negative Gaussian peaks that will serve as "mutational barriers" that limit the range of phenotypes that can be reached. The deeper the peak, the less probable it is that a genotype will map to a phenotype within the peak. This idea captured by the following mutational landscape

$$GP(\underline{x}) = C - \sum_{p=1}^{P} B_p \exp\left(-\frac{1}{2}(\underline{x} - \underline{\hat{x}}^{(p)})^T \Sigma_p^{-1} (\underline{x} - \underline{\hat{x}}^{(p)})\right), \tag{S17}$$

where $C$ is a constant offset—set to be the negative of the lowest peak value to have the maximum value of 1, and $B_p$ is the depth of the $p$-th peak, and $\underline{\hat{x}}^{(p)}$ is the $N$-dimensional vector of optimal phenotypes for the $p$-th peak.

##### B.2. Metropolis-Hastings Algorithm

The Metropolis-Hastings algorithm is a Markov Chain Monte Carlo (MCMC) method used to sample from a target probability distribution $\pi(\underline{x})$. In our evolutionary context, $\pi(\underline{x})$ can be thought of as the steady-state distribution of phenotypes under the combined influence of selection and mutation. We can define this distribution as

$$\pi(\underline{x}) \propto F_E(\underline{x})^{\beta_F} GP(\underline{x})^{\beta_{GP}}, \tag{S18}$$

where $\beta_F$ and $\beta_{GP}$ are parameters that separately control the influence of fitness and genotype-phenotype density, respectively. $\beta_F$ can be interpreted as a parameter controlling the strength of selection, akin to an effective population size, while $\beta_{GP}$ controls the ruggedness of the mutational landscape. For the purposes of this study, we made the simplifying assumption that $\beta_F = \beta_{GP} = \beta$, as this was sufficient to explore the effects of varying evolutionary stochasticity on the overall landscape. This parameter $\beta$ can be seen as an inverse temperature parameter (from statistical mechanics) that controls the level of stochasticity in the system. A higher $\beta$ means the system is more likely to move towards higher fitness and genotype-phenotype density. In a sense, $\beta$ can be related to the effective population size, controlling the directness of the evolutionary dynamics.

At each step, we propose a new phenotype $\underline{x}'$ from a proposal distribution $q(\underline{x}'|\underline{x})$. For simplicity, we can use a symmetric proposal distribution, such as a Gaussian centered at the current phenotype. Further, we constrain the proposal distribution to be symmetric, i.e.,

$$q(\underline{x}'|\underline{x}) = q(\underline{x}|\underline{x}'). \tag{S19}$$

This symmetry simplifies the acceptance probability later on.

The acceptance probability $P_{\text{accept}}$ for moving from $\underline{x}$ to $\underline{x}'$ is given by

$$P_{\text{accept}} = \min\left(1, \frac{\pi(\underline{x}')q(\underline{x}|\underline{x}')}{\pi(\underline{x})q(\underline{x}'|\underline{x})}\right). \tag{S20}$$

Given the symmetry of the proposal distribution, $q(\underline{x}'|\underline{x}) = q(\underline{x}|\underline{x}')$ , so the acceptance probability simplifies to

$$P_{\text{accept}} = \min\left(1, \frac{\pi(\underline{x}')}{\pi(\underline{x})}\right). \tag{S21}$$

Substituting Equation S18 into Equation S21, we get the general acceptance probability as

$$P_{\text{accept}} = \min\left(1, \frac{F_E(\underline{x}')^{\beta_F} GP(\underline{x}')^{\beta_{GP}}}{F_E(\underline{x})^{\beta_F} GP(\underline{x})^{\beta_{GP}}}\right). \tag{S22}$$

Given our simplifying assumption that $\beta_F = \beta_{GP} = \beta$, this can be rewritten as

$$P_{\text{accept}} = \min\left(1, \frac{F_E(\underline{x}')^{\beta} GP(\underline{x}')^{\beta}}{F_E(\underline{x})^{\beta} GP(\underline{x})^{\beta}}\right), \tag{S23}$$

This expression shows that the acceptance probability depends on the difference in fitness and genotype-phenotype density between the current and proposed phenotypes.

##### B.3. Effect of Inverse Temperature

Figure D shows the effect of the inverse temperature parameter $\beta$ influences evolutionary trajectories in our Metropolis-Hastings framework. Through a series of panels showing both phenotypic trajectories (upper) and fitness dynamics (lower), we can observe how different $\beta$ values affect the balance between exploration and exploitation in phenotypic space.

At low $\beta$ values (left panels), the evolutionary trajectories exhibit highly stochastic behavior, with populations taking meandering paths through the phenotypic landscape. This represents a regime where genetic drift plays a dominant role. As $\beta$ increases (moving right), the trajectories become increasingly deterministic, with populations taking more direct paths toward fitness peaks, while avoiding genotype-to-phenotype density valleys. This transition reflects a shift from drift-dominated to selection-dominated dynamics, where populations more strictly follow the local fitness gradient. The fitness trajectories in the lower panels further reinforce this pattern, showing more erratic fitness fluctuations at low $\beta$ and more monotonic increases at high $\beta$, consistent with stronger selective pressures.

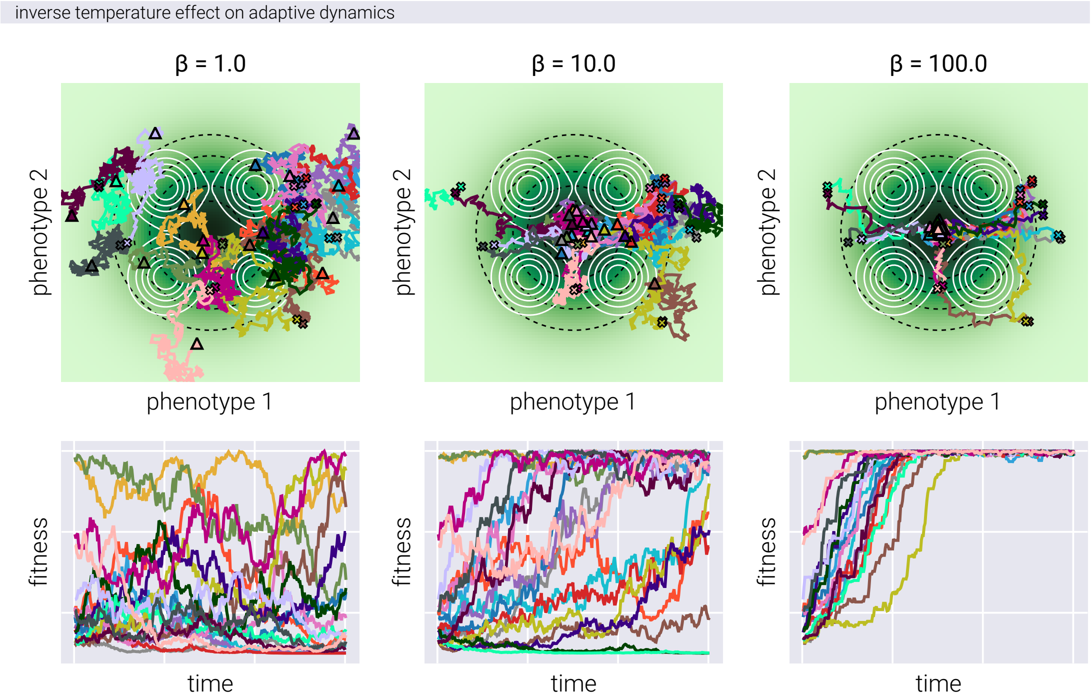

**Figure D. Effect of inverse temperature on adaptive dynamics**. Upper panels: Population trajectories on a landscape with a single fitness peak and four genotype-phenotype density peaks. Black dashed lines indicate fitness landscape contours, while white solid lines show genotype-phenotype density contours. Colored lines represent independent population trajectories, with initial positions (crosses) and final positions (triangles). Lower panels: Temporal evolution of population fitness. Panels from left to right show increasing values of inverse temperature ($\beta$), demonstrating increasingly deterministic adaptive trajectories.

### C. Riemannian Hamiltonian Variational Autoencoders Mathematical Background

The bulk of this work makes use of the variational autoencoder variation developed by [3] known as Riemannian Hamiltonian Variational Autoencoder (RHVAE). Understanding this model requires some background that we provide in the following sections. It is not necessary to follow every mathematical derivation in detail for these sections, as they don't directly pertain to the main contribution of this work. However, we believe that some intuition behind some of the concepts that inspired the RHVAE model can go a long way to understand the model. We invite the reader to refer to the original paper [3] and the references therein for additional details.

Let us start by reviewing the variational autoencoder model setting and the variational inference framework.

#### C.1. Variational Autoencoder Model Setting

Given a dataset $\underline{x}^{1:N} = \{\underline{x}_1, \underline{x}_2, \ldots, \underline{x}_N\}$, where each individual observation $\underline{x}_i \in \mathcal{X} \subseteq \mathbb{R}^D$, where the superscript denotes a set of $N$ observations, the variational autoencoder aims to fit a joint distribution $\pi_\theta(\underline{x}, \underline{z})$ where the data generative process is assumed to involve a set of latent—unobserved—continuous variables $\underline{z} \in \mathcal{Z} \subseteq \mathbb{R}^d$, where usually $d \ll D$. In other words, the VAE assumes that the data we get to observe is a consequence of a set of hidden variables $\underline{z}$ that we cannot measure directly. We assume that these variables live in a much lower-dimensional space than the one we observe and we are trying to learn this joint distribution such that, if desired, we can generate new data points by sampling from this distribution. Since the data we observe is a "consequence" of the latent variables, we express this joint distribution as

$$\pi_\theta(\underline{x}, \underline{z}) = \pi_\theta(\underline{x} | \underline{z})\, \pi_{\text{prior}}(\underline{z}), \tag{S24}$$

where $\pi_{\text{prior}}(\underline{z})$ is the prior distribution over latent variables and $\pi_\theta(\underline{x}|\underline{z})$ is the likelihood of the data given the latent variables. This looks exactly as the problem one confronts when performing Bayesian inference. What distinguishes the VAE from other Bayesian inference problems, as we will see, is that in many cases, we do not know the functional form of both the prior and the likelihood function that generated our data, but we have enough data samples that we hope to be able to learn these function using the neural networks function approximation capabilities. Thus, the objective of the VAE is to parameterize this likelihood function via a neural network with parameters $\theta$ given some prior distribution over the latent variables. For simplicity, it is common to choose a standard normal prior distribution, i.e., $\pi_{\text{prior}}(\underline{z}) = \mathcal{N}(\underline{0}, \underline{\underline{I}}_d)$, where $\underline{\underline{I}}_d$ is the $d$-dimensional identity matrix [4, 5]. This way, once we have learned the parameters of the neural network, we can generate new data points by sampling from this prior distribution and running the resulting samples through the likelihood function encoded by this neural network.

Our objective then becomes to find the parameters $\theta$ that maximize the probability of observing the data we actually observed. This is equivalent to maximizing the so-called marginal likelihood of the data, i.e.,

$$\max_{\theta} \pi_{\theta}(\underline{x}) = \max_{\theta} \int d^d\underline{z}\, \pi_{\theta}(\underline{x}, \underline{z}). \tag{S25}$$

But there are two problems with this objective:

1. The marginalization over the latent variables is intractable for most interesting distributions.
2. The distribution $\pi_{\theta}(\underline{x})$ is in general unknown and we need to somehow approximate it.

The way around these problems is to introduce a family of parametric distributions $q_{\phi}(\underline{z} \mid \underline{x})$, i.e., distributions that can be completely characterized by a set of parameters (think of Gaussians being a parametric family on the mean and the variance), and to approximate the true marginal distribution with one of these parametric distributions. This conceptual change is at the core of approximate methods for Bayesian inference known as variational inference. The objective then becomes to find the parameters $\phi$ that, when used to approximate the posterior distribution $\pi_{\theta}(\underline{z}|\underline{x})$, it is as close as possible to this posterior distribution. To mathematize this objective, we establish that we want to minimize the Kullback-Leibler divergence (also known as the relative entropy) between the variational distribution $q_{\phi}(\underline{z} \mid \underline{x})$ and the posterior distribution $\pi_{\theta}(\underline{z} \mid \underline{x})$, i.e., we aim to find $q^*_{\phi}$ such that

$$q^*_{\phi}(\underline{z} \mid \underline{x}) = \min_{\phi} D_{KL}\left(q_{\phi}(\underline{z} \mid \underline{x}) \| \pi_{\theta}(\underline{z} \mid \underline{x})\right). \tag{S26}$$

After some algebraic manipulation involving the well-known Gibbs inequality, we can show that this objective is equivalent to maximizing the so-called evidence lower bound (ELBO), $\mathcal{L}$, of the marginal likelihood, i.e.,

$$\mathcal{L} = \left\langle \log \pi_{\theta}(\underline{x}, \underline{z}) - \log q_{\phi}(\underline{z} \mid \underline{x}) \right\rangle_{q_{\phi}} \leq \log \pi_{\theta}(\underline{x}). \tag{S27}$$

Thus, the optimization problem we want to solve invovles estimating the ELBO given some data $\underline{x}$. However, the ELBO is an expectation over the variational distribution $q_{\phi}(\underline{z} \mid \underline{x})$, which is unknown. Therefore, we need to somehow estimate this expectation. Arguably, the most important contribution of the VAE framework is the introduction of the reparametrization trick [4]. This seemingly innocuous trick allows us to compute an unbiased estimate of the ELBO and its gradient, such that we can maximize it via gradient ascent. For more details on the reparametrization trick, we refer the reader to [4].

More recent attempts have focused on modifying $q_{\phi}(\underline{z} \mid \underline{x})$ to get a better approximation of the true posterior $\pi_{\theta}(\underline{z} \mid \underline{x})$. The basis of the Riemannian Hamiltonian Variational

Autoencoder (RHVAE) framework takes inspiration from some of these approaches. Let's gain some intuition on each of them.

##### C.2. MCMC and VI (Salimans & Kingma 2015)

One such approach to improve the accuracy of the variational posterior approximation consists in adding a fix number of Markov Chain Monte Carlo (MCMC) steps to the variational posterior approximation, targeting the true posterior [6]. In MCMC, rather than optimizing the parameters of a parametric distribution, we sample an initial position $\underline{z}_0$ from a simple distribution $q(\underline{z}_0)$ or $q(\underline{z}_0 \mid \underline{x})$ and subsequently, apply a stochastic transition operator $T(\underline{z}_{t+1} \mid \underline{z}_t)$ to draw a new value $\underline{z}_{t+1}$ from the distribution

$$\underline{z}_{t+1} \sim T(\underline{z}_{t+1} \mid \underline{z}_t). \tag{S28}$$

Applying this operator over and over again, we build a Markov chain

$$\underline{z}_0 \rightarrow \underline{z}_1 \rightarrow \underline{z}_2 \rightarrow \cdots \rightarrow \underline{z}_\tau, \tag{S29}$$

whose long-term behavior—the so-called stationary distribution—is a very good approximation of the true posterior distribution $\pi_\theta(\underline{z} \mid \underline{x})$. In other words, by carefully engineering the transition operator $T(\underline{z}_{t+1} \mid \underline{z}_t)$, and applying it over and over again, the "histogram" of the samples $\{\underline{z}_t\}_{t=0}^{\tau}$ will converge to the true posterior distribution $\pi_\theta(\underline{z} \mid \underline{x})$.

The central idea to combine MCMC with variational inference is to interpret the Markov chain

$$q_\phi(\underline{z}|\underline{x}) = q_\phi(\underline{z}_0 \mid \underline{x}) \prod_{t=1}^{\tau} T(\underline{z}_t \mid \underline{z}_{t-1}, \underline{x}) \tag{S30}$$

as a variational approximation in an expanded space of variables that includes the original latent variables $\underline{z}$ and the auxiliary variables $\underline{Z} = \{\underline{z}_0, \underline{z}_1, \ldots, \underline{z}_{\tau-1}\}$. Note that $\underline{Z}$ reaches up to $\tau - 1$ steps, since we consider the final step $\underline{z}_\tau$ to be the original latent variable $\underline{z}$. In other words, our variational distribution now writes

$$q_\phi(\underline{z}_\tau, \underline{Z} \mid \underline{x}) = q_\phi(\underline{z}_0 \mid \underline{x}) \prod_{t=1}^{\tau} T(\underline{z}_t \mid \underline{z}_{t-1}, \underline{x}). \tag{S31}$$

Integrating this expression into the ELBO (Equation S27) consists of substituting the variational distribution $q_\phi(\underline{z} \mid \underline{x})$ with the expression in Equation S31, i.e.,

$$\mathcal{L}_{\text{aux}} = \left\langle \log\left[\pi_\theta\left(\underline{x}, \underline{z}_T\right) r\left(\underline{Z} \mid \underline{z}_T, \underline{x}\right)\right] - \log q\left(\underline{Z}, \underline{z}_T \mid \underline{x}\right)\right\rangle_{q(\underline{Z}, \underline{z}_T|\underline{x})} \tag{S32}$$

where $r(\underline{Z} \mid \underline{z}_T, \underline{x})$ is an auxiliary inference distribution that we can choose freely. This auxiliary distribution is necessary to ensure we account for the auxiliary variables $\underline{Z}$ that are now part of our variational distribution. By splitting the joint distribution in Equation S31 as

$$q_\phi(\underline{z}_\tau, \underline{Z} \mid \underline{x}) = q_\phi(\underline{Z} \mid \underline{z}_\tau, \underline{x}) q_\phi(\underline{z}_\tau \mid \underline{x}), \tag{S33}$$

and substituting this expression into Equation S32, we get

$$\mathcal{L}_{\text{aux}} = \left\langle \log\left[\pi_\theta\left(\underline{x}, \underline{z}_T\right) r\left(\underline{Z} \mid \underline{z}_T, \underline{x}\right)\right] - \log\left[q_\phi(\underline{Z} \mid \underline{z}_\tau, \underline{x}) q_\phi(\underline{z}_\tau \mid \underline{x})\right]\right\rangle_{q_\phi(\underline{Z}|\underline{z}_\tau, \underline{x}) q_\phi(\underline{z}_\tau|\underline{x})}. \tag{S34}$$

Rearranging the terms, we get

$$\begin{aligned}
\mathcal{L}_{\text{aux}} &= \left\langle \log \frac{\pi_\theta\left(\underline{x}, \underline{z}_\tau\right)}{q_\phi(\underline{z}_\tau \mid \underline{x})} - \log \frac{q_\phi(\underline{Z} \mid \underline{z}_\tau, \underline{x})}{r(\underline{Z} \mid \underline{z}_\tau, \underline{x})} \right\rangle_{q_\phi(\underline{Z}|\underline{z}_\tau, \underline{x}) q_\phi(\underline{z}_\tau|\underline{x})}, \\
&= \left\langle \log \frac{\pi_\theta\left(\underline{x}, \underline{z}_\tau\right)}{q_\phi(\underline{z}_\tau \mid \underline{x})} \right\rangle_{q_\phi(\underline{Z}|\underline{z}_\tau, \underline{x}) q_\phi(\underline{z}_\tau|\underline{x})} - \left\langle \log \frac{q_\phi(\underline{Z} \mid \underline{z}_\tau, \underline{x})}{r(\underline{Z} \mid \underline{z}_\tau, \underline{x})} \right\rangle_{q_\phi(\underline{Z}|\underline{z}_\tau, \underline{x}) q_\phi(\underline{z}_\tau|\underline{x})}.
\end{aligned} \tag{S35}$$

From here, we note that the first term in Equation S35 is the original ELBO with the extra expectation taken over the auxiliary variables $\underline{Z}$. However, this term does not depend on the auxiliary variables $\underline{Z}$, thus, we can take it out of the expectation, i.e.,

$$\mathcal{L}_{\text{aux}} = \mathcal{L} - \left\langle \log \frac{q_\phi(\underline{Z} \mid \underline{z}_\tau, \underline{x})}{r(\underline{Z} \mid \underline{z}_\tau, \underline{x})} \right\rangle_{q_\phi(\underline{Z}|\underline{z}_\tau, \underline{x}) q_\phi(\underline{z}_\tau|\underline{x})}. \tag{S36}$$

For the second term, taking the expectation over the auxiliary variables makes it equivalent to the KL divergence between the variational distribution $q_\phi(\underline{Z} \mid \underline{z}_\tau, \underline{x})$ and the auxiliary distribution $r(\underline{Z} \mid \underline{z}_\tau, \underline{x})$, i.e.,

$$\mathcal{L}_{\text{aux}} = \mathcal{L} - \left\langle D_{KL}\left[q\left(\underline{Z} \mid \underline{z}_T, \underline{x}\right) \| r\left(\underline{Z} \mid \underline{z}_T, \underline{x}\right)\right]\right\rangle_{q(\underline{z}_T|\underline{x})} \leq \mathcal{L} \leq \log[\pi_\theta(\underline{x})]. \tag{S37}$$

where the inequality follows from the non-negativity of the KL divergence and the fact that the ELBO is a lower bound to the marginal likelihood. Since the variational distribution $q_\phi(\underline{z}_\tau, \mid \underline{x})$ we care about comes from the marginalization of the variational distribution over the auxiliary variables $\underline{Z}$,

$$q_\phi(\underline{z}_\tau \mid \underline{x}) = \int d\underline{Z}\, q_\phi(\underline{Z}, \underline{z}_\tau \mid \underline{x}), \tag{S38}$$

it is now a very rich family of distributions that can be used to better approximate the true posterior distribution $\pi_\theta(\underline{z} \mid \underline{x})$. However, we are still left with the task of choosing the auxiliary distribution $r(\underline{Z} \mid \underline{z}_\tau, \underline{x})$. What Salimans & Kingma [6] propose is to define this distribution also as a Markov chain, but in reverse order, i.e.,

$$r(\underline{Z} \mid \underline{z}_\tau, \underline{x}) = \prod_{t=1}^{\tau} T^\dagger(\underline{z}_{t-1} \mid \underline{z}_t, \underline{x}), \tag{S39}$$

where $T^\dagger(\underline{z}_{t-1} \mid \underline{z}_t, \underline{x})$ is the reverse of the transition operator $T(\underline{z}_{t+1} \mid \underline{z}_t, \underline{x})$. Notice that the dependence on $\underline{z}_\tau$ is dropped.

This choice of the auxiliary distribution leads to an upper bound on the marginal log likelihood, of the form

$$\begin{aligned} \log \pi_\theta(\underline{x}) &\geq \left\langle \log \pi_\theta\left(\underline{x}, \underline{z}_\tau\right) - \log q_\phi\left(\underline{z}_0, \ldots, \underline{z}_\tau \mid \underline{x}\right) + \log r\left(\underline{z}_0, \ldots, \underline{z}_{\tau-1} \mid \underline{x}, \underline{z}_\tau\right)\right\rangle_{q_\phi} \\ &= \left\langle \log\left[\frac{\pi_\theta\left(\underline{x}, \underline{z}_\tau\right)}{q_\phi\left(\underline{z}_0 \mid \underline{x}\right)}\right] + \sum_{t=1}^{\tau} \log\left[\frac{T^\dagger\left(\underline{z}_{t-1} \mid \underline{z}_t, \underline{x}\right)}{T\left(\underline{z}_t \mid \underline{z}_{t-1}, \underline{x}\right)}\right]\right\rangle_{q_\phi}. \end{aligned} \tag{S40}$$

In other words, we can interpret the Markov chain as a parametric distribution over the latent variables $\underline{z}$ that we can use to approximate the true posterior distribution $\pi_\theta(\underline{z} \mid \underline{x})$. The unbiased estimator of the marginal likelihood is then given by

$$\hat{\pi}_\theta(\underline{x}) = \frac{\pi_\theta(\underline{x}, \underline{z}_T) \prod_{t=1}^{T} T^\dagger(\underline{z}_{t-1} \mid \underline{z}_t, \underline{x})}{q_\phi(\underline{z}_0 \mid \underline{x}) \prod_{t=1}^{T} T(\underline{z}_t \mid \underline{z}_{t-1}, \underline{x})}. \tag{S41}$$

Let's unpack this expression. When performing MCMC, we build a Markov chain

$$\underline{z}_0 \to \underline{z}_1 \to \underline{z}_2 \to \cdots \to \underline{z}_T, \tag{S42}$$

where $\underline{z}_0 \sim q_\phi(\underline{z} \mid \underline{x})$ is the initial position in the chain. At each step, we sample a new position from the transition kernel $T(\underline{z}_t \mid \underline{z}_{t-1}, \underline{x})$, which is a function of the current position and the observed data. This way, we build a Markov chain that converges to the true posterior distribution. Therefore, the denominator of Equation S41 computes the probability of the initial position of the chain $q_\phi(\underline{z}_0 \mid \underline{x})$ times the product of the transition probabilities for each of the $\tau$ steps, $\prod_{t=1}^{\tau} T(\underline{z}_t \mid \underline{z}_{t-1}, \underline{x})$.

The numerator of Equation S41 computes the equivalent probability, but in reverse order, i.e., starts by sampling the final position of the chain $\underline{z}_T$ from the joint distribution $\pi_\theta(\underline{x}, \underline{z}_T)$ and then samples the previous positions of the chain $\underline{z}_{T-1}, \underline{z}_{T-2}, \ldots, \underline{z}_0$ from the transition kernel run backwards, $T^\dagger(\underline{z}_{t-1} \mid \underline{z}_t, \underline{x})$. Including a Markov chain into the variational posterior approximation has the effect of trading off computational efficiency for better posterior approximation. In other words, to improve the quality of the posterior approximation we make use of arguably the most accurate family of inference methods known to date—Markov Chain Monte Carlo (MCMC)—at the cost of increased computational cost. Intuitively, our estimate of the unbiased gradient of the ELBO becomes more and more accurate the more steps we include in the Markov chain, but at the same time, the cost of computing this gradient becomes higher and higher.

##### C.3. Normalizing Flows (Rezende & Mohamed 2015)

Another approach to imrpove the posterior approximation is to consider the composition of simple smooth invertible transformations [7], such that a variable $\underline{z}_K$ consists of $K$ of such transformations of a random variable $\underline{z}_0$, sampled from a simple distribution, i.e., $\underline{z}_K$ is built from chaining $K$ simple transformations, $f_x^k$, of $\underline{z}_0$,

$$\underline{z}_K = f_x^K \circ f_x^{K-1} \circ \cdots \circ f_x^1(\underline{z}_0), \tag{S43}$$

where $f_i \circ f_j(x) = f_i(f_j(x))$. Because we define the transformations to be smooth and invertible, the change of variables formula tells us that the density of $\underline{z}_K$ writes

$$q_\phi(\underline{z}_K \mid \underline{x}) = q_\phi(\underline{z}_0 \mid \underline{x}) \prod_{k=1}^{K} |\det J_{f_x^k}|^{-1}, \tag{S44}$$

where

$$J_{f_x^k} = \frac{\partial f_x^k}{\partial \underline{z}}, \tag{S45}$$

is the Jacobian matrix of the transformation $f_x^k$. In other words, the transformation of the random variable upon a composition of simple transformations is a product of the Jacobians of each transformation. Thus, if we choose a sequence of simple transformations, with a simple Jacobian matrix, we can construct a more complex distribution over the latent variables. Again, this has the effect of trading off computational efficiency for better posterior approximation, this time in the form of smooth invertible transformations of the original variational distribution. In other words, say that approximating the posterior distribution using a simple distribution, such as a Gaussian, is too simple. We can use a sequence of smooth invertible transformations to transform this simple distribution into a more complex one that can better approximate the true posterior. This again, comes at the cost of increased computational cost.

##### C.4. Hamiltonian Markov Chain Monte Carlo (HMC)

Although seemingly unrelated to the previous two approaches, the Hamiltonian Markov Chain Monte Carlo (HMC) framework provides a way to construct a Markov chain that is informed by the target distribution we are trying to approximate. This property is exploited by the predecessor of the RHVAE framework, the Hamiltonian Variational Autoencoder that we will review in the next section. But first, let us give a brief overview of the main ideas of the HMC framework. We direct the interested reader to the excellent review [8] for a more detailed treatment of the topic.

In the HMC framework, a random variable $\underline{z}$ is assumed to live in an Euclidean space with a target density $\pi(\underline{z} \mid \underline{x})$ derived from a potential $U_{\underline{x}}(\underline{z})$, such that the distribution writes

$$\pi(\underline{z} \mid \underline{x}) = \frac{e^{-U_{\underline{x}}(\underline{z})}}{\int d^d\underline{z}\, e^{-U_{\underline{x}}(\underline{z})}}, \tag{S46}$$

where we define the potential energy as

$$U_{\underline{x}}(\underline{z}) \equiv -\log \pi(\underline{z} \mid \underline{x}). \tag{S47}$$

Notice that upon substitution of this definition into Equation S46, we get the trivial equality

$$\pi(\underline{z} \mid \underline{x}) = \frac{\exp[-(-\log \pi(\underline{z} \mid \underline{x}))]}{\int d^d\underline{z}\, \exp[-(-\log \pi(\underline{z} \mid \underline{x}))]} = \frac{\pi(\underline{z} \mid \underline{x})}{\int d^d\underline{z}\, \pi(\underline{z} \mid \underline{x})} = \pi(\underline{z} \mid \underline{x}). \tag{S48}$$

Since it is most of the time impossible to sample directly from $\pi(\underline{z} \mid \underline{x})$, an independent auxiliary variable $\underline{\rho} \in \mathbb{R}^d$ is introduced and used to sample $\underline{z}$ more efficiently. This variable—referred as the momentum—is such that:

$$\underline{\rho} \sim \mathcal{N}(\underline{0}, \underline{\underline{M}}), \tag{S49}$$

where $\underline{\underline{M}}$ is the so-called mass matrix. The idea behind HMC is then to work with a distribution that extends the target distribution $\pi(\underline{z} \mid \underline{x})$ to a distribution over both position and momentum variables, i.e.,

$$\pi(\underline{z}, \underline{\rho} \mid \underline{x}) = \pi(\underline{z} \mid \underline{x}, \underline{\rho})\pi(\underline{\rho} \mid \underline{x}) = \pi(\underline{z} \mid \underline{x})\pi(\underline{\rho}), \tag{S50}$$

where we assume the value of $\underline{z}$ does not directly depend on the momentum variable $\underline{\rho}$ but only on the observed data $\underline{x}$, and that the momentum distribution $\pi(\underline{\rho})$ is independent of all other variables. Using Equation S46, the density of this extended distribution can be analogously written as

$$\pi(\underline{z}, \underline{\rho} \mid \underline{x}) = \frac{e^{-H_{\underline{x}}(\underline{z}, \underline{\rho})}}{\iint d^d\underline{z}\, d^d\underline{\rho}\, e^{-H_{\underline{x}}(\underline{z}, \underline{\rho})}}. \tag{S51}$$

where $H_{\underline{x}}(\underline{z}, \underline{\rho})$ is the so-called Hamiltonian, defined as

$$H_{\underline{x}}(\underline{z}, \underline{\rho}) = -\log \pi(\underline{z}, \underline{\rho} \mid \underline{x}) = -\log \pi(\underline{z} \mid \underline{x}) - \log \pi(\underline{\rho}). \tag{S52}$$

Substituting $\pi(\underline{\rho})$ from Equation S49 into Equation S52 gives

$$\begin{aligned} H_{\underline{x}}(\underline{z}, \underline{\rho}) &= -\log \pi(\underline{z} \mid \underline{x}) + \frac{1}{2}\left[\log((2\pi)^d |\underline{\underline{M}}|) + \underline{\rho}^T \underline{\underline{M}}^{-1} \underline{\rho}\right] \\ &= U_{\underline{x}}(\underline{z}) + \kappa(\underline{\rho}). \end{aligned} \tag{S53}$$

In physical systems, the Hamiltonian gives the total energy of a system having a position $\underline{z}$ and a momentum $\underline{\rho}$. $U_{\underline{x}}(\underline{z})$ is referred as the potential energy (since it is position-dependent) and $\kappa(\underline{\rho})$ is the kinetic energy, since it is momentum-dependent.

The point of writing the extended target distribution in this form is that we can take advantage of one specific desirable property of Hamiltonian dynamics: Hamiltonian flows are volume preserving [8]. In other words, the volume in phase space defined by the position and momentum variables is invariant under Hamiltonian dynamics. What this means for our purposes of sampling from a posterior distribution $\pi(\underline{z} \mid \underline{x})$ is that a walker starting in some position $\underline{z}$ can "traverse" the volume of the target distribution very efficiently traveling through lines of equal probability. In this sense, the Hamiltonian dynamics can be seen as a way to explore the posterior distribution $\pi(\underline{z} \mid \underline{x})$ very efficiently. To do so, we compute the time evolution of a walker exploring the space of the position and momentum variables using Hamilton's equations

$$\begin{aligned} \frac{d\underline{z}}{dt} &= \frac{\partial H_{\underline{x}}}{\partial \underline{\rho}}, \\ \frac{d\underline{\rho}}{dt} &= -\frac{\partial H_{\underline{x}}}{\partial \underline{z}}. \end{aligned} \tag{S54}$$

One can show that for the particular choice of momentum distribution in Equation S49, these equations take the form

$$\begin{aligned} \frac{d\underline{z}}{dt} &= \underline{\underline{M}}^{-1} \underline{\rho}, \\ \frac{d\underline{\rho}}{dt} &= -\nabla_{\underline{z}} \log \pi(\underline{z} \mid \underline{x}), \end{aligned} \tag{S55}$$

where $\nabla_{\underline{z}}$ is the gradient with respect to the position variable $\underline{z}$. Unfortunately, the resulting system of partial differential equations is almost always intractable analytically. Therefore, we must use specialized numerical integrators to solve it. The most popular choice is the so-called leapfrog integrator, which is a symplectic integrator that preserves the volume of the phase space. To update the position and momentum variables, the leapfrog integrator takes three steps:

1. A half step for the momentum:

$$\underline{\rho}\left(t+\frac{\epsilon}{2}\right)=\underline{\rho}(t)-\frac{\epsilon}{2}\nabla_{\underline{z}}H_{\underline{x}}\left(\underline{z}(t),\underline{\rho}(t)\right). \tag{S56}$$

2. A full step for the position:

$$\underline{z}\left(t+\epsilon\right)=\underline{z}(t)+\epsilon\nabla_{\underline{\rho}}H_{\underline{x}}\left(\underline{z}(t),\underline{\rho}\left(t+\frac{\epsilon}{2}\right)\right). \tag{S57}$$

3. A half step for the momentum:

$$\underline{\rho}(t+\epsilon)=\underline{\rho}\left(t+\frac{\epsilon}{2}\right)-\frac{\epsilon}{2}\nabla_{\underline{z}}H_{\underline{x}}\left(\underline{z}(t+\epsilon),\underline{\rho}\left(t+\frac{\epsilon}{2}\right)\right), \tag{S58}$$

where $\epsilon$ is the so-called leapfrog step size.

Without going further into the details of the HMC framework, we hope the reader can imagine how using this symplectic integrator, a walker aiming to explore the posterior distribution $\pi(\underline{z} \mid \underline{x})$ can traverse the volume of the target distribution very efficiently.

With this conceptual background, we are now ready to review the predecessor of the RHVAE framework, the Hamiltonian Variational Autoencoder framework.

#### C.5. Hamiltonian Variational Autoencoder (HVAE)

Given the dynamics defined in Equation S55, and the numerical integrator defined in Equation S56, Equation S57, and Equation S58, we can now take $K$ iterations of the leapfrog integrator to sample a point $(\underline{z}_K, \underline{\rho}_K)$ from the extended target distribution defined in Equation S50. This transition from $(\underline{z}_0, \underline{\rho}_0)$ to $(\underline{z}_K, \underline{\rho}_K)$ via the symplectic integrator can be thought of as a transformation of the form

$$\{\phi^{(K)}_{\epsilon,\underline{x}} : \mathbb{R}^d \times \mathbb{R}^d \to \mathbb{R}^d \times \mathbb{R}^d\}, \tag{S59}$$

i.e., $\phi^{(K)}_{\epsilon,\underline{x}}$ is a function that takes some input value $(\underline{z}, \underline{\rho})$ and advances it $K$ steps in phase space using the leapfrog integrator. The function includes $\epsilon$ to remind the step size of the integrator and $\underline{x}$ to remind its data dependence. To advance $\ell$ steps, we simply compose a one step iteration $(\ell = 1)$ with the $\ell - 1$ function, i.e.,

$$\phi^{(\ell)}_{\epsilon,\underline{x}} = \phi^{(\ell-1)}_{\epsilon,\underline{x}} \circ \phi^{(1)}_{\epsilon,\underline{x}} \tag{S60}$$

By induction, we can see that $\phi_{\epsilon,\underline{x}}^{(\ell)}$ can be built by $\ell$ compositions of $\phi_{\epsilon,\underline{x}}^{(1)}$, defining the entire transformation as

$$\phi_{\epsilon,\underline{x}}^{(K)} = \phi_{\epsilon,\underline{x}}^{(1)} \circ \phi_{\epsilon,\underline{x}}^{(1)} \circ \cdots \circ \phi_{\epsilon,\underline{x}}^{(1)}. \tag{S61}$$

It is no coincidence that this resembles the composition of functions used in the normalizing flows framework described earlier in Equation S43. We can then think of a $K$-step leapfrog integrator trajectory that takes

$$\left(\underline{z}_0, \underline{\rho}_0\right) \to \left(\underline{z}_K, \underline{\rho}_K\right) \tag{S62}$$

as an invertible transformation formed by composing $K$ one-step leapfrog integrator trajectories. The original HVAE framework [9] includes an extra step in between each leapfrog step that we now review. This step consisted of a *tempering* step where an initial temperature $\beta_0$ is proposed and the momentum is decreased by a factor

$$\alpha_k = \sqrt{\frac{\beta_{k-1}}{\beta_k}}, \tag{S63}$$

after each leapfrog step $k$. This simply means that for step $k$, the momentum is computed as

$$\underline{\rho}_k = \alpha_k \, \underline{\rho}_{k-1}. \tag{S64}$$

The temperature is then updated as

$$\sqrt{\beta_k} = \left[\left(1 - \frac{1}{\sqrt{\beta_0}}\right)\frac{k^2}{K^2} + \frac{1}{\sqrt{\beta_0}}\right]^{-1}. \tag{S65}$$

This tempering step tries to produce an effect similar to that of Annealed Importance Sampling (AIS) [10], where the temperature is decreased gradually to produce a smoother distribution that can be better approximated by the variational distribution. Thus, the transformation $\mathcal{H}_{\underline{x}}$ used in [9] takes the form

$$\mathcal{H}_{\underline{x}} = g^K \circ \phi_{\epsilon,\underline{x}}^{(1)} \circ g^{K-1} \circ \cdots \circ \phi_{\epsilon,\underline{x}}^{(1)} \circ g^0 \circ \phi_{\epsilon,\underline{x}}^{(1)}. \tag{S66}$$

Since each transformation is smooth and differentiable, this entire transformation is amenable to the reparameterization trick used in the variational autoencoder framework. We can then use this transformation to access an unbiased estimator of the gradient of the ELBO with respect to the variational parameters $\phi$.

As mentioned in Section C.3, for a series of smooth invertible transformations that define a normalizing flow, the change of variables formula tells us that the density of the transformed variable writes

$$q_\phi(\underline{z}_K \mid \underline{x}) = q_\phi(\underline{z}_0 \mid \underline{x}) |\det J_{\mathcal{H}_{\underline{x}}}|^{-1}. \tag{S67}$$

For our extended posterior distribution, we have that the initial point is sampled as

$$(\underline{z}_0, \underline{\rho}_0) \sim q_\phi(\underline{z}_0, \underline{\rho}_0 \mid \underline{x}) = q_\phi(\underline{z}_0 \mid \underline{x})\, q_\phi(\underline{\rho}_0), \tag{S68}$$

meaning that only the initial position is sampled from the variational distribution, while the momentum is sampled from a standard normal distribution. For the Jacobian, each step consists of the composition of two functions:

1. The leapfrog integrator step $\phi^{(1)}_{\epsilon,\underline{x}}$.
2. The tempering step $g^k$.

Therefore, the determinant of the Jacobian of the transformation is given by the product of the determinants of each of the steps, i.e.,

$$|\det J_{\mathcal{H}_{\underline{x}}}| = \prod_{k=0}^{K} |\det J_{g^k}| |\det J_{\phi^{(1)}_{\epsilon,\underline{x}}}|. \tag{S69}$$

The resulting variational distribution is then given by

$$\begin{aligned} q_\phi(\underline{z}_K, \underline{\rho}_K \mid \underline{x}) &= q_\phi(\underline{z}_0 \mid \underline{x}) q_\phi(\underline{\rho}_0) |\det J_{\mathcal{H}_{\underline{x}}}|^{-1} \\ &= q_\phi(\underline{z}_0 \mid \underline{x}) q_\phi(\underline{\rho}_0) \left[ \prod_{k=0}^{K} \left|\det J_{g^k}\right| \left|\det J_{\phi^{(1)}_{\epsilon,\underline{x}}}\right| \right]^{-1} \end{aligned} \tag{S70}$$

However, recall that Hamiltonian flows are volume preserving. This means that the volume of the phase space is invariant under the transformation and thus the determinant of the Jacobian is equal to one, i.e.,

$$|\det J_{\phi^{(1)}_{\epsilon,\underline{x}}}| = 1. \tag{S71}$$

For the tempering step, since $(\underline{z}, \underline{\rho}) \to (\underline{z}, \alpha_k \underline{\rho})$, the determinant of the Jacobian is given by

$$|\det J_{g^k}| = \alpha_k^d = \left( \frac{\beta_{k-1}}{\beta_k} \right)^{\frac{d}{2}}. \tag{S72}$$

With this setting in hand [9] proposes an algorithm to estimate the gradient of the ELBO with respect to the variational parameters that expands the the usual procedure to includes a series of leapfrog steps and a tempering step (see Algorithm 1 in [9] for details).

##### C.6. Riemannian Hamiltonian MCMC

To improve the representational power of the variational posterior approximation, we can make the reasonable assumption that the latent variables $\underline{z}$ live in a Riemannian manifold $\mathcal{M}$ endowed with a Riemannian metric $\underline{\underline{G}}(\underline{z})$. Although not entirely correct, one can think of this Riemannian metric as some sort of "position dependent scale bar" for a flat map representing a curved space. In the context of the HMC framework, this Riemannian metric can be included if we allow the momentum variable to be given by

$$\underline{\rho} \sim \mathcal{N}\left(\underline{0}, \underline{\underline{G}}(\underline{z})\right), \tag{S73}$$

i.e., the auxiliary momentum variable is no longer independent of the position variable $\underline{z}$ but rather is distributed according to a normal distribution with mean zero and covariance matrix given by the position-dependent Riemannian metric $\underline{\underline{G}}(\underline{z})$. The Hamiltonian governing the dynamics of the system then becomes

$$H_{\underline{x}}^{\mathcal{M}}(\underline{z}, \underline{\rho}) = U_{\underline{x}}^{\mathcal{M}}(\underline{z}) + \kappa_{\underline{x}}^{\mathcal{M}}(\underline{z}, \underline{\rho}), \tag{S74}$$

where the $\mathcal{M}$ superscript reminds us that we now consider the latent variables $\underline{z}$ living on a Riemannian manifold. Although the potential energy $U_{\underline{x}}^{\mathcal{M}}(\underline{z})$ is still given by Equation S47, the kinetic energy term is now position-dependent, given by

$$\kappa_{\underline{x}}^{\mathcal{M}}(\underline{z}, \underline{\rho}) = \frac{1}{2}\log((2\pi)^d |\underline{\underline{G}}(\underline{z})|) + \frac{1}{2}\underline{\rho}^T \underline{\underline{G}}(\underline{z})^{-1} \underline{\rho} \tag{S75}$$

Our target distribution $\pi(\underline{z} \mid \underline{x})$ is still given by Equation S50, but with the Hamiltonian given by Equation S74. To show that we still recover the target distribution with this change in the Hamiltonian, let us substitute Equation S74 into Equation S50. This results in

$$\pi(\underline{z} \mid \underline{x}) = \frac{\displaystyle\int d^d\underline{\rho}\, \exp\left\{-U_{\underline{x}}^{\mathcal{M}}(\underline{z}) - \frac{1}{2}\log((2\pi)^d|\underline{\underline{G}}(\underline{z})|) - \frac{1}{2}\underline{\rho}^T\underline{\underline{G}}(\underline{z})^{-1}\underline{\rho}\right\}}{\displaystyle\iint d^d\underline{z}\, d^d\underline{\rho}\, \exp\left\{-U_{\underline{x}}^{\mathcal{M}}(\underline{z}) - \frac{1}{2}\log((2\pi)^d|\underline{\underline{G}}(\underline{z})|) - \frac{1}{2}\underline{\rho}^T\underline{\underline{G}}(\underline{z})^{-1}\underline{\rho}\right\}}. \tag{S76}$$

Rearranging terms, we can see that the target distribution is given by

$$\pi(\underline{z} \mid \underline{x}) = \frac{e^{-U_{\underline{x}}^{\mathcal{M}}(\underline{z})} \int d^d\underline{\rho} \left[ \frac{e^{-\frac{1}{2}\underline{\rho}^T \underline{\underline{G}}(\underline{z})^{-1}\underline{\rho}}}{(2\pi)^d \sqrt{|\underline{\underline{G}}(\underline{z})|}} \right]}{\int d^d\underline{z}\, e^{-U_{\underline{x}}^{\mathcal{M}}(\underline{z})} \int d^d\underline{\rho} \left[ \frac{e^{-\frac{1}{2}\underline{\rho}^T \underline{\underline{G}}(\underline{z})^{-1}\underline{\rho}}}{(2\pi)^d \sqrt{|\underline{\underline{G}}(\underline{z})|}} \right]}. \tag{S77}$$

where we can recognize the terms in square brackets as the probability density function of a normal distribution. Thus, integrating out the momentum variable $\underline{\rho}$ from the target distribution, we obtain

$$\pi(\underline{z} \mid \underline{x}) = \frac{e^{-U_{\underline{x}}^{\mathcal{M}}(\underline{z})}}{\int d^d\underline{z}\, e^{-U_{\underline{x}}^{\mathcal{M}}(\underline{z})}}, \tag{S78}$$

i.e., the correct target distribution as defined in Equation S46. Therefore, including the position-dependent Riemannian metric in the momentum variable does not change the target distribution. However, the same cannot be said for the Hamiltonian equations of motion, as we will see next.

Recall that the Hamiltonian equations in Equation S54 require us to compute the gradient of the potential energy with respect to the position variable $\underline{z}$ and the gradient of the kinetic energy with respect to the momentum variable $\underline{\rho}$. For the position variable, our result from Equation S55 still holds, with the only difference that the mass matrix is now given by the Riemannian metric $\underline{\underline{G}}(\underline{z})$. For a single entry of $z_i$, dynamics are given by

$$\frac{dz_i}{dt} = \frac{\partial H_{\underline{x}}^{\mathcal{M}}(\underline{z}, \underline{\rho})}{\partial \rho_i} = \left(\nabla_{\underline{z}} H_{\underline{x}}^{\mathcal{M}}(\underline{z}, \underline{\rho})\right)_i = \left(\underline{\underline{G}}(\underline{z})^{-1}\underline{\rho}\right)_i, \tag{S79}$$

where $(\cdot)_i$ denotes the $i$-th entry of a vector.

For the momentum variable, we use three relatively standard results from matrix calculus:

1. The gradient of the inverse of a matrix function with respect to the argument is given by

$$\frac{\partial \underline{\underline{G}}(\underline{z})^{-1}}{\partial \underline{z}} = -\underline{\underline{G}}(\underline{z})^{-1} \frac{d\underline{\underline{G}}(\underline{z})}{d\underline{z}} \underline{\underline{G}}(\underline{z})^{-1}. \tag{S80}$$

2. The gradient of the determinant of a matrix function with respect to the argument is given by

$$\frac{d}{d\underline{z}} \det \underline{\underline{G}}(\underline{z}) = \mathrm{tr}\left(\mathrm{adj}(\underline{\underline{G}}(\underline{z}))\frac{d\underline{\underline{G}}(\underline{z})}{d\underline{z}}\right) = \det \underline{\underline{G}}(\underline{z})\, \mathrm{tr}\left(\underline{\underline{G}}(\underline{z})^{-1}\frac{d\underline{\underline{G}}(\underline{z})}{d\underline{z}}\right), \tag{S81}$$

where $\mathrm{adj}$ is the adjoint operator, $\mathrm{tr}$ is the trace operator, and $\det$ is the determinant operator.

3. The gradient of the logarithm of the determinant of a matrix function with respect to the argument is given by

$$\frac{d}{d\underline{z}} \log(\det \underline{\underline{G}}(\underline{z})) = \mathrm{tr}\left(\underline{\underline{G}}(\underline{z})^{-1}\frac{d\underline{\underline{G}}(\underline{z})}{d\underline{z}}\right). \tag{S82}$$

Given these results, we can now compute the dynamics of the momentum variable. For a single entry of $\underline{\rho}$, $\rho_i$, the resulting expression is of the form

$$\frac{d\rho_i}{dt} = \frac{\partial H^{\mathcal{M}}_{\underline{x}}(\underline{z}, \underline{\rho})}{\partial z_i} = \frac{\partial \ln \pi(\underline{z} \mid \underline{x})}{\partial z_i} - \frac{1}{2}\mathrm{tr}\left(\underline{\underline{G}}(\underline{z})\frac{\partial \underline{\underline{G}}(\underline{z})}{\partial z_i}\right) + \frac{1}{2}\underline{\rho}^T \underline{\underline{G}}(\underline{z})^{-1}\frac{\partial \underline{\underline{G}}(\underline{z})}{\partial z_i}\underline{\underline{G}}(\underline{z})^{-1}\underline{\rho}. \tag{S83}$$

As before, we can adapt the leapfrog integrator to this setting to produce an unbiased estimator of the gradient of the ELBO with respect to the variational parameters. However, the previously presented leapfrog integrator is no longer volume preserving in this setting due to the position-dependent Riemannian metric. Thus, we need to use a generalized version of the leapfrog integrator. This has been previously derived and [3] lists the following steps for the generalized leapfrog integrator as

1. A half step for the momentum variable

$$\underline{\rho}\left(t + \frac{\varepsilon}{2}\right) = \underline{\rho}(t) - \frac{\varepsilon}{2}\nabla_{\underline{z}} H^{\mathcal{M}}_{\underline{x}}\left(\underline{z}(t), \underline{\rho}\left(t + \frac{\varepsilon}{2}\right)\right). \tag{S84}$$

2. A full step for the position variable

$$\underline{z}(t+\varepsilon) = \underline{z}(t) + \frac{\varepsilon}{2}\left[\nabla_{\underline{z}} H^{\mathcal{M}}_{\underline{x}}\left(\underline{z}(t), \underline{\rho}\left(t + \frac{\varepsilon}{2}\right)\right) + \nabla_{\underline{\rho}} H^{\mathcal{M}}_{\underline{x}}\left(\underline{z}(t+\varepsilon), \underline{\rho}\left(t + \frac{\varepsilon}{2}\right)\right)\right]. \tag{S85}$$

3. A half step for the momentum variable

$$\underline{\rho}(t+\varepsilon) = \underline{\rho}\left(t + \frac{\varepsilon}{2}\right) - \frac{\varepsilon}{2}\nabla_{\underline{z}} H^{\mathcal{M}}_{\underline{x}}\left(\underline{z}(t+\varepsilon), \underline{\rho}(t+\varepsilon)\right). \tag{S86}$$

Since the left hand side terms appear on the right hand side, these equations must be solved using fixed point iterations. This means that the numerical implementation of this generalized integrator iterates a step multiple times until it finds a "fixed point", i.e., a point that, when fed to the right hand side of the equation, does not change the value of the left hand side. In practice, we set the number of iterations to a small number.

##### C.7. Riemannian Hamiltonian Variational Autoencoder

The Riemannian Hamiltonian Variational Autoencoder (RHVAE) method extends the HVAE idea to account for non-Euclidean geometry of the latent space. This means that the latent variables $\underline{z}$ live on a Riemannian manifold. This manifold is endowed with a Riemannian metric $\underline{\underline{G}}(\underline{z})$, that is position-dependent. We navigate through this manifold using the Hamiltonian equations of motion, utilizing the geometric information encoded in the Riemannian metric.

As with the HAVE, using the generalized leapfrog integrator (Equation S84, Equation S85, Equation S86) along with a tempering step creates a smooth mapping

$$(\underline{z}_0, \underline{\rho}_0) \rightarrow (\underline{z}_K, \underline{\rho}_K), \tag{S87}$$

that can be thought of as a kind of normalizing flow informed by the target distribution and the geometry of the latent space. Using the same procedure as the HVAE that includes a tempering step, the resulting variational distribution takes the form of Equation S70, with the only difference that the momentum variable is now position-dependent, i.e.,

$$q_\phi(\underline{z}_K, \underline{\rho}_K \mid \underline{x}) = q_\phi(\underline{z}_0 \mid \underline{x}) q_\phi(\underline{\rho}_0 \mid \underline{z}_0) \left[ \prod_{k=1}^{K} \left| \det \underline{\underline{J}}_{g^k} \right| \left| \det \underline{\underline{J}}_{\phi^{(1)}_{\epsilon,\underline{x}}} \right| \right]^{-1}, \tag{S88}$$

where, as before, $\underline{\underline{J}}_{\phi^{(1)}_{\epsilon,\underline{x}}}$ is the Jacobian of the generalized leapfrog integrator with step size $\epsilon$, and $\underline{\underline{J}}_{g^k}$ is the Jacobian of the tempering step. Since we established that Hamiltonian dynamics are volume preserving, the determinant of the Jacobian of the tempering step is one, i.e., $\left| \det \underline{\underline{J}}_{\phi^{(1)}_{\epsilon,\underline{x}}} \right| = 1$. Moreover, since we are using the same type of tempering step as in the HVAE, we have the same result for the determinant of the Jacobian of the tempering step as in Equation S72. After substituting these results into Equation S88, we obtain

$$q_\phi(\underline{z}_K, \underline{\rho}_K \mid \underline{x}) = q_\phi(\underline{z}_0 \mid \underline{x}) q_\phi(\underline{\rho}_0 \mid \underline{z}_0) \beta_0^{-d/2}, \tag{S89}$$

where $d$ is the dimension of the latent space. The main difference from the HVAE is that the term $q_\phi(\underline{\rho}_0 \mid \underline{z}_0)$ depends on the position $\underline{z}_0$ of the latent variables. To establish this dependence, we must define the functional form for the metric tensor $\underline{\underline{G}}(\underline{z})$.

In RHVAE, the metric is learned from the data. However, looking at the generalized leapfrog integrator, we see that the inverse and the determinant of the metric tensor are required.

Thus, rather than defining the metric tensor directly, we define the inverse of the metric tensor $\underline{\underline{G}}(\underline{z})^{-1}$, and use the fact that

$$\det \underline{\underline{G}}(\underline{z}) = \det \underline{\underline{G}}(\underline{z})^{-1}. \tag{S90}$$

By doing so, we do not have to invert the metric tensor for every leapfrog step. Chadebec, Mantoux, and Allassonnière [3] proposes an inverse metric tensor of the form

$$\underline{\underline{G}}(\underline{z}) = \sum_{i=1}^{N} \underline{\underline{L}}_{\Psi_i} \underline{\underline{L}}_{\Psi_i}^{\top} \exp\left( -\frac{\left\| \underline{z} - \underline{\mu}(\underline{x}_i) \right\|_2^2}{T^2} \right) + \lambda \underline{\underline{\mathbb{I}}}_l, \tag{S91}$$

where $\underline{\underline{L}}_{\Psi_i}$ is a lower triangular matrix with positive diagonal entries, $\top$ is the transpose operator, $\underline{\mu}(\underline{x}_i)$ is the mean of the $i$-th component of the dataset, $T$ is a temperature parameter to smooth the metric tensor, $\lambda$ is a regularization parameter, and $\underline{\underline{\mathbb{I}}}_l$ is the $l \times l$ identity matrix. This last term with $\lambda$ is set for the metric tensor not to be zero. However, usually $\lambda$ is set to a small value, e.g., $10^{-3}$. The terms $\underline{\mu}(\underline{x}_i)$ are referred to as the "*centroids*" and are given by the mean of the variational posterior, such that,

$$q_\phi(\underline{z}_i \mid \underline{x}_i) = \mathcal{N}\left( \underline{\mu}(\underline{x}_i), \underline{\underline{\Sigma}}(\underline{x}_i) \right). \tag{S92}$$

Intuitively, we can think of $\underline{\underline{L}}_{\Psi_i}$ as the triangular matrix in the Cholesky decomposition of $\underline{\underline{G}}^{-1}(\underline{\mu}(\underline{x}_i))$ up to a regularization factor. This matrix is learned using a neural network with parameters $\Psi_i$, mapping

$$\underline{x}_i \rightarrow \underline{\underline{L}}_{\Psi_i}. \tag{S93}$$

The hyperparameters $T$ and $\lambda$ could be learned from the data as well, but, for simplicity, we set them to constants. We invite the reader to refer to [3] for a more detailed discussion on the training procedure.

#### D. Advantages of Nonlinear Manifold Learning for Fitness Landscapes

In this section, we provide a theoretical justification for why nonlinear dimensionality reduction methods may offer advantages over linear methods when modeling fitness landscapes. We build upon the concept of a causal manifold that captures the relationship between genotypes and their fitness across multiple environments.

#### D.1. Theoretical Framework: The Causal Manifold

The central concept in our analysis is what we call the causal manifold $\mathcal{M}_c$—a low-dimensional space that encodes the information necessary to predict fitness across multiple environments. Another way to think about this manifold is in a data-generating process perspective: the causal manifold is the underlying space from which the fitness data is sampled. We define two key functions:

1. The mapping from genotype to manifold coordinates:

$$\phi : g \in G \to \underline{z} \in \mathcal{M}_c. \tag{S94}$$

2. The mapping from manifold coordinates to fitness in environment $E$:

$$F_E : \underline{z} \in \mathcal{M}_c \to f_E \in \mathbb{R}. \tag{S95}$$

A critical assumption in linear approaches is that $F_E$ takes the form

$$F_E(\underline{z}) = \underline{\beta} \cdot \underline{z} + b, \tag{S96}$$

where $\underline{\beta}$ is a vector of coefficients and $b$ is a scalar offset. However, this assumption may not hold in general. While local linearity is guaranteed by Taylor's theorem for small changes in $\underline{z}$

$$F_E(\underline{z} + \Delta\underline{z}) \approx F_E(\underline{z}) + \nabla F_E(\underline{z}) \cdot \Delta\underline{z} + \mathcal{O}(\Delta\underline{z}^2), \tag{S97}$$

global linearity across the entire manifold is a much stronger constraint that may not reflect the true complexity of biological systems.

#### D.2. Limitations of Linear Methods

Applying linear dimensionality reduction techniques such as PCA/SVD constructs a flat manifold of dimension $d$ to approximate our fitness data. Through SVD, our $N \times M$ data matrix $\underline{\underline{D}}$ ($N$ being the number of environments and $M$ being the number of genotypes) containing fitness profiles is factorized as

$$\underline{\underline{D}} = \underline{\underline{U}}\,\underline{\underline{\Sigma}}\,\underline{\underline{V}}^T. \tag{S98}$$

The coordinates of genotype $i$ in this linear manifold are given by

$$\underline{z}_i = \begin{bmatrix} \sigma_1 v_{i1} \\ \sigma_2 v_{i2} \\ \vdots \\ \sigma_d v_{id} \end{bmatrix}. \tag{S99}$$

The key property of this linear approach is that Euclidean distances in the manifold directly correspond to differences in fitness profiles in the truncated space

$$\|\underline{z}_1 - \underline{z}_2\| = \|\underline{\hat{f}}_1 - \underline{\hat{f}}_2\|. \tag{S100}$$

While mathematically elegant and computationally tractable, this linear constraint may limit our ability to capture the true structure of the underlying phenotypic space efficiently.

##### D.3. Mathematical Advantages of Nonlinear Manifolds

A critical insight comes from the Whitney Embedding Theorem, which states that any smooth $d'$-dimensional manifold can be embedded in a Euclidean space of dimension $2d'+1$. This theorem explains the "unreasonable effectiveness of linear maps" while simultaneously highlighting their limitations. If the true causal manifold has intrinsic dimension $d'$ but is nonlinear, capturing its structure with a linear approximation generally requires $d > d'$ dimensions.

To formalize this intuition, we consider the causal manifold as a Riemannian manifold with a position-dependent metric tensor $\underline{\underline{G}}(\underline{z})$. On a nonlinear manifold, this tensor is not the identity matrix, meaning

$$\|\underline{z}_1 - \underline{z}_2\| \neq \|\underline{\hat{f}}_1 - \underline{\hat{f}}_2\|. \tag{S101}$$

Instead, the distance between points on a Riemannian manifold is given by

$$d_{\mathcal{M}_c}(\underline{z}_1, \underline{z}_2) = \min_{\underline{\gamma}} \int_0^1 dt \sqrt{\underline{\dot{\gamma}}(t)^T \, \underline{\underline{G}}(\underline{\gamma}(t)) \, \underline{\dot{\gamma}}(t)}, \tag{S102}$$

where $\underline{\gamma}$ is a parametric curve with $\underline{\gamma}(0) = \underline{z}_1$ and $\underline{\gamma}(1) = \underline{z}_2$. The curve that minimizes this distance is called a **geodesic**.

The key insight is that when the true causal manifold is nonlinear, approximating it with a linear method requires additional dimensions to compensate for the curvature. This is reflected in the singular value spectrum, where

$$\|\underline{\hat{f}}_1 - \underline{\hat{f}}_2\|^2 = \sum_{j=1}^{d} \sigma_j^2 (v_{1j} - v_{2j})^2. \tag{S103}$$

In this case, dimensions beyond the intrinsic dimensionality $d'$ (with smaller singular values) are effectively "compensating" for the curvature of the true manifold.

##### D.4. Nonlinear Methods for Manifold Learning

To learn nonlinear manifolds directly, we employ variational autoencoders (VAEs) that can approximate the joint distribution between fitness profiles $\underline{f}$ and latent variables $\underline{z}$ representing coordinates on the causal manifold

$$\pi(\underline{f}, \underline{z}) \approx \pi(\underline{f}|\underline{z})\pi(\underline{z}). \tag{S104}$$

VAEs employ two key neural networks

1. An encoder function that approximates the posterior distribution:

$$\Phi^{(E)} : \underline{f} \in \mathbb{R}^N \to \underline{\begin{bmatrix} \underline{\mu}^{(E)} \\ \log \underline{\sigma}^{(E)} \end{bmatrix}} \in \mathbb{R}^{2d}. \tag{S105}$$

2. A decoder function that maps latent points to fitness profiles:

$$\Psi^{(D)} : \underline{z} \in \mathcal{M}_c \to \underline{\mu}^{(D)} \in \mathbb{R}^N. \tag{S106}$$

For Riemannian Hamiltonian Variational Autoencoders (RHVAEs), used throughout this work, we add a third network that explicitly learns the metric tensor $\underline{\underline{G}}(\underline{z})$, allowing the model to capture the geometric structure of the manifold directly.

##### D.5. Comparative Advantages of Nonlinear Methods

Nonlinear methods offer several theoretical advantages over linear approaches:

1. **Dimensionality Efficiency**: Nonlinear methods can potentially represent the same information with fewer dimensions. If the true causal manifold has intrinsic dimension $d'$ but is nonlinear, linear methods may require $d > d'$ dimensions to achieve comparable accuracy.
2. **Geometric Fidelity**: By learning the metric tensor directly, nonlinear methods can capture the true geometric structure of the fitness landscape, including features like multiple peaks, valleys, and saddle points that may be difficult to represent in a linear space.
3. **Predictive Power**: If the true relationship between genotype and fitness is nonlinear, nonlinear methods should provide better predictive accuracy, especially for genotypes distant from those in the training set.

4. **Information Compression**: By capturing the curvature of the manifold explicitly rather than through additional dimensions, nonlinear methods can provide a more compact and interpretable representation of the fitness landscape.

##### D.6. Empirical Evidence and Caveats

While the theoretical advantages of nonlinear methods are clear, empirical validation is essential. We observe this advantage in practice through several metrics:

1. **Reconstruction Error**: Nonlinear methods achieve lower reconstruction error for the same latent dimension compared to linear methods.
2. **Singular Value Spectrum**: The slow decay of singular values in the data matrix suggests inherent nonlinearity in the fitness landscape.
3. **Generalization Performance**: Nonlinear methods show better performance when predicting fitness for held-out genotypes.

However, it's important to note that nonlinear methods come with their own challenges:

1. **Overfitting Risk**: Nonlinear methods may introduce spurious complexity when the true structure is simple.
2. **Interpretation Challenges**: The biological meaning of nonlinear latent dimensions can be less straightforward than those from linear methods.
3. **Training Complexity**: Nonlinear methods typically require more data and computational resources for effective training.
4. **Validation Requirements**: Demonstrating that a learned nonlinear structure reflects true biological relationships rather than mathematical artifacts requires careful validation.

In this work, we address these challenges through rigorous cross-validation, comparison with linear baselines, and direct analysis of the learned metric structure to ensure biological relevance of our nonlinear representations.

#### E. Cross-Validation Methodology for Evaluating Latent Space Predictive Power

To rigorously evaluate whether nonlinear latent space coordinates capture more information about the underlying phenotypic state than linear projections, we implemented a cross-validation framework that tests each model's ability to predict responses to unseen antibiotics. This approach extends the bi-cross validation methodology described by Kinsler, Geiler-Samerotte, and Petrov [11], adapting it for both linear (PCA/SVD) and nonlinear (VAE and RHVAE) dimensionality reduction techniques.

#### E.1. Theoretical Background

The core insight of our cross-validation approach is that if latent space coordinates genuinely capture the underlying phenotypic state of the system, they should enable accurate prediction of cellular responses to antibiotics not used in training. We test this hypothesis by systematically holding out one antibiotic at a time, learning latent space coordinates without information from that antibiotic, and then evaluating how well these coordinates predict the response to the held-out antibiotic.

#### E.2. Linear Bi-Cross Validation (SVD)

For linear models, we implement the bi-cross validation method of Kinsler, Geiler-Samerotte, and Petrov [11]. Given our $IC_{50}$ data matrix $\underline{\underline{X}} \in \mathbb{R}^{m \times n}$ where rows represent antibiotics and columns represent genotypes, we partition the matrix into four quadrants

$$\underline{\underline{X}} = \begin{bmatrix} \underline{\underline{A}} & \underline{\underline{B}} \\ \underline{\underline{C}} & \underline{\underline{D}} \end{bmatrix} \tag{S107}$$

Here, $\underline{\underline{A}}$ represents the target quadrant containing the held-out antibiotic-genotype combinations we aim to predict. Specifically, for a given held-out antibiotic $i$:

- $\underline{\underline{A}} \in \mathbb{R}^{1 \times k}$ contains $IC_{50}$ values for antibiotic $i$ and the validation set genotypes ($k$ genotypes)
- $\underline{\underline{B}} \in \mathbb{R}^{1 \times (n-k)}$ contains $IC_{50}$ values for antibiotic $i$ and the training set genotypes
- $\underline{\underline{C}} \in \mathbb{R}^{(m-1) \times k}$ contains $IC_{50}$ values for all other antibiotics and the validation set genotypes
- $\underline{\underline{D}} \in \mathbb{R}^{(m-1) \times (n-k)}$ contains $IC_{50}$ values for all other antibiotics and the training set genotypes

The theoretical foundation of this approach rests on the observation that for a full-rank matrix, we can estimate $\underline{\underline{A}}$ as

$$\underline{\underline{A}} = \underline{\underline{B}}\,\underline{\underline{D}}^{+}\underline{\underline{C}} \tag{S108}$$

where $\underline{\underline{D}}^{+}$ is the Moore-Penrose pseudoinverse of $\underline{\underline{D}}$. To evaluate how well a low-rank approximation performs, we compute rank-$r$ approximations of $\underline{\underline{D}}$ using SVD:

1. Perform SVD on $\underline{\underline{D}}$:

$$\underline{\underline{D}} = \underline{\underline{U}}\,\underline{\underline{\Sigma}}\,\underline{\underline{V}}^{T} \tag{S109}$$

2. Create rank-$r$ approximation by retaining only the top $r$ singular values:

$$\underline{\underline{D}}_r = \underline{\underline{U}}\,\underline{\underline{\Sigma}}_r\,\underline{\underline{V}}^T \tag{S110}$$

3. Compute the predicted matrix:

$$\hat{\underline{\underline{A}}}_r = \underline{\underline{B}}\,\underline{\underline{D}}_r^+\,\underline{\underline{C}} \tag{S111}$$

4. Calculate the mean squared error (MSE) between $\underline{\underline{A}}$ and $\hat{\underline{\underline{A}}}_r$:

$$\text{MSE}_r = \frac{1}{|\underline{\underline{A}}|}\sum_{i,j}(\underline{\underline{A}}_{ij} - \hat{\underline{\underline{A}}}_{r,ij})^2 \tag{S112}$$

By examining how MSE varies with rank, we can determine the minimum dimensionality needed to accurately predict the held-out antibiotic data.

#### E.3. Non-Linear Cross-Validation (VAE and RHVAE)

For nonlinear models (VAE and RHVAE), we adapt the cross-validation approach to account for the different model architecture. The procedure for each held-out antibiotic $i$ consists of two phases:

##### E.3.1. Phase 1: Training the Encoder

First, we train a complete model (encoder + decoder) on $IC_{50}$ data from all antibiotics except the held-out antibiotic $i$. This yields an 85%-15% train-validation split of genotypes for robust training. Denoting the set of all antibiotics as $\mathcal{A}$ and the held-out antibiotic as $a_i$, we train on the data matrix $\underline{\underline{X}}_{\mathcal{A}\setminus\{a_i\}}$.

For the VAE, we maximize the evidence lower bound (ELBO):

$$\mathcal{L}(\theta, \phi) = \left\langle \log p_\theta(\underline{x}|\underline{z}) \right\rangle_{q_\phi(\underline{z}|\underline{x})} - D_{KL}(q_\phi(\underline{z}|\underline{x})||\pi(\underline{z})) \tag{S113}$$

where $\underline{x}$ represents the $\log IC_{50}$ values for a genotype across all antibiotics except $a_i$, $\underline{z}$ is the latent representation, $q_\phi(\underline{z}|\underline{x})$ is the encoder, and $p_\theta(\underline{x}|\underline{z})$ is the decoder.

For the RHVAE, we similarly maximize the ELBO but with modifications to account for the Riemannian geometry of the latent space. The model learns a position-dependent metric tensor $\underline{\underline{G}}(\underline{z})$ along with the encoder and decoder parameters.

After training, the encoder maps inputs $\underline{x}$ to latent coordinates $\underline{z}$ that capture the underlying phenotypic state without information from antibiotic $a_i$.

###### E.3.2. Phase 2: Training the Missing Antibiotic Decoder

Once the encoder is trained, we freeze its parameters—effectively fixing the latent space coordinates for each genotype. We then train a decoder-only model to predict the response to the held-out antibiotic $a_i$ using a 50%-50% train-validation split of genotypes

$$\underline{z} = \text{Encoder}_\phi(\underline{x}) \tag{S114}$$

$$\hat{y}_i = \text{Decoder}_\theta(\underline{z}) \tag{S115}$$

where $\hat{y}_i$ is the predicted $IC_{50}$ value for antibiotic $a_i$.

This training maximizes the likelihood of the observed $IC_{50}$ values given the latent coordinates:

$$\mathcal{L}_{\text{decoder}} = \left\langle \log p_\theta(y_i|\underline{z}) \right\rangle_{q_\phi(\underline{z}|\underline{x})} \tag{S116}$$

For both VAE and RHVAE, we use a 2-dimensional latent space to ensure fair comparison with linear methods. After training, we evaluate the model's predictive performance on the validation set by computing the mean squared error between predicted and actual $IC_{50}$ values.

##### E.4. Implementation Details

The specific implementation involved several key components:

1. **Data Organization**: The $IC_{50}$ values were organized in a 3D tensor with dimensions [antibiotics $\times$ genotypes $\times$ MCMC samples], where the MCMC samples represent posterior samples from the Bayesian inference procedure used to estimate $IC_{50}$ values in Iwasawa et al. [1].

2. **Training Protocol**: For each held-out antibiotic:

   - The full model (encoder + decoder) was trained for the linear mapping from all antibiotics except the held-out one to the latent space.
   - The encoder parameters were frozen, and a new decoder was trained to map from the latent space to the held-out antibiotic's $IC_{50}$ values.
   - A tempering parameter $\beta_0$ was set to 0.3 to improve stability during training.
   - Training used the Adam optimizer with a learning rate of $10^{-3}$.
   - Models were trained for 50 epochs with a batch size of 256.

3. **Model Architecture**:

- The encoder consisted of a joint logarithmic encoder that maps inputs to latent space.
- The decoder was a simple network mapping from latent space to antibiotic responses.
- For RHVAE, the model incorporated a metric chain with position-dependent Riemannian metric.

4. **Evaluation Metrics**: We assessed predictive performance using mean squared error (MSE) between actual and predicted $IC_{50}$ values on the validation set.

##### E.5. Comparative Analysis

To fairly compare the predictive power of linear and nonlinear dimensionality reduction techniques, we plotted the MSE for SVD at different ranks alongside the MSE for 2D VAE and 2D RHVAE models. This visualization, shown in Figure 7 in the main text, demonstrates that for all antibiotics, the 2D nonlinear latent space coordinates provide more accurate predictions than any number of linear dimensions.

The key finding is that the nonlinear latent spaces capture the underlying phenotypic structure of the system more effectively than linear projections, thus enabling better predictions of out-of-sample data. This validates our hypothesis that the phenotypic space of antibiotic resistance has an inherently nonlinear structure that is better represented by VAE and especially RHVAE models.

##### E.6. Cross-Validation comparison of 2D and 3D latent spaces

In Figure 7 in the main text, we tested the predictive power of 2D nonlinear latent space coordinates compared to linear latent space coordinates. There, we showed via a custom cross-validation scheme that the 2D latent space coordinates were more predictive of out-of-sample data than the linear latent space coordinates. Here, we repeat a similar analysis, but using 3D latent space coordinates.

Following the same cross-validation scheme as in the main text, we trained a full model on all but one antibiotic (85%-15% splits for training and validation data). This generates the latent space coordinates without using any information from the antibiotic that was left out. Next, we froze the encoder parameters —equivalent to fixing the latent space coordinates— and trained a decoder-only model on the missing antibiotic data (50%-50% splits for training and validation data). Figure E shows the results of this analysis. We can see that other than a couple of exceptions (KM and NFLX), the 3D latent space coordinates are marginally more predictive of out-of-sample data than the 2D latent space coordinates. This suggests that the phenotypic changes associated with the experimental setup can be captured with two effective degrees of freedom.

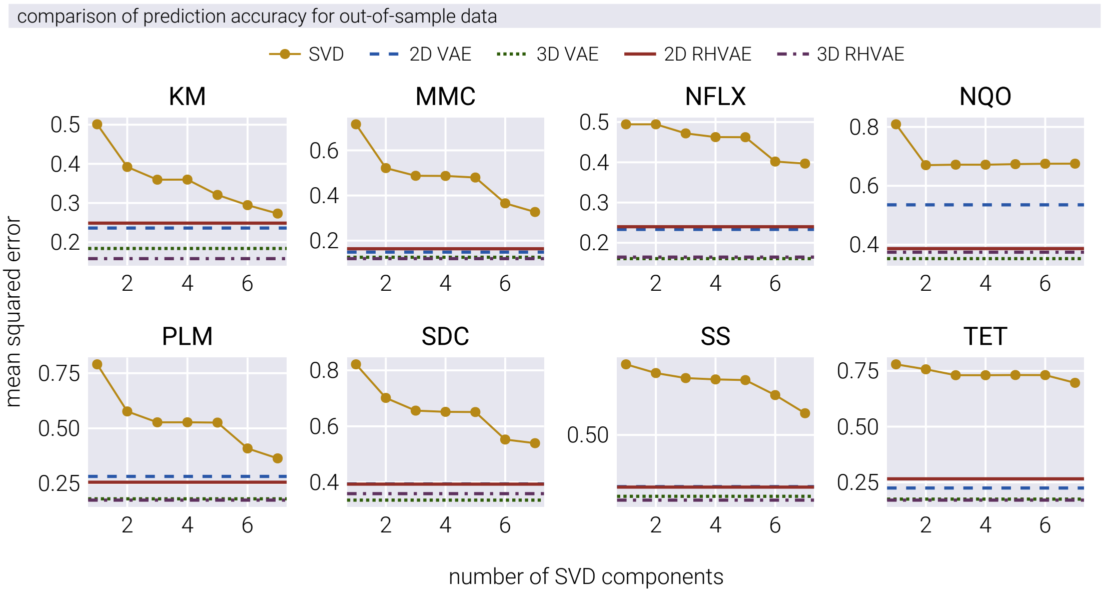

**Figure E. Comparison of 2D and 3D nonlinear latent space models for predicting out-of-sample antibiotic data**. Reconstruction error for each missing antibiotic as a function of linear dimensions used in SVD cross-validation. Horizontal lines represent the accuracy of nonlinear models: 2D-VAE (dark blue, dashed), 2D-RHVAE (dark red, solid), 3D-VAE (dark green, dotted), and 3D-RHVAE (dark purple, dash-dotted). The 3D models show marginal improvement over their 2D counterparts for most antibiotics, suggesting that two effective degrees of freedom capture most of the relevant phenotypic variation.

#### F. Geodesics in Latent Space

In this section, we explore an interesting and surprising property of some of the resulting evolutionary trajectories in latent space. As detailed in the main text, we trained an RHVAE on the $IC_{50}$ data from [1]. The data used to train the RHVAE consists of a matrix with eight rows—one per each antibiotic—and $\approx$ 1300 columns representing one of the lineages at some time point in the experiment. However, at no point, the RHVAE has any knowledge of what genotype or time point corresponds to any particular column in the data matrix. All it sees at every epoch is a random subset of columns of this matrix. Nevertheless, it is interesting to ask whether there is some regularity in the resulting evolutionary trajectories in the learned latent space. In other words, once the RHVAE is trained, we can map the time series corresponding to a particular lineage to the corresponding latent space trajectory. A few of these trajectories are shown in Figure F as gold connected points.

One of the unique features of the RHVAE model is the co-learning of the metric tensor in the latent space. A space endowed with such mathematical structure gives us the ability to compute the shortest path between any two points, commonly referred to as a geodesic. Geodesics are the generalization of the idea of a straight line in Euclidean space to curved spaces, and it is a fundamental concept in Riemannian geometry. Let us define this curve as $\underline{\gamma}(t)$ where $t \in [0, 1]$. This means that the geodesic is a parametric curve $\underline{\gamma}$ for which we only need to define the initial and final points of the curve, $\underline{\gamma}(0) = \underline{z}_0$ and $\underline{\gamma}(1) = \underline{z}_1$. What makes it a geodesic is that it minimizes the length of the curve connecting the initial and final points, i.e., it minimizes

$$L(\underline{\gamma}) = \int_0^1 dt \sqrt{\dot{\underline{\gamma}}(t)^T \underline{\underline{G}}(\underline{\gamma}(t)) \dot{\underline{\gamma}}(t)} \tag{S117}$$

where $\dot{\underline{\gamma}}(t) = \frac{d}{dt}\underline{\gamma}(t)$ is the velocity vector of the curve and $\underline{\underline{G}}(\underline{\gamma}(t))$ is the metric tensor at the point $\underline{\gamma}(t)$. Computing this geodesic is a non-trivial task, especially for a data-derived metric tensor with no closed-form expression. Fortunately, we can leverage the generality of neural networks to parameterize the curve and compute the geodesic numerically [12]. Again, all we need to do is define the initial and final points of the curve, provide the metric tensor learned by the RHVAE, and let the neural network approximate the shortest path between the two points. Figure F shows the corresponding curves between the initial point (black crosses) and the final point (black triangles) as red curves for a few lineages. We highlight the fact that the curves are not straight lines in the latent space, but rather curved paths that follow the curvature of the latent space. For these few examples, we see that the geodesic curves are very similar to the gold curves, which are the actual trajectories of the lineages in the latent space. This rather surprising result suggests that the best spatial representation for the evolutionary trajectories coincides with the shortest path in the latent space.

So far, these few examples suggest that the shortest path in the latent space coincides with the actual evolutionary trajectory. Since the latent space coordinates of any data point capture information about the resistance profile of the corresponding strain, we expect that

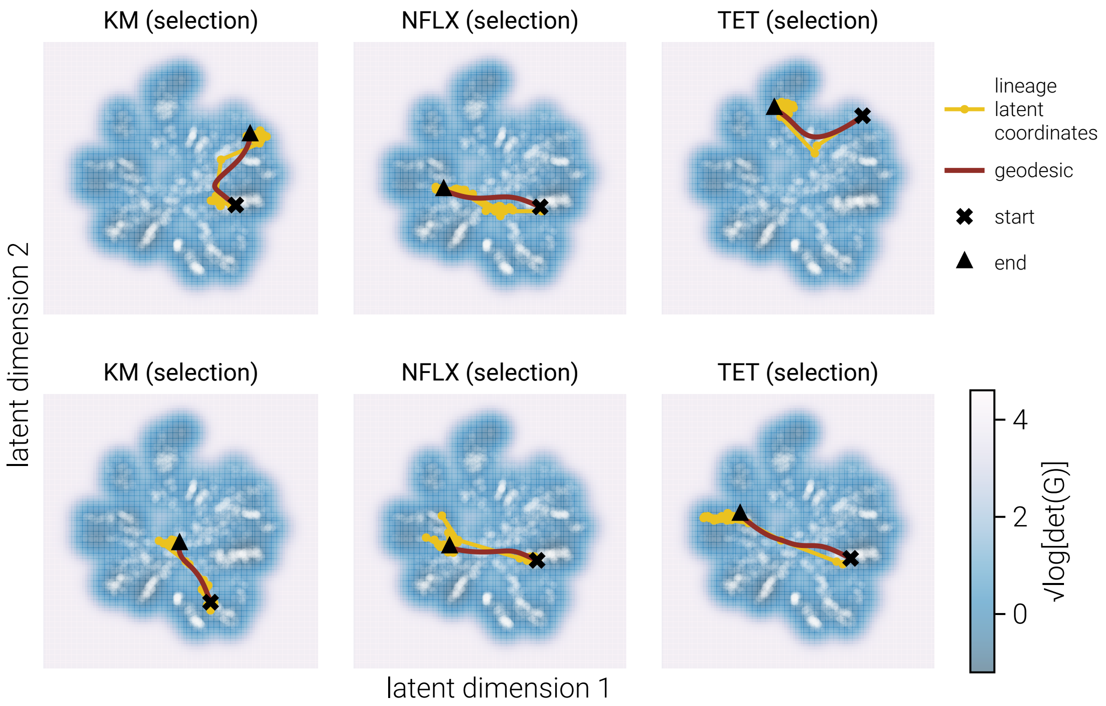

**Figure F. Geodesic paths in latent space follow evolutionary trajectories** The figure shows the latent space representation of evolutionary trajectories from the Iwasawa et al. dataset [1]. Background coloring represents the metric volume (determinant of the metric tensor), with lighter regions indicating higher local curvature. Gold connected points show actual evolutionary trajectories of different lineages, while red curves represent the computed geodesics (shortest paths) between initial points (black crosses) and final points (black triangles). The striking similarity between geodesics and actual trajectories suggests that evolution in phenotype space tends to follow paths that minimize distance in the geometry-informed latent space.

the qualitative matching between the geodesic and the actual trajectory should mean that the resulting resistance profile predicted by the geodesic is very similar to the actual experimental resistance profile. To see if this is the case, we must decode the latent space trajectory back to the original space. Doing so projects back all of the points sampled along the geodesic curve to the original data space with eight antibiotic resistance values. However, there is a catch: the geodesic curve is a smooth continuous function of time, but the resistance profile of a strain is a discrete set of values with different step sizes. In other words, when constructing the geodesic curve, we define a continuous function connecting the initial and final point with no sudden jumps in the coordinates. But there is no *a priori* reason that the resulting experimental curve should be smooth. This statement is not necessarily a consequence of the discrete sampling of the data with a one-day time step that could be resolved by increasing the sampling frequency, but rather a consequence of the genotype-phenotype map. Although in the simulations presented in the main text we assume that all evolutionary steps follow certain distribution of step sizes as a convenient mathematical simplification, the complexity of the genotype-phenotype map might be such that this simplification is not valid. To emphasize this point even further, we can think of the geodesic curve as a prediction of what the cross-resistance values for the different antibiotics ought to be without information about when those values must evolve in time.

The consequence of this is that it is not clear how to align the continuous geodesic curve with the discrete experimental data. We can, however, make use of methodologies developed for similar purposes in the form of time-warping algorithms. In this case, we can use dynamic time warping (DTW) to align the continuous geodesic curve with the discrete experimental data. DTW is a well-known algorithm in the signal processing community for aligning two time series with different step sizes. Effectively, DTW finds the optimal pairing of points between the two time series, with the constraint that the alignment is monotonic, i.e., if point $x_i$ in one time series is aligned to point $y_j$ in the other time series, then $x_{i+1}$ is either aligned to $y_j$ or $y_{j+1}$, never stepping backwards. We use this algorithm to align the points sampled along the geodesic curve in latent space with the resulting experimental curve also in latent space. This point, although subtle, is important: the alignment is done in latent space, not in the original data space. This means that the alignment is done based on the metric tensor learned by the RHVAE, and not on the original data.

Once we have the aligned latent space trajectory, we can decode it back to the original data space and plot the resulting experimental curve. Figure G and Figure H show the results of this procedure for the same few lineages shown in Figure F. On the left, we show the same latent space trajectory as in Figure F, on the right, we show the resulting $IC_{50}$ curves for each of the eight antibiotics. The gold curves show the original experimental curves, while the red curves show the geodesic-predicted curves after the dynamic time warping alignment. Although far from perfect, the alignment is remarkably good considering that the predictions were drawn from only knowing the initial and final points of the trajectory in latent space and then computing the shortest path between them. This surprising result begs for further experimental investigation along with extensive theoretical analysis of why this phenomenon occurs. One tantalizing, although highly speculative, possibility is that evolution proceeds in phenotype space following a least action principle, i.e., the path taken by the evolutionary trajectory is the one that minimizes the distance traveled in phenotype space subject to the constraints of phenotype accessibility, as modeled by our genotype-phenotype density function in the main text.

### G. Geometry-Informed Latent Space Reconstruction of Antibiotic Resistance Landscapes

In the main text, Figure 5 shows the ability of the RHVAE to reconstruct the underlying fitness landscapes' topography from simulated data. There, we demonstrated that the relative position, number of peaks, and overall shape of the fitness landscapes are well-reproduced by the RHVAE latent space. It is therefore interesting to show the same reconstructions using the experimental data from [1].

After training an RHVAE model on the $IC_{50}$ data from [1]—the same model used in the main text in Figure 6—we apply the same methodology to reconstruct the fitness landscapes for the eight antibiotics used in the experiment. Figure I shows the reconstructed fitness landscapes for all eight antibiotics along with the latent space metric volume. One obvious difference between the landscapes reconstructed from the simulated and experimental data is the lack of clear peaks in the experimental landscapes. Although it is possible that such peaks do not exist, it is more likely that the lack of peaks is due to the limited resolution of the experimental data, where only a few initial genotypes were measured.

Given the lack of regularity in the resulting landscapes, we also show the three-dimensional representations of the resistance landscapes for different antibiotics using experimental data from [1]. Figure J shows the same landscapes as in Figure I, but using the z-axis to represent the fitness value. From this 3D perspective, the lack of clear peaks is more apparent.

#### G.1. Trajectories in Experimentally-reconstructed Fitness Landscapes

Given these reconstructed fitness landscapes, we can now explore the evolutionary trajectories of different lineages in the experimental data. In [1], the authors evolved genotypes in the presence of three different antibiotics: *Tetracycline* (TET), *Kanamycin* (KAN), and *Norfloxacin* (NFLX). Naively, if these fitness landscapes are representative of the underlying landscape available to the genotypes, we would expect the trajectories of these lineages to follow a noisy gradient ascent-like path in the latent space, following the fitness gradient predicted by the reconstructed landscapes. Figure K, Figure L, and Figure M show the evolutionary trajectories of the lineages evolved in the presence of *Kanamycin*, *Tetracycline*, and *Norfloxacin*, respectively. The left and right panels show the same trajectories in the 2D and 3D projection of the latent space, respectively. One of the problematic aspects of these trajectories is that in none of the cases do the trajectories follow a path towards the predicted highest fitness peak. We strongly suspect this is a problem with the experimental setup, rather than the entirety of our approach. Our suspicion is that the selection of initial genotypes completely biased the reconstruction of the fitness landscapes. Some of

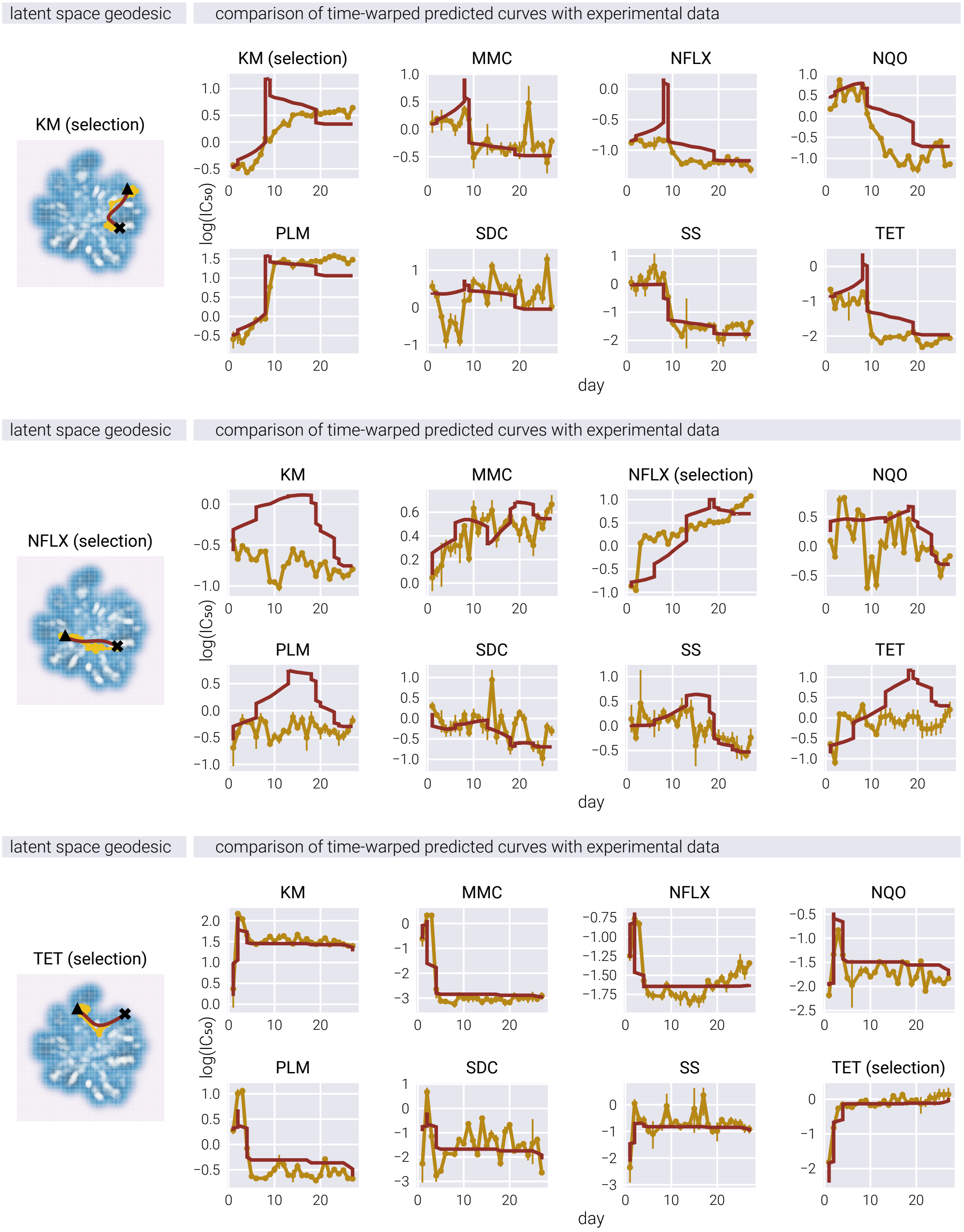

**Figure G. Geodesic predictions of evolutionary trajectories match experimental data after time alignment.** Left panel shows latent space trajectories (gold: experimental path; red: geodesic prediction) for selected lineages. Right panel shows the corresponding IC50 values for all eight antibiotics over time, comparing experimental measurements (gold) with geodesic predictions after dynamic time warping alignment (red).

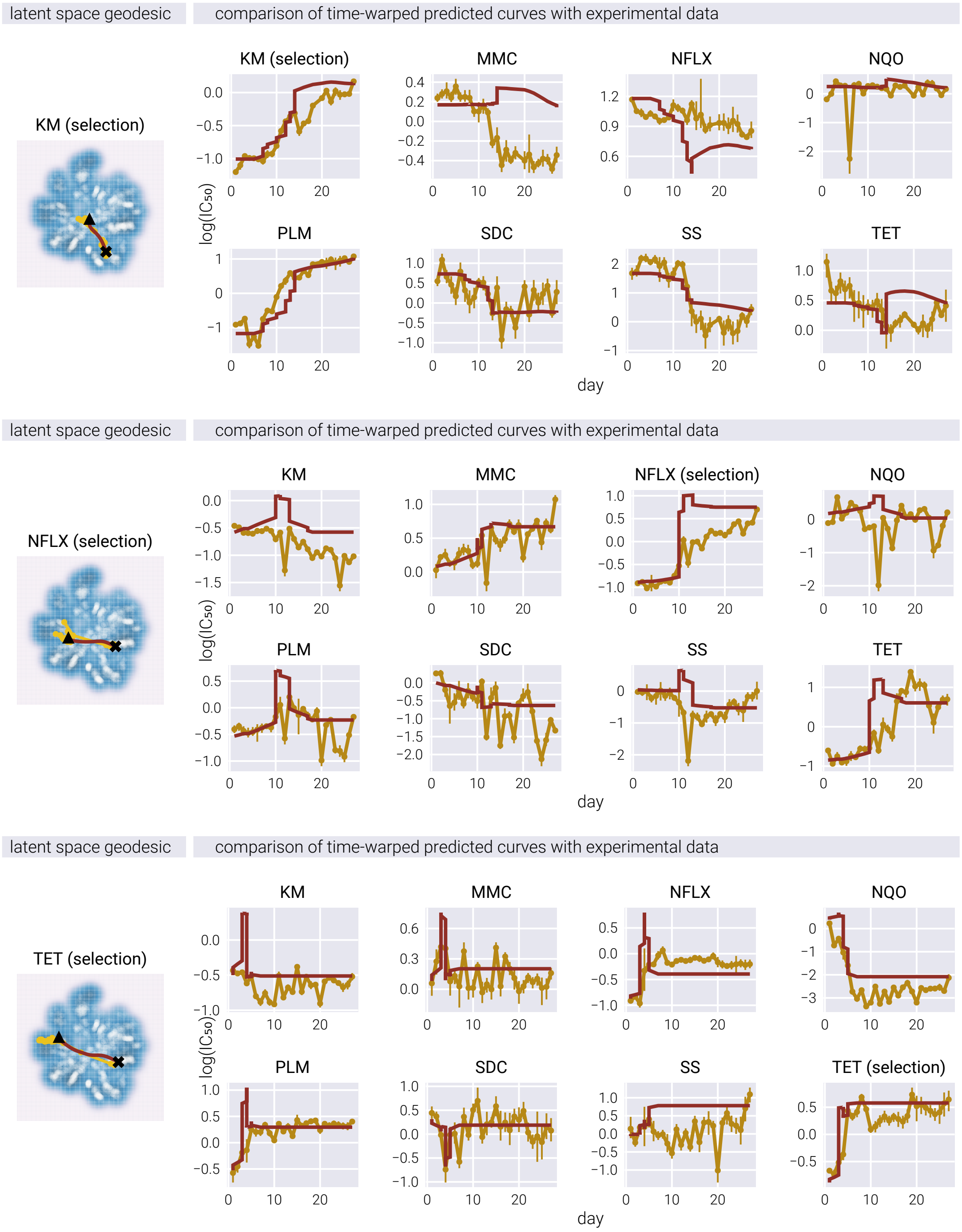

**Figure H. Additional examples of geodesic-based predictions of antibiotic resistance evolution.** Each panel pair shows another set of lineages where geodesic paths in latent space (left, red curves) were used to predict the temporal evolution of resistance profiles (right). Despite only using initial and final points as inputs, the geodesic predictions (red) capture many features of the actual evolutionary trajectories (gold) after time alignment.

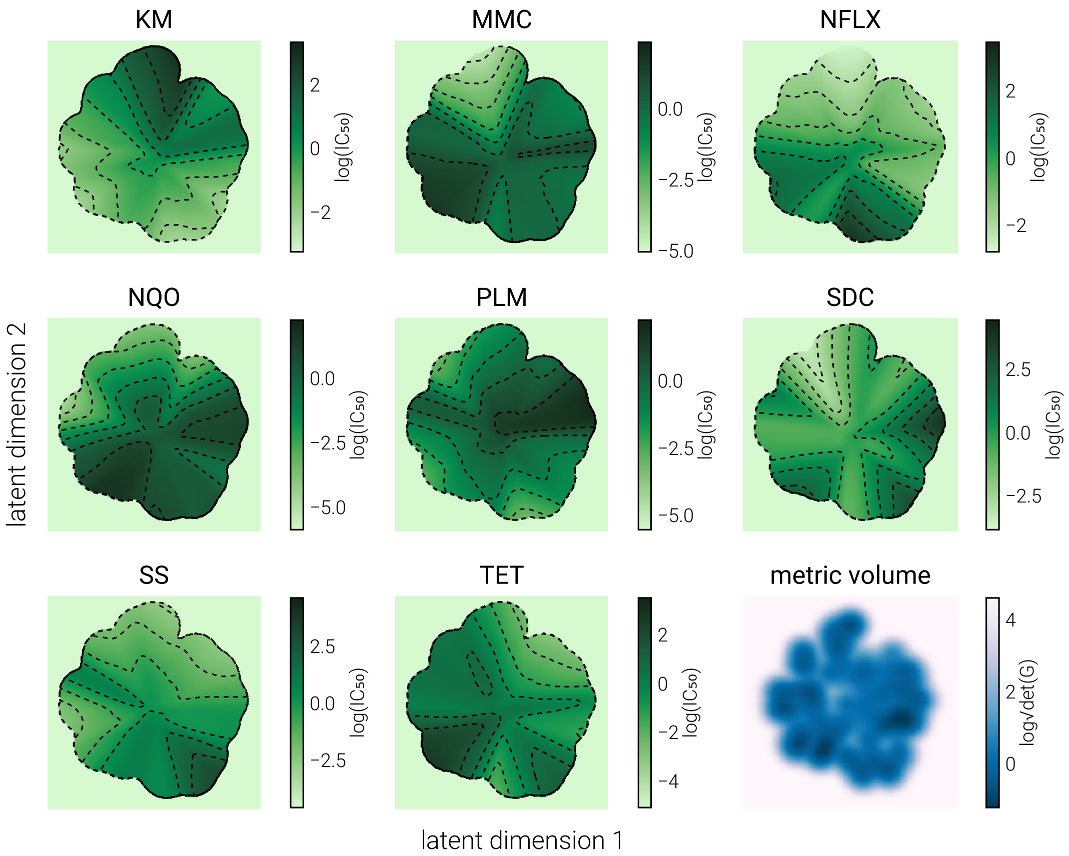

**Figure I. Reconstructed 2D antibiotic resistance landscapes from experimental data**. The first eight panels show the antibiotic resistance landscape reconstructions using a 2D RHVAE trained on the $IC_{50}$ data from [1]. Dotted lines show different level curves on the fitness landscape. The last panel shows the metric volume (the determinant of the metric tensor) of the latent space. The darker the color, the flatter the latent space.

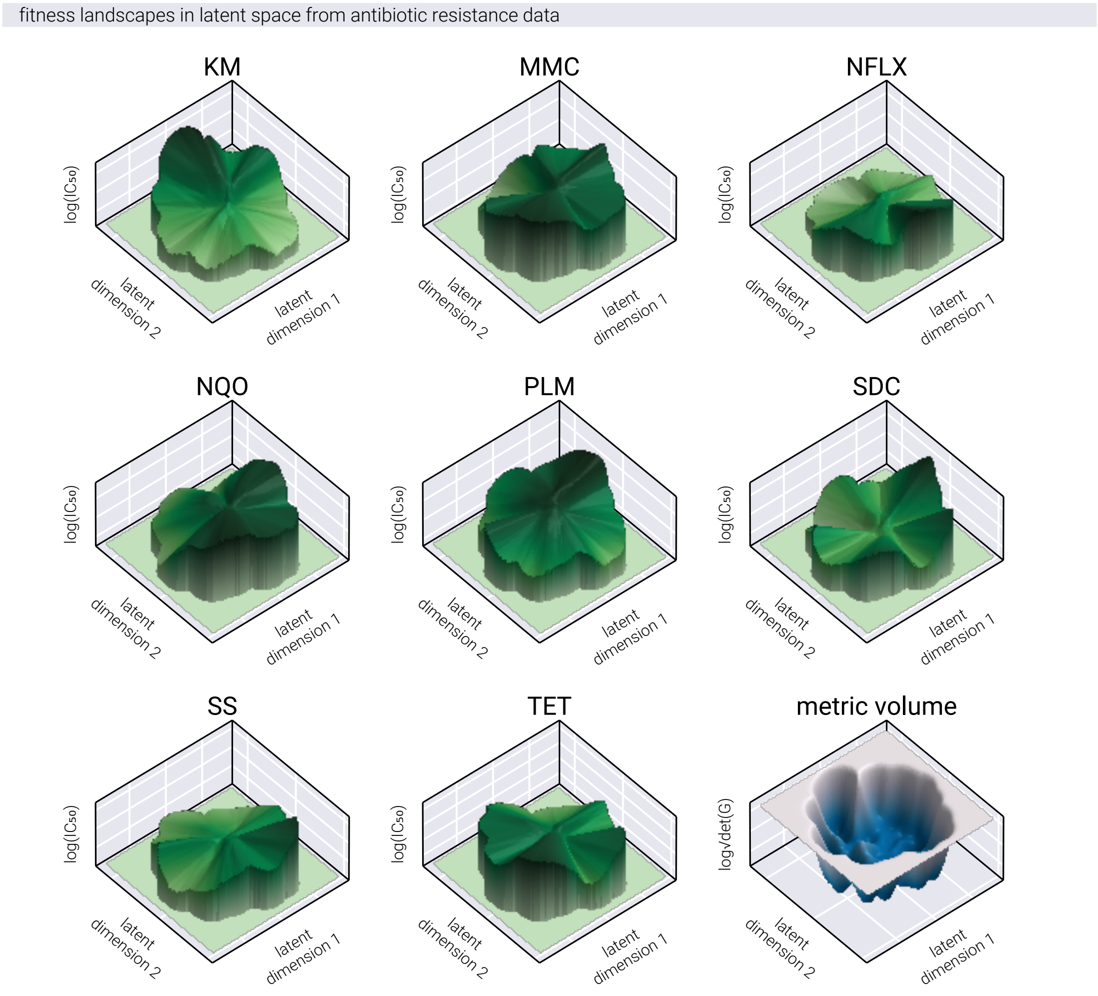

**Figure J. 3D visualization of reconstructed antibiotic resistance landscapes**. Three-dimensional representations of the same resistance landscapes shown in Figure I, using experimental data from [1]. The z-axis represents the fitness value (IC ). This 3D perspective more clearly illustrates the lack of distinct peaks in the experimental data, which may be due to limited sampling of initial genotypes rather than the absence of such features in the actual fitness landscapes.

the lineages selected by the authors were previously evolved in the presence of one of these antibiotics, to then be re-evolved in the presence of the other antibiotics. In other words, there were some strains in the initial pool that were already highly resistant to, say, *Kanamycin*, and then these strains were re-evolved in the presence of *Tetracycline*. The presence of these strains explains the existence of those peaks that none of the trajectories reach, since these strains evolved for much longer in the corresponding antibiotic compared to this experimental setup. However, further investigation is needed to determine the cause of this behavior.

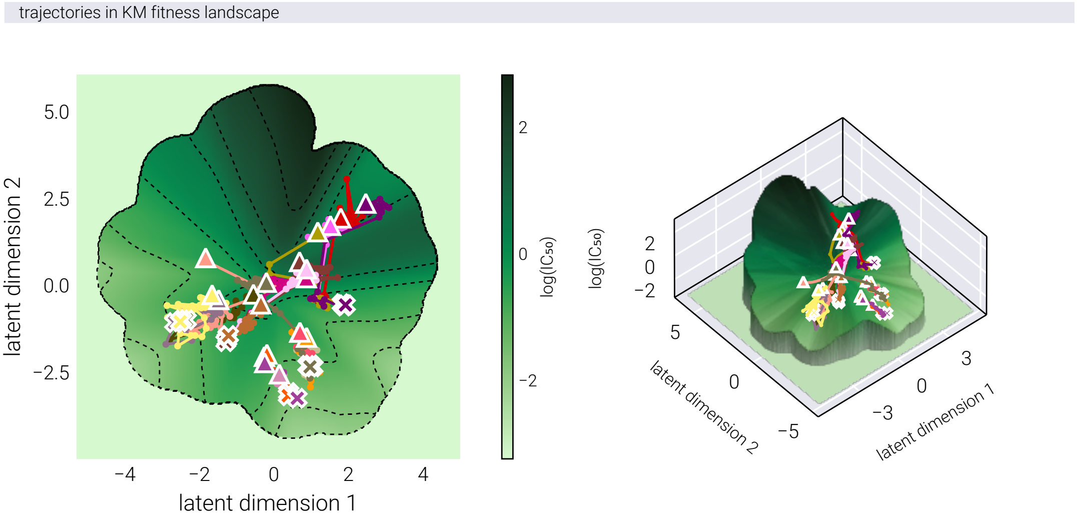

**Figure K. Evolutionary trajectories of lineages evolved in Kanamycin**. The left panel shows the 2D projection of evolutionary trajectories in the latent space overlaid on the Kanamycin resistance landscape. The right panel shows the same trajectories in a 3D projection. Different colors represent different lineages. Cross symbols indicate the initial point of the trajectory, while triangles indicate the final point. Note that trajectories do not follow paths toward the highest predicted fitness peaks, suggesting potential biases in the experimental setup or limitations in landscape reconstruction.

#### H. Neural Network Architecture

##### H.1. RHVAE Overview

The neural network architecture implemented in this study is a Riemannian Hamiltonian Variational Autoencoder (RHVAE) [3]. The network was implemented using the `Flux.jl` framework [13] via the `AutoEncoderToolkit.jl` package [14], and consists of three main components: an encoder, a decoder, and a metric chain network for the Riemannian metric computations.

##### H.2. Model Parameters

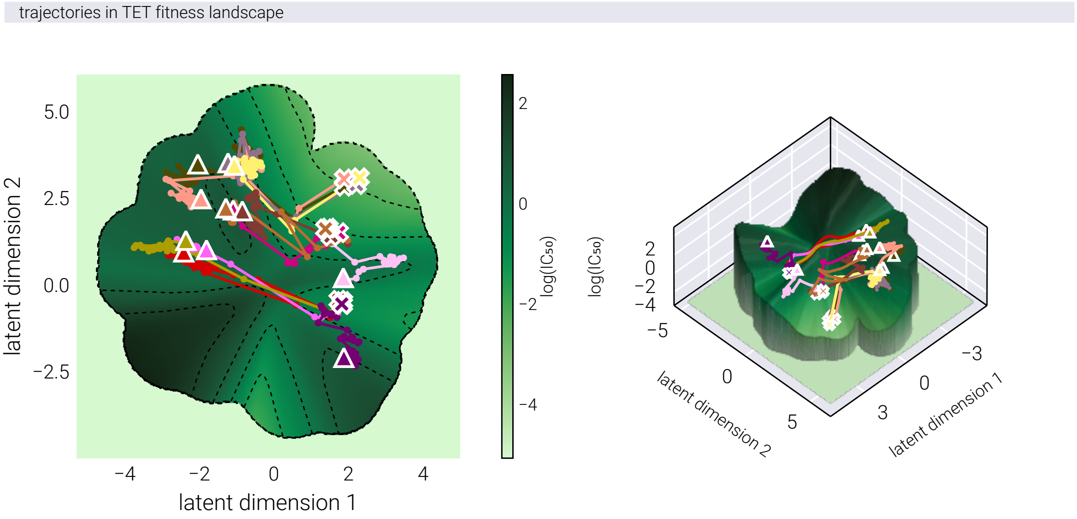

**Figure L. Evolutionary trajectories of lineages evolved in Tetracycline**. The left panel shows the 2D projection of evolutionary trajectories in the latent space overlaid on the Tetracycline resistance landscape. The right panel shows the same trajectories in a 3D projection. Different colors represent different lineages. Cross symbols indicate the initial point of the trajectory, while triangles indicate the final point. Similar to the Kanamycin case, trajectories do not consistently follow gradient ascent paths toward fitness peaks.

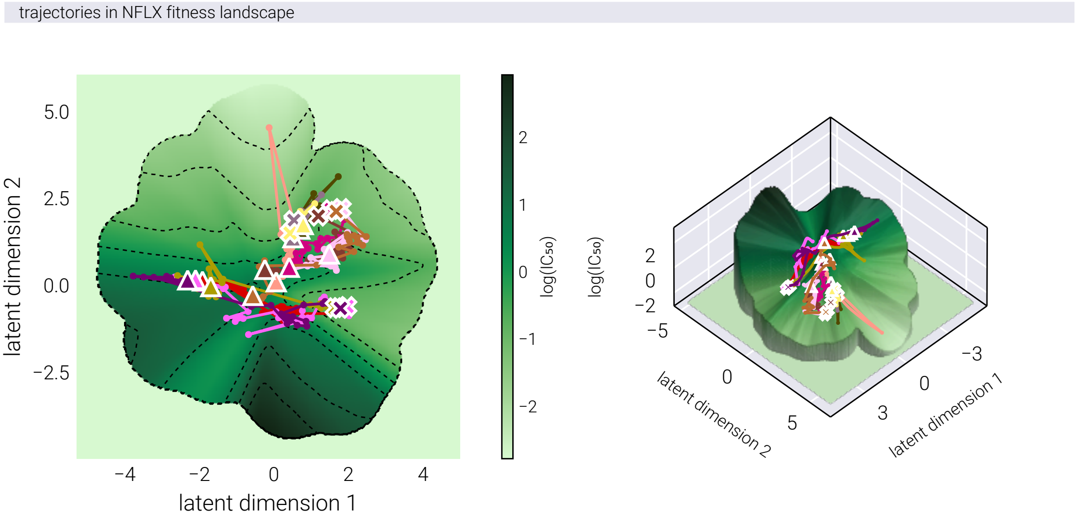

**Figure M. Evolutionary trajectories of lineages evolved in Norfloxacin**. The left panel shows the 2D projection of evolutionary trajectories in the latent space overlaid on the Norfloxacin resistance landscape. The right panel shows the same trajectories in a 3D projection. Different colors represent different lineages. Cross symbols indicate the initial point of the trajectory, while triangles indicate the final point. As with other antibiotics, the evolutionary paths do not align with the predicted fitness gradients, potentially due to pre-evolved strains in the initial pool biasing landscape reconstruction.

| Parameter | Value | Description |
|---|---|---|
| Latent Dimensions | 2 | Dimensionality of the latent space |
| Hidden Layer Size | 128 | Number of neurons in hidden layers |
| Temperature (T) | 0.8 | Temperature parameter for RHVAE |
| Regularization ( ) | 0.01 | Regularization parameter |
| Number of Centroids | 256 | Number of centroids for the manifold approximation |

#### H.3. Encoder Architecture

The encoder implements a joint Gaussian log-encoder with the following structure:

| Layer | Output Dimensions | Activation Function |
|---|---|---|
| Input | n_env* | |
| Dense 1 | 128 | Identity |
| Dense 2 | 128 | LeakyReLU |
| Dense 3 | 128 | LeakyReLU |
| Dense 4 | 128 | LeakyReLU |
| $\mu$ output | 2 | Identity |
| log output | 2 | Identity |

* n_env is the number of environments on which fitness is determined to train the model.

#### H.4. Decoder Architecture

The decoder implements a simple Gaussian decoder with the following structure:

| Layer | Output Dimensions | Activation Function |
|---|---|---|
| Input | 2 | |
| Dense 1 | 128 | Identity |
| Dense 2 | 128 | LeakyReLU |
| Dense 3 | 128 | LeakyReLU |
| Dense 4 | 128 | LeakyReLU |
| Output | n_env | Identity |

#### H.5. Metric Chain Architecture

The metric chain computes the Riemannian metric tensor with the following structure:

| Layer | Output Dimensions | Activation Function |
|---|---|---|
| Input | `n_env` | |
| Dense 1 | 128 | Identity |
| Dense 2 | 128 | LeakyReLU |
| Dense 3 | 128 | LeakyReLU |
| Dense 4 | 128 | LeakyReLU |
| Diagonal Output | 2 | Identity |
| Lower Triangular Output | 1 | Identity |

##### H.6. Data Preprocessing

The input data underwent the following preprocessing steps:

1. Logarithmic transformation of fitness values.
2. Z-score standardization (mean = 0, std = 1) per environment.
3. K-medoids clustering to select centroids for the manifold approximation.

##### H.7. Implementation Details

The model was implemented using:

- `Julia` programming language.
- `Flux.jl` for neural network architecture.
- `AutoEncoderToolkit.jl` for RHVAE-specific components.
- Random seed set to 42 for reproducibility.

The complete model state and architecture were saved in `JLD2` format for reproducibility and future use.

All of the code used to implement the model is available in the GitHub repository for this project.

#### I. Neural Geodesic Architecture

The neural network architecture implemented in this study is a Neural Geodesic model designed to find geodesics on the latent space manifold of a Riemannian Hamiltonian Variational Autoencoder (RHVAE) [12]. The network is implemented using the Flux.jl framework and learns to parameterize geodesic curves between points in the latent space.

##### I.1. Model Parameters

| Parameter | Value | Description |
|---|---|---|
| Hidden Layer Size | 32 | Number of neurons in hidden layers |
| Input Dimension | 1 | Time parameter t [0,1] |
| Output Dimension | 2 | Dimensionality of the latent space |
| Endpoints | z_init=[0,0], z_end=[1,1] | Start and end points of the geodesic |

##### I.2. Neural Geodesic Architecture

The neural geodesic implements a feed-forward network that maps from the time domain to the latent space with the following structure:

| Layer | Output Dimensions | Activation Function |
|---|---|---|
| Input | 1 | |
| Dense 1 | 32 | Identity |
| Dense 2 | 32 | TanH |
| Dense 3 | 32 | TanH |
| Dense 4 | 32 | TanH |
| Output | 2 | Identity |

##### I.3. Architectural Details

1. **Input Processing**
   - Takes a single time parameter $t \in [0, 1]$.
   - Maps to the latent space dimension (2D in this implementation).
   - Designed to learn smooth geodesic curves.
2. **Network Structure**
   - Follows a deep architecture with 4 hidden layers.
   - Uses `tanh` activation for smooth curve generation.
   - Identity activation at input and output layers for unrestricted range.
3. **Output Constraints**
   - Network outputs must satisfy boundary conditions:
     - $\gamma(0) = \underline{z}_{\text{init}}$.
     - $\gamma(1) = \underline{z}_{\text{end}}$.
   - Generated curve represents path in latent space.

##### I.4. Implementation Details

The model is implemented with:

- `Julia` programming language.
- `Flux.jl` for neural network architecture.
- `AutoEncode.diffgeo` package for geodesic-specific components.
- Random seed set to 42 for reproducibility.

The model is designed to work in conjunction with a pre-trained RHVAE model, using its latent space structure to inform the geodesic computation. The complete model state and architecture are saved in `JLD2` format for reproducibility and future use.

##### I.5. Integration with RHVAE

This neural geodesic model complements the RHVAE architecture by:

1. Using the same latent space dimensionality.
2. Learning geodesics that respect the Riemannian metric of the RHVAE
3. Providing a parameterized way to interpolate between latent points

#### J. Latent Space Alignment via Procrustes Analysis

When working with different dimensionality reduction techniques such as PCA, VAE, and RHVAE, the resulting latent spaces often have arbitrary orientations that make direct comparisons challenging. To facilitate meaningful comparisons between these latent representations and the ground truth phenotype space, we employed Procrustes analysis. This section details the mathematical foundations and implementation of our alignment procedure.

##### J.1. Mathematical Formulation

Procrustes analysis finds an optimal rigid transformation (rotation, scaling, and potentially translation) that aligns one set of points with another while minimizing the sum of squared differences. Given two sets of points $\underline{\underline{X}}$ and $\underline{\underline{Y}}$, where each column represents a point in $\mathbb{R}^d$, the objective is to find a transformation of $\underline{\underline{X}}$ that best aligns with $\underline{\underline{Y}}$.

The Procrustes problem can be formulated as:

$$\min_{R,s} \|\underline{\underline{Y}} - s\underline{\underline{R}}\,\underline{\underline{X}}\|_F^2 \tag{S118}$$

where:

- $\underline{\underline{R}}$ is a $d \times d$ orthogonal rotation matrix ($\underline{\underline{R}}^T \underline{\underline{R}} = \underline{\underline{I}}$)
- $s$ is a scalar scaling factor
- $\| \cdot \|_F$ denotes the Frobenius norm

If the data is centered (mean-subtracted), the transformation involves only rotation and scaling. The solution to this optimization problem is obtained through Singular Value Decomposition (SVD).

##### J.2. Algorithmic Procedure

Our implementation of Procrustes analysis follows these steps:

1. **Data Standardization**: Each latent representation (PCA, VAE, RHVAE) and the ground truth phenotype data are standardized to have zero mean and unit standard deviation along each dimension:

$$\hat{\underline{\underline{X}}} = \frac{\underline{\underline{X}} - \underline{\mu}_X}{\underline{\sigma}_X} \tag{S119}$$

where $\underline{\mu}_X$ and $\underline{\sigma}_X$ are the mean and standard deviation vectors computed across all points in the respective space.

2. **Inner Product Computation**: We compute the inner product matrix between the standardized ground truth phenotype data $\hat{\underline{\underline{Y}}}$ and each standardized latent representation $\hat{\underline{\underline{X}}}$:

$$\underline{\underline{A}} = \hat{\underline{\underline{Y}}} \hat{\underline{\underline{X}}}^T \tag{S120}$$

3. **Singular Value Decomposition**: We perform SVD on the inner product matrix:

$$\underline{\underline{A}} = \underline{\underline{U}} \underline{\underline{\Sigma}} \underline{\underline{V}}^T \tag{S121}$$

where $\underline{\underline{U}}$ and $\underline{\underline{V}}$ are orthogonal matrices, and $\underline{\underline{\Sigma}}$ is a diagonal matrix of singular values.

4. **Rotation Matrix Computation**: The optimal rotation matrix is given by:

$$\underline{\underline{R}} = \underline{\underline{U}} \underline{\underline{V}}^T \tag{S122}$$

5. **Scaling Factor Computation**: The optimal scaling factor is calculated as:

$$s = \frac{\text{tr}(\underline{\underline{\Sigma}})}{\|\underline{\underline{\hat{X}}}\|_F^2} \tag{S123}$$

where tr$(\underline{\underline{\Sigma}})$ is the trace of $\underline{\underline{\Sigma}}$ (sum of singular values) and $\|\underline{\underline{\hat{X}}}\|_F^2$ is the squared Frobenius norm of $\underline{\underline{\hat{X}}}$.

6. **Transformation Application**: Each latent representation is transformed using the corresponding rotation matrix:

$$\underline{\underline{X}}_{\text{aligned}} = \underline{\underline{R}}\,\underline{\underline{\hat{X}}} \tag{S124}$$

7. **Similarity Metric Calculation**: We compute a correlation-like measure to quantify the goodness-of-fit between the aligned spaces:

$$\rho = \frac{\text{tr}(\underline{\underline{\Sigma}})}{\|\underline{\underline{\hat{X}}}\|_F \|\underline{\underline{\hat{Y}}}\|_F} \tag{S125}$$

This metric ranges from 0 (no similarity) to 1 (perfect alignment).

##### J.3. Significance for Comparative Analysis

The Procrustes alignment enables several critical aspects of our analysis:

1. **Direct Visual Comparison**: By aligning all latent spaces to the same orientation as the ground truth phenotype space, we can directly visualize and compare the structural preservation properties of each dimensionality reduction technique.
2. **Quantitative Assessment**: The correlation metric $\rho$ provides a quantitative measure of how well each latent representation preserves the geometric structure of the ground truth phenotype space.
3. **Geometric Structure Preservation**: The alignment allows us to assess how features such as local neighborhoods, relative distances, and global structure are preserved in each latent representation.
4. **Trajectory Analysis**: For evolutionary trajectories, the alignment enables comparison of path lengths, directional changes, and convergence patterns across different representations.

This alignment procedure forms the foundation for the comparative analysis presented in the main text, where we evaluate the effectiveness of different dimensionality reduction techniques in capturing the underlying structure of the phenotype space.

### K. Alternative Metropolis-Kimura Evolutionary Dynamics

In the main text, we described a simple algorithm for evolving a population as a biased random walk on a phenotypic space controlled by both a fitness function and a genotype-phenotype density function. Here, we propose an alternative algorithm that combines elements of the Metropolis-Hastings algorithm with Kimura's classical population genetics theory.

#### K.1. Metropolis-Kimura Algorithm

In Section B, we described a simple framework with a fitness function and a genotype-phenotype density function. Here, we use the same framework but introduce a biologically motivated algorithm for the evolutionary dynamics that combines elements of the Metropolis-Hastings algorithm with Kimura's classical population genetics theory. This approach implements a two-step process that separately models:

1. The probability of mutation occurring (mutation accessibility) as governed by the genotype-phenotype density.
2. The probability of fixation within a population (selection) determined by the fitness effect of such mutation and the effective population size.

##### K.1.1. Biological Motivation

In natural populations, evolution proceeds through two distinct processes:

1. **Mutation**: New variants arise through random genetic changes. The probability of specific mutations depends on molecular mechanisms and constraints of the genotype-phenotype map.
2. **Fixation**: Once a mutation occurs, it must spread through the population to become fixed. The probability of fixation depends on the selection coefficient (fitness advantage) and the effective population size.

##### K.1.2. Mathematical Framework

###### K.1.2.1. Mutation Probability

The probability of a mutation from phenotype $\underline{x}$ to $\underline{x}'$ is modeled using a Metropolis-like criterion based on the genotype-phenotype density:

$$\pi_{\text{mut}}(\underline{x} \to \underline{x}') = \min\left(1, \left(\frac{GP(\underline{x}')}{GP(\underline{x})}\right)^{\beta}\right) \tag{S126}$$

Here, $GP(\underline{x})$ represents the genotype-phenotype density at phenotype $\underline{x}$, and $\beta$ is a parameter controlling the strength of mutational constraints. The higher $\beta$ is, the less likely a mutation going downhill in genotype-phenotype density is to be accepted.

###### K.1.2.2. Kimura's Fixation Probability

Once a mutation occurs, its probability of fixation in a population of effective size $N$ is given by Kimura's formula

$$\pi_{\text{fix}}(\underline{x} \to \underline{x}') = \frac{1 - e^{-2s}}{1 - e^{-2Ns}} \tag{S127}$$

where $s$ is the selection coefficient. We compute this selection coefficient as the difference in fitness between the new and old phenotype divided by the fitness of the old phenotype:

$$s = \frac{F_E(\underline{x}') - F_E(\underline{x})}{F_E(\underline{x})} \tag{S128}$$

This equation captures a fundamental result from population genetics: beneficial mutations ($s > 0$) have a higher probability of fixation, with the probability approaching $2s$ for small positive selection coefficients and approaching 1 for large positive selection coefficients. Deleterious mutations ($s < 0$) have an exponentially decreasing probability of fixation as population size increases.

##### K.1.3. Overall Acceptance Probability

The overall acceptance probability—defined as the probability of a proposed step $\underline{x} \to \underline{x}'$—is the product of the mutation probability and the fixation probability:

$$\pi_{\text{accept}}(\underline{x} \to \underline{x}') = \pi_{\text{mut}}(\underline{x} \to \underline{x}') \cdot \pi_{\text{fix}}(\underline{x} \to \underline{x}') \tag{S129}$$

This two-step process better reflects the biological reality of evolution, where both mutational accessibility and selection contribute to evolutionary trajectories.

##### K.2. Effect of Population Size

Figure N shows the effect of population size parameter $N$ influences evolutionary trajectories in our Metropolis-Kimura framework. Through a series of panels showing both phenotypic trajectories (upper) and fitness dynamics (lower), we can observe how different population sizes affect the balance between exploration and exploitation in phenotypic space.

At small population sizes (left panels), the evolutionary trajectories exhibit highly stochastic behavior, with populations taking meandering paths through the phenotypic landscape. This represents a regime where genetic drift plays a dominant role. As population size increases (moving right), the trajectories become increasingly deterministic, with populations taking more direct paths toward fitness peaks, while avoiding genotype-to-phenotype density valleys. This transition reflects a shift from drift-dominated to selection-dominated dynamics, where populations more strictly follow the local fitness gradient. The fitness trajectories in the lower panels further reinforce this pattern, showing more erratic fitness fluctuations at small population sizes and more monotonic increases at large population sizes, consistent with stronger selective pressures.

##### K.3. Main Text Results

Having developed this alternative algorithm for the population dynamics, we reproduce the results from the main text using this new algorithm. In particular, Figure 2, 4, and 5 in the main text used synthetic data generated by the algorithm described in Section B. We now show that the same results can be obtained using the Metropolis-Kimura algorithm.

###### K.3.1. Figure 2 Main Text

As shown in Figure O, the evolutionary trajectories generated with the Metropolis-Kimura algorithm are qualitatively identical to those in the main text. The lineages evolve towards regions of higher fitness while respecting the constraints imposed by the genotype-phenotype density, confirming that our main conclusions are robust to the specific choice of the evolutionary simulation algorithm.

###### K.3.2. Figure 4 Main Text

Similarly, Figure P demonstrates that when we analyze the synthetic data from the Metropolis-Kimura simulation, our main findings regarding the latent space hold. The RHVAE continues to outperform both the standard VAE and PCA in preserving the geometric structure of the original phenotypic space and achieves superior reconstruction accuracy. This confirms that the advantages of the geometry-informed approach are not an artifact of the simulation method.

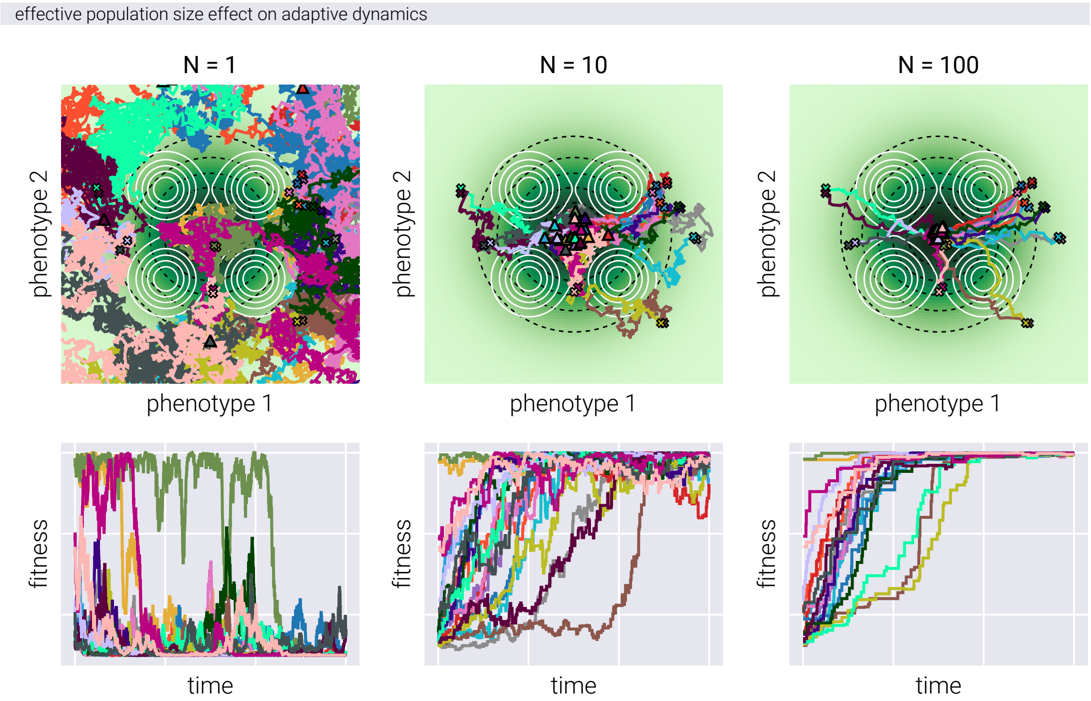

**Figure N. Effect of population size on adaptive dynamics**. Upper panels: Population trajectories on a landscape with a single fitness peak and four genotype-phenotype density peaks. Black dashed lines indicate fitness landscape contours, while white solid lines show genotype-phenotype density contours. Colored lines represent independent population trajectories, with initial positions (crosses) and final positions (triangles). Lower panels: Temporal evolution of population fitness. Panels from left to right show increasing values of population size ($N$), demonstrating increasingly deterministic adaptive trajectories.

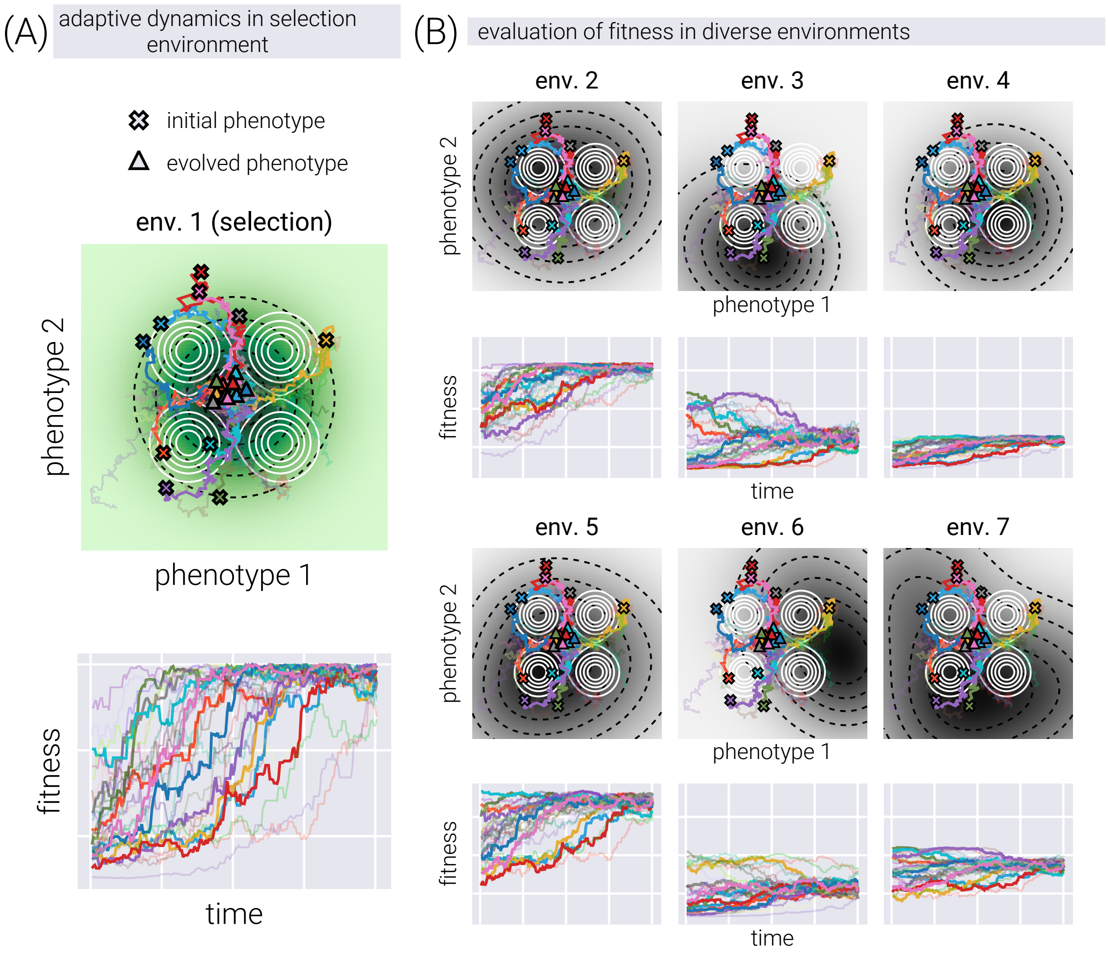

**Figure O. Evolutionary dynamics on phenotype space with Metropolis-Kimura dynamics.** (A) Top: Metropolis-like evolutionary dynamics on phenotype space. Each line represents the trajectory of a lineage as it evolves over time, with crosses and triangles denoting the initial phenotypic coordinate of a few selected lineages. Note that trajectories tend to move towards higher fitness values, avoiding low genotype-to-phenotype density regions. Bottom: Fitness over time of the same trajectories shown above. (B) The same trajectories shown in (A) overlaid on different fitness maps determined by different environments. Although the phenotypic coordinates of the genotypes remain the same (top panels), the resulting fitness readouts change as the topography of the environment-dependent fitness landscape changes (bottom panels).

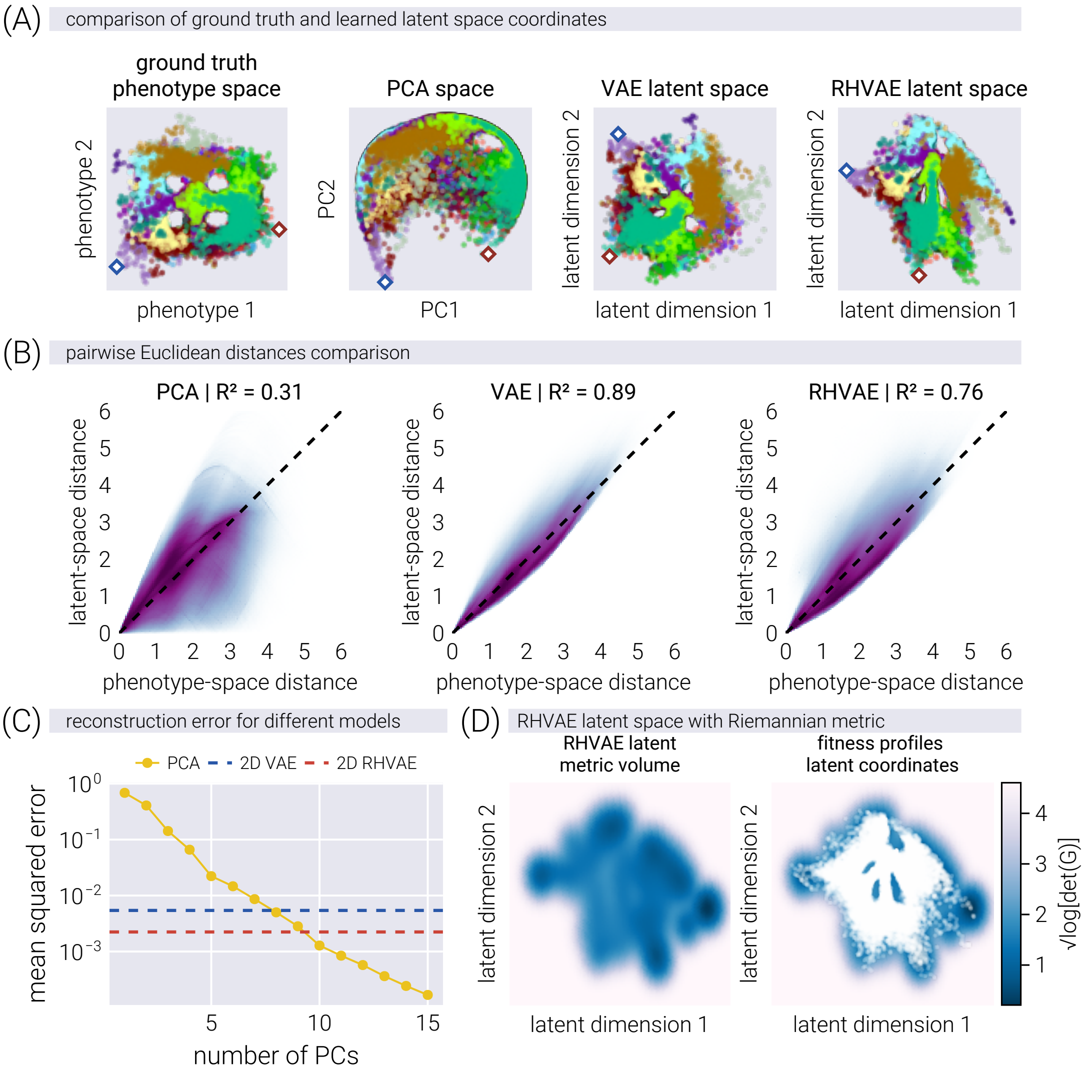

**Figure P. Geometry-informed latent space captures phenotypic structure and improves reconstruction accuracy**. (A) Latent space coordinates of simulated fitness profiles colored by lineage, after alignment with the ground truth phenotype space using Procrustes analysis. The RHVAE better preserves phenotypic relationships than PCA or vanilla VAE, as evidenced by the highlighted points (diamond markers). (B) Comparison of all pairwise Euclidean distances between genotypes in the ground truth phenotypic space versus the different latent spaces, showing RHVAE better preserves the original distances. Given the large number of data points and the even larger number of pairwise comparisons, the data are shown as a smear rather than single points. (C) Reconstruction error (MSE) comparison showing 2D-RHVAE achieves accuracy comparable to a 10-dimensional PCA model. (D) Metric volume visualization of the RHVAE latent space (left) and with projected data points (right), revealing regions of high curvature that correspond to areas of low genotype-phenotype density in the simulation.

##### K.3.3. Figure 5 Main Text

Finally, Figure Q shows the fitness landscape reconstructions from the data generated with the Metropolis-Kimura dynamics. Consistent with the results in the main text, the RHVAE model successfully reconstructs the complex, nonlinear topography of the landscapes across different environments. This result further strengthens our conclusion that the RHVAE framework is a powerful tool for inferring fitness landscapes from high-dimensional data, regardless of the precise evolutionary dynamics used to generate it.

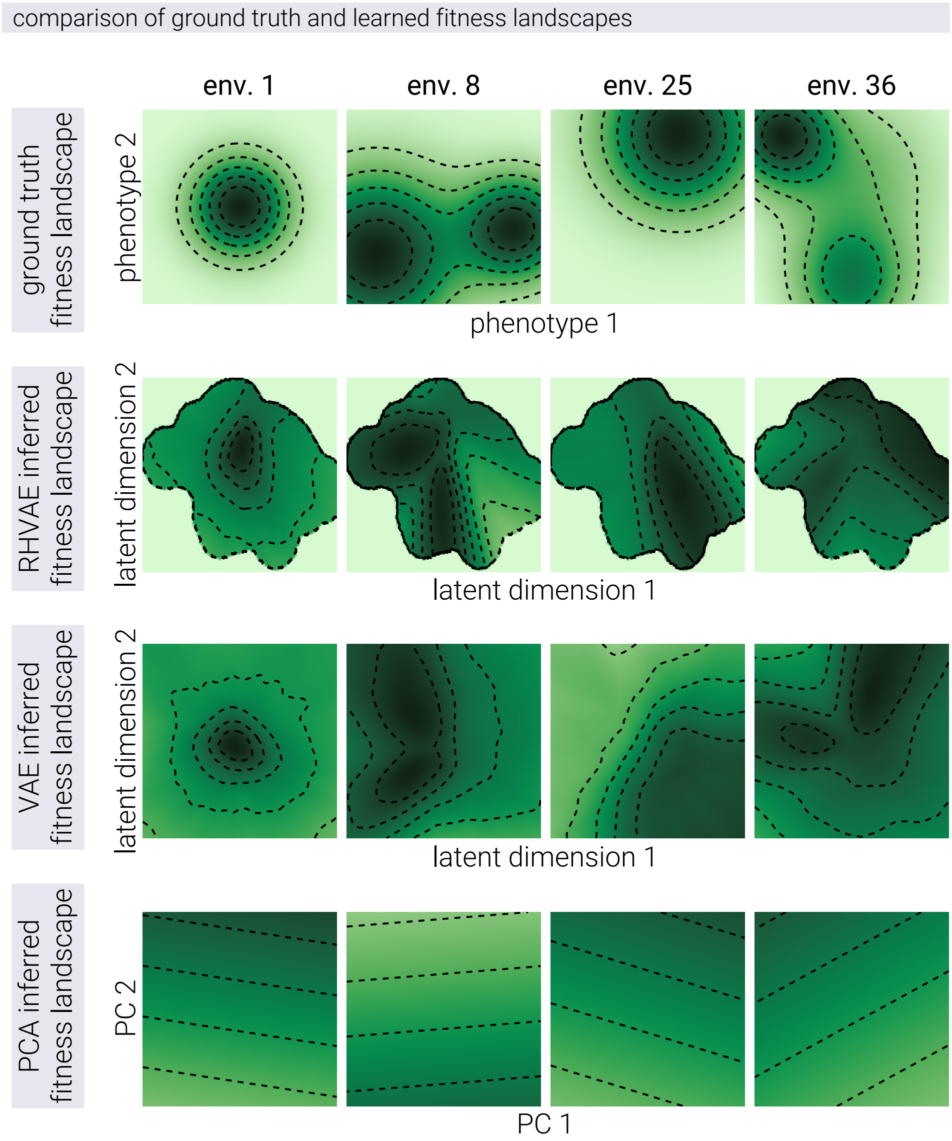

**Figure Q. Geometry-informed latent space enables accurate reconstruction of complex fitness landscapes**. Comparison of ground truth and reconstructed fitness landscapes across multiple environments. The first row shows examples of the original simulated fitness landscapes used to generate the data. Subsequent rows show reconstructions using the RHVAE (second row), VAE (third row), and PCA (fourth row) models. The RHVAE accurately captures the underlying topography, including the number and relative positions of fitness peaks. While the VAE captures general landscape shapes, it lacks a geometric metric to define prediction reliability regions. PCA fails to represent the nonlinear structure of the landscapes, demonstrating the limitations of linear dimensionality reduction for complex fitness landscape reconstruction.
